# Modeling and Analysis of the Macronutrient Signaling Network in Budding Yeast

**DOI:** 10.1101/2020.02.15.950881

**Authors:** Amogh P. Jalihal, Pavel Kraikivski, T. M. Murali, John J. Tyson

## Abstract

In eukaryotes, distinct nutrient signals are relayed by specific plasma membrane receptors to signal transduction pathways that are interconnected in complex information-processing networks. The function of these networks is to govern robust cellular responses to unpredictable changes in the nutritional environment of the cell. In the budding yeast, *Saccharomyces cerevisiae*, these nutrient signaling pathways and their interconnections have been well characterized. However the complexity of the signaling network confounds the interpretation of the overall regulatory ‘logic’ of the control system. Here, we propose a literature-curated molecular mechanism of the integrated nutrient signaling network in budding yeast, focusing on early temporal responses to carbon and nitrogen signaling. We build a computational model of this network to reconcile literature-curated quantitative experimental data with our proposed molecular mechanism. We evaluate the robustness of our estimates of the model’s kinetic parameter values. We test the model by comparing predictions made in mutant strains with qualitative experimental observations made in the same strains. Finally, we use the model to predict nutrient-responsive transcription factor activities in a number of mutant strains undergoing complex nutrient shifts.

## Introduction

In yeast and other eukaryotes, the sensing and transmission of distinct classes of macronutrient signals are carried out by specialized signaling pathways [1]. These environmental signals, which may be synergistic or antagonistic, are processed within the cell by crosstalk among the signal-transduction pathways in order to produce an adaptive and robust response to the nutritional environment of the cell. This response ultimately manifests itself physiologically in a modulation of the growth rate of the cell, by regulation of global transcriptional and translational activities [2, 3]. Adaptive modulation of the global cellular growth state is crucial for the survival of unicellular organisms that are exposed to fluctuating environments.

Budding yeast *Saccharomyces cerevisiae* is an ideal eukaryotic organism in which to study nutrient signaling for three reasons: it is a unicellular ‘model’ organism whose metabolism has been thoroughly characterized from many angles, its genetic constitution is well known and easily manipulated by standard techniques, and there is a large degree of molecular homology between yeast and mammalian nutrient signaling systems [4, 5]. Despite this high degree of homology, the nutrient signaling system plays divergent roles in yeast and in mammals. In buding yeast cells, the default state is exponential growth, and nutrient starvation signals inhibit cell growth. In mammalian cells, on the other hand, the default state is the inhibition of growth, despite abundant and continuous availability of nutrients in the blood stream. Cell growth and division is stimulated by hormonal signals that trigger growth pathways in competent cells. Dysregulation of the default state in mammalian cells leads to a (yeast-like) growth and proliferation state that is characteristic of cancer cells.

In yeast, distinct pathways sense and signal the sufficiency of the two major classes of macronutrients, carbon and nitrogen. The Ras/cAMP/PKA pathway largely conveys the glucose status of the environment, suppressing carbon-adaptation and general stress responses when glucose is plentiful, while up-regulating ribosome biogenesis and overall growth rate [6, 7, 8, 9]. When glucose is in short supply, the Snf1 pathway shuts off catabolite repression and up-regulates carbon adaptation responses [10, 11]. In the presence of rich nitrogen sources, the TORC1 pathway activates nitrogen catabolite repression (NCR) and up-regulates ribosome biogenesis via the Sch9-kinase branch [12, 13, 14, 15]. In response to declining nitrogen status, the Tap42-Sit4/PP2A phosphatase branch activates nitrogen adaptation responses [16].

While the upstream sensing mechanisms are nutrient specific, the downstream signal-processing network is characterized by crosstalk among the signaling pathways just described. Consequently, how the cell responds to specific combinations of nutrients depends on complex interactions among key regulatory enzymes in the network [17, 18, 19, 20, 21]. In yeast, the pathways responsive to particular nutrients have been well studied, but the molecular decision-making carried out by the network *as a whole* remains unclear. The complexity of the system obfuscates the interpretation of experimental observations, based on genetic and environmental perturbations that interrogate the nutrient adaptation responses [22].

In the study of such complex biochemical control systems, dynamical modeling has proven to be useful in determining cellular behaviors that result from entangled molecular interactions 23, 24, 25]. These dynamical models are often presented as systems of nonlinear ordinary differential equations (ODEs) built from biochemical rate laws describing the reaction steps that constitute the biochemical interaction network. Previous authors have published ODE models (see Box 1) of nutrient signaling in yeast cells in restricted physiological contexts. In this paper, we construct a model of broader scope based on a literature-curated diagram of carbon and nitrogen signaling, focusing on the short-term response of yeast cells to nutrient stresses. The nonlinear ODE model based on this diagram is calibrated with time-series measurements of key regulatory proteins as well as perturbation experiments carried out on wild-type and mutant strains. Finally, we use the calibrated model to predict nutrient responses of yeast cells under novel conditions.

## Results

### A proposal for the nutrient signaling network in budding yeast

In order to construct a coherent and comprehensive regulatory network that is consistent with a broad range of experimental results, we first gathered relevant data on nutrient signaling from the published literature along with hypotheses about the underlying molecular mechanisms. On the basis of this information we propose a molecular regulatory network for nutrient signaling in budding yeast in Figure 1.

**Figure 1:**
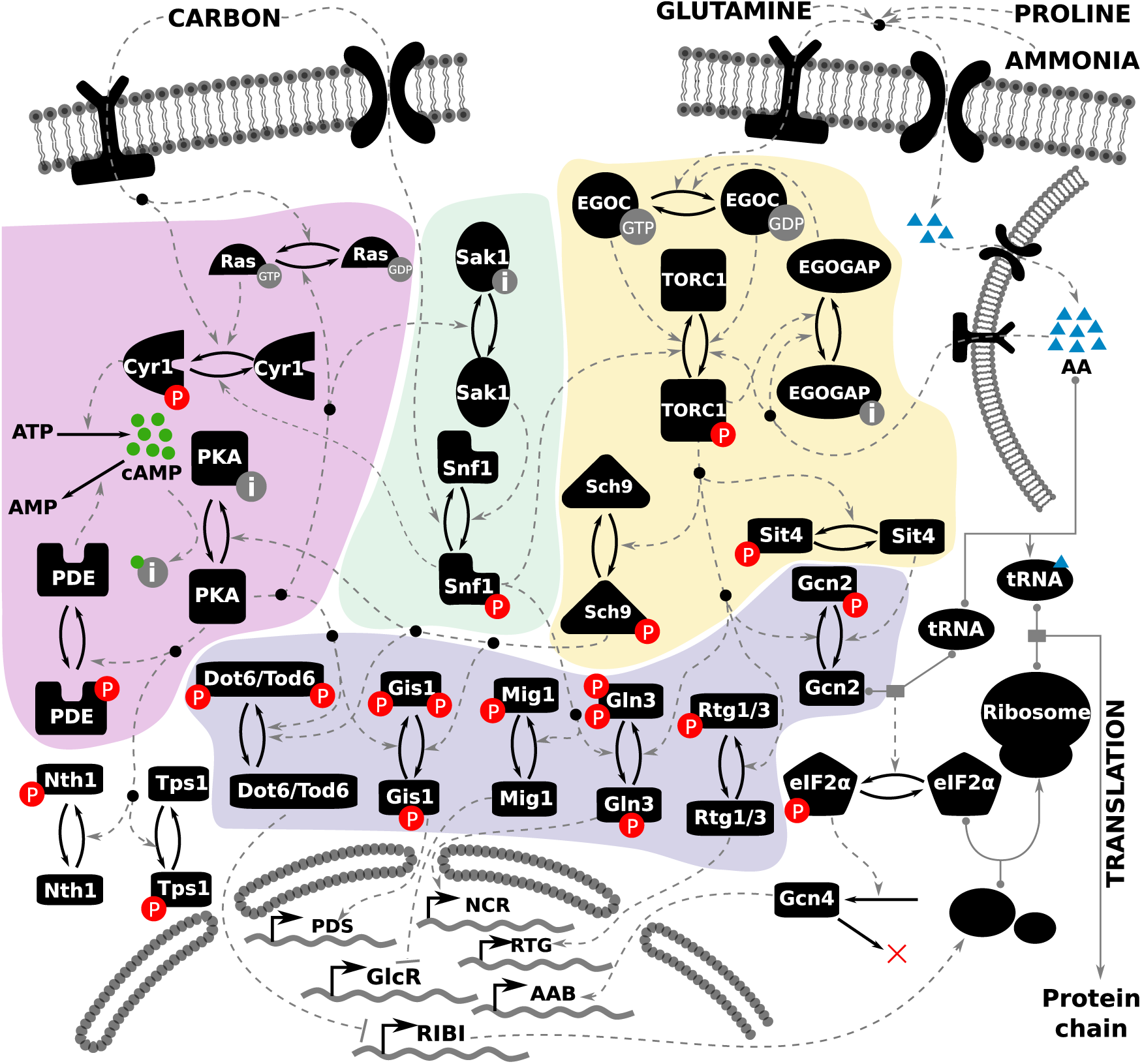
Literature-curated molecular regulatory network of nutrient signaling in yeast cells. cAMP-PKA pathway is depicted on the left (highlighted in purple), the Snf1 pathway in the middle (highlighted in green) and the TORC1 pathway on the right (highlighted in yellow). The readouts of the model are the activities of the transcription factors towards the bottom of the diagram (highlighted in blue), whose target genes at the bottom of the figure comprise various regulons: GlcR, glucose-repressed genes; PDS, post-diauxic shift element; NCR, nitrogen catabolite repression genes; RTG, retrograde genes; AAB, amino acid biosynthesis; and RIBI, ribosome biogenesis. Each black icon represents a molecular species. Post-translational modifications are represented by a pair of solid arrows pointing in opposite directions. Phosphorylation is indicated by ‘P’ in a red circle. The inactive form of a species is indicated by the letter ‘i’ in a gray circle. Guanidylation is indicated by ‘GTP’ in a grey circle. Regulatory signals are represented by dashed gray lines. Complex formation is indicated by a solid gray line, with the binding partners indicated by gray circles. The inputs to the model are shown at the top of the figure, represented by one carbon input, and three nitrogen inputs: glutamine, ammonia and proline. The intracellular amino acid pool stored in the vacuole is represented by the membrane-bound structure on the top right.

In yeast, a set of nutrient-specific membrane-bound receptors sense and transduce signals to the down-stream signaling pathways within the cell [1]. In our model, the nutrient receptors are abstracted away; the notion of nutrient sufficiency of carbon and nitrogen is represented on a scale from 0 to 1, where 0 represents starvation and 1 represents abundance. There are four such nutrient inputs for ‘Carbon’ (i.e., glucose) and three classes of nitrogen: ‘Glutamine’ (high quality), ‘Proline’ (poor quality), and ‘Ammonia’ (which requires a carbon source). The output of the network is the regulation of transcription factors governing global metabolic transitions (shown at the bottom of the figure). Since the activities of these transcriptional regulators will determine the growth status of the cell, they are used as readouts of the cellular state in response to a nutrient input. Table 1 summarizes the interacting components of the model, their regulators, and their targets.

**Table 1:** The molecular interactions in our proposed nutrient signaling model. For each model variable, the corresponding upstream regulators and downstream targets are listed.

| Variables | Regulators | Targets |
| --- | --- | --- |
| Carbon | model input | Cyr1, Ras |
| Glutamine<sub>ext</sub> | model input | Glutamine |
| NH<sub>4</sub> | model input | Glutamine |
| Proline | model input | Glutamine |
| Glutamine | Glutamine<sub>ext</sub>, NH<sub>4</sub>, Proline | TORC1, EGO C |
| Ras | Glucose [47], PKA [48] | Cyr1 |
| Cyr1 | Ras, Glucose [7] | cAMP |
| cAMP | Cyr1, PDE [49] | PKA |
| PKA | cAMP [50], Sch9 [51] | Gis1, Dot6, Sak1,<br>PDE, Mig1, Tps1,<br>Trehalase, Ras |
| PDE | PKA [50] | cAMP |
| EGOC | Glutamine, N sources, EGO GAP [30] | TORC1 |
| EGOGAP | Glutamine, TORC1 [52] | EGOC |
| TORC1 | Glutamine, EGOC, EGO GAP, Snf1 [22] | Sch9, Gln3, Rtg1/3 |
| Sch9 | TORC1 [53] | PKA, Dot6, Gis1, Gcn2 |
| Sak1 | PKA [54] | Snf1 |
| Snf1 | Sak1, Carbon [55] | Mig1, Gln3, Cyr1 |
| Gis1 | PKA, Sch9 [56] | Post-Diauxic Shift genes |
| Mig1 | Snf1 [40], PKA (via Reg1/Glc7) [57] | Fermentation |
| Gln3 | Snf1, TORC1 [58] | NCR genes |
| Rtg1/3 | TORC1 [59] | Retrograde genes |
| Gcn2 | Sch9 [60], Snf1 [10] | Gcn4 |
| Gcn4 | tRNA, Gcn2, Glutamine [61] | Amino Acid Biosynthesis genes |
| Dot6 | Sch9, PKA [62] | RIBI |
| Tps1 | PKA [63] | model output |
| Nth1 | PKA [63] | model output |
| Rib | Dot6 [64] | model output |

In view of the large number of interacting components and the relative paucity of high-resolution data on their dynamics, we are compelled to simplify the model to make the problem more manageable without—we hope—sacrificing its most important details. In the following sections, we describe the key simplifications we have made.

#### Storage molecules

In yeast, amino acids are stored both in the cytosol and in a vacuole [26, 27, 28, 29]. The regulation of these distinct, dynamic nutrient pools is complex. Since glutamine has been shown to be the primary nutrient input to TORC1 30], a single variable representing intracellular glutamine level, depicted as AA in Figure 1, is used to quantify intracellular nitrogen sufficiency. In the absence of external glutamine, but in the presence of sufficient carbon sources, yeast cells can synthesize glutamine from ammonia. The uptake of ammonia is regulated by ammonia transporters that are expressed in response to declining glutamine levels [31]. Finally in the presence of poor nitrogen sources like proline, yeast cells activate amino acid biosynthesis pathways. Nitrogen stores are depicted on the top right of Figure 1, where the lipid bilayer depicts the yeast vacuole membrane.

###### The cAMP-PKA pathway

Using a nonlinear ODE model, Garmendia-Torres *et al*., 2007 investigated the PKA-mediated regulation of Msn2/4 nucleocytoplasmic oscillations, capturing the essential feedback loops in the PKA activation mechanism [32]. Gonze *et al*., 2008 proposed a stochastic version of this model [33]. Cazzaniga *et al*., 2008 investigated the guanine nucleotide exchange reactions in the PKA activation mechanism but failed to capture the short time-scale behavior of cAMP dynamics [34, 35]. Stewart-Ornstein *et al*., 2017 presented a simplified mechanism to investigate, both computationally and experimentally, the upstream mechanism of cAMP activation 36]. Williamson *et al*., 2009 created a full model of the PKA pathway, incorporating substantial mechanistic details with respect to alternative PKA activation mechanisms, with a focus on understanding features of cAMP regulation [37]. In order to account for experimental observations not captured by these models, Gonzales *et al*., 2013 used a simplified mechanism to investigate strain-specific cAMP oscillatory dynamics 38]. All the models described in this paragraph have focused on the mechanism of cAMP regulation, or the regulation of Msn2/4 nucleocytoplasmic oscillations purely in response to glucose abundance or starvation.

###### The Snf1 pathway

Garcia-Salcedo *et al*., 2014 systematically explored a series of hypothetical interactions in the Snf1 pathway in order to identify a high confidence network topology that recapitulated experimentally observed phenotypes [39]. More recently Welkenhuysen *et al*., 2017 [40] have investigated the structure of the Snf1 pathway in the context of glucose starvation.

###### The TORC1 pathway

Many mathematical models of mTOR signaling have been proposed. Of particular relevance to our work, Vinod *et al*. 2009 investigated the role of mTOR in metabolic regulations in response to insulin signaling [41]. Sonntag *et al*. 2012 carried out a systematic investigation of regulatory mechanisms underlying the interactions between AMPK and mTORC2 [42]. Pezze *et al*. 2016 expanded this model to study its response to amino acids [43]. Thobe *et al*. 2017 studied the regulation of mTORC2 using logical modeling [44].

###### Integrated pathway models

Sengupta *et al*., 2007 studied the steady-state properties of an integrated model of the cAMP-PKA pathway and the MAPK signaling system in regulating the pseudohyphal transition of yeast cells to response to nitrogen starvation [45]. Recently Welkenhuysen *et al*., 2019 created a Boolean model of the crosstalk between various signaling pathways [46], to show how the network confers robustness to diverse nutrient environments.

In order to achieve maximal growth rate, the cell requires a sufficient amount of ATP. Instead of modeling the dynamic regulation of cellular energy charge (EC), we assume that EC is proportional to the carbon sufficiency of the environment, while acknowledging that the true relationship is highly non-linear [65]. Finally, in response to carbon starvation, the activity of trehalose synthase (Tps1) is upregulated in order to synthesize the storage molecule trehalose, and upon relief from starvation, the PKA pathway activates trehalase (Nth1) in order to consume the stored trehalose [63]. The activities of Tps1 and Nth1 are used as readouts to characterize the yeast cell’s response to carbon shifts. We do not model the changing levels of trehalose or glycogen because they are not directly relevant to the short term responses of the carbon signaling pathway.

#### Nitrogen sensing

See the area of Figure 1 that is highlighted in yellow. The precise mechanism of nitrogen sensing in yeast by TORC1 remains controversial. The TOR complex has been shown to interact at the vacuolar membrane with multiple G-proteins, which sense the levels of distinct intracellular amino acid pools [52, 1]. The EGO complex, comprised of the G-proteins Gtr1/2 and their membrane anchors Ego1/2/3, is activated by various subunit-specific GTPase activating proteins (GAPs). For example Lst4/7 has been shown to be a GAP specific to Gtr2 [66], while the SEACIT complex is the Gtr1-GAP [67]. We simplify this mechanism by letting one variable (EGOC) represent the activity of the EGO complex and another (EGOGAP) represent the activity of a generic GAP protein.

#### PKA activation

See the area of Figure 1 that is highlighted in pink. Several published models have explored the complexity of Protein Kinase A activation via the Ras-cAMP mechanism with a focus on cAMP dynamics (Box 1). In our version of this pathway, we include reactions for cAMP sythesis (by adenylate cyclase, Cyr1) and degradation (by phosphodiesterase, PDE) and shift our attention to the interactions between the PKA pathway and the rest of the nutrient signaling system. Glucose has been shown to stimulate Cyr1 both directly [68, 69] and indirectly, via Ras2 [47]. cAMP activates the PKA heterotetramer, here represented by a single species ‘PKA’. PKA activates PDE, which converts cAMP to AMP. In our model, ‘PDE’ is a single variable representing phosphodiesterases Pde1/2/3.

#### The Snf1 pathway

See the area of Figure 1 that is highlighted in green. Snf1 is inhibited by the glucose-dependent phosphatase Reg1/Glc7; however, the exact mechanism remains unclear. Hence, in our regulatory network we simply assume a carbon source-dependent inhibition of Snf1 activity. During glucose starvation, Snf1 is activated by Sak1/Tos3/Elm1 kinases 39], which are represented by a single variable ‘Sak1’ in our model.

#### Crosstalk between pathways

PKA and Snf1 have been shown to mutually inhibit each other: Snf1 inhibits Cyr1 activity thereby inhibiting PKA; while PKA inhibits Sak1 kinase, thereby inhibiting Snf1 [54, 70]. While Castermans *et al*. have speculated that glucose-activated phosphatases might be regulated by the PKA pathway [57], Nicastro *et al*. propose that PKA might inhibit Sak1 kinase activity [54]. Here, we assume PKA-dependent Snf1 dephosphorylation occurs via the inhibition of Sak1/Tos3/Elm1 activity. The relationship between Snf1 and TORC1 is less clear. While glucose-dependent TORC1/Sch9 kinase activity has been reported in multiple publications, the mechanism by which TORC1 activity is modulated by the presence of glucose remains unclear [53]. Based on data from Hughes-Hallett *et al*. and Prouteau *et al*., we assume that Snf1 inhibits TORC1 [71, 72]. Finally, the relationship between Sch9 and PKA is confounded by the complex and overlapping regulation of their many shared downstream targets [19]. While early studies on the PKA pathway found an apparent negative interaction between Sch9 and PKA [73], the exact relationship between these kinases remains unclear. Zhang *et al*. claim that Sch9 interacts with and inhibits PKA at multiple levels [51, 74]. Here, we assume a direct inhibition of PKA by Sch9.

### Creating a mathematical model

The component parts of the network (discussed above) are derived from experimental evidences in the literature. When we assemble these components into a comprehensive signaling network (Figure 1), it is not clear that the full network is still consistent with all the individual experiments. To answer this question, we converted our proposed regulatory mechanism into a system of ODEs using the Standard Component Modeling framework (‘Methods’, Section S1). The full set of 25 ODEs, along with details of the representation of certain abstract variables (like the intracellular nutrient levels) are presented in Section S1.1. The ODEs contain 128 parameters (Table S2). Before we can simulate the behavior of the model equations, we must estimate the values of these parameters. To make these estimates, we need some experimental data on the time-courses of molecular variables in the model and some observations of the steady-state characteristics of the signaling network under a variety of conditions. In the next section, we describe the experimental datasets we collected and how we estimate the parameter values, and in later sections we use the parametrized model to make predictions about nutritional responses of yeast cells under novel conditions.

### Parametrizing the model with experimental data

In the following two subsections, we describe the two types of quantitative data used to calibrate the model: (1) time-course measurements and (2) steady-state nutrient-shift data, related to the abundance/activity of particular molecules in the nutrient signaling pathway in both wild-type (*wt*) and mutant backgrounds. From this data we derived a set of parameter values that provides a ‘best’ fit to the experimental observations, using parameter-estimation tools that minimize the sum of squared differences between experimental measurements and model simulations (see ‘Methods’ section). In the third subsection, we find alternative sets of parameter values that fit the quantitative data nearly as well as the best-fitting, ‘optimal’ parameter set.

#### Comparing simulation results with experimental time-course data

First, we combed the literature for experimental time-course measurements of molecular species in our model (see Supplementary Table 2). In Figure 2 (a)-(j), we compare the experimental data (open circles) to time-series simulations (in red) of the model using the optimal parameter values.

**Figure 2:**
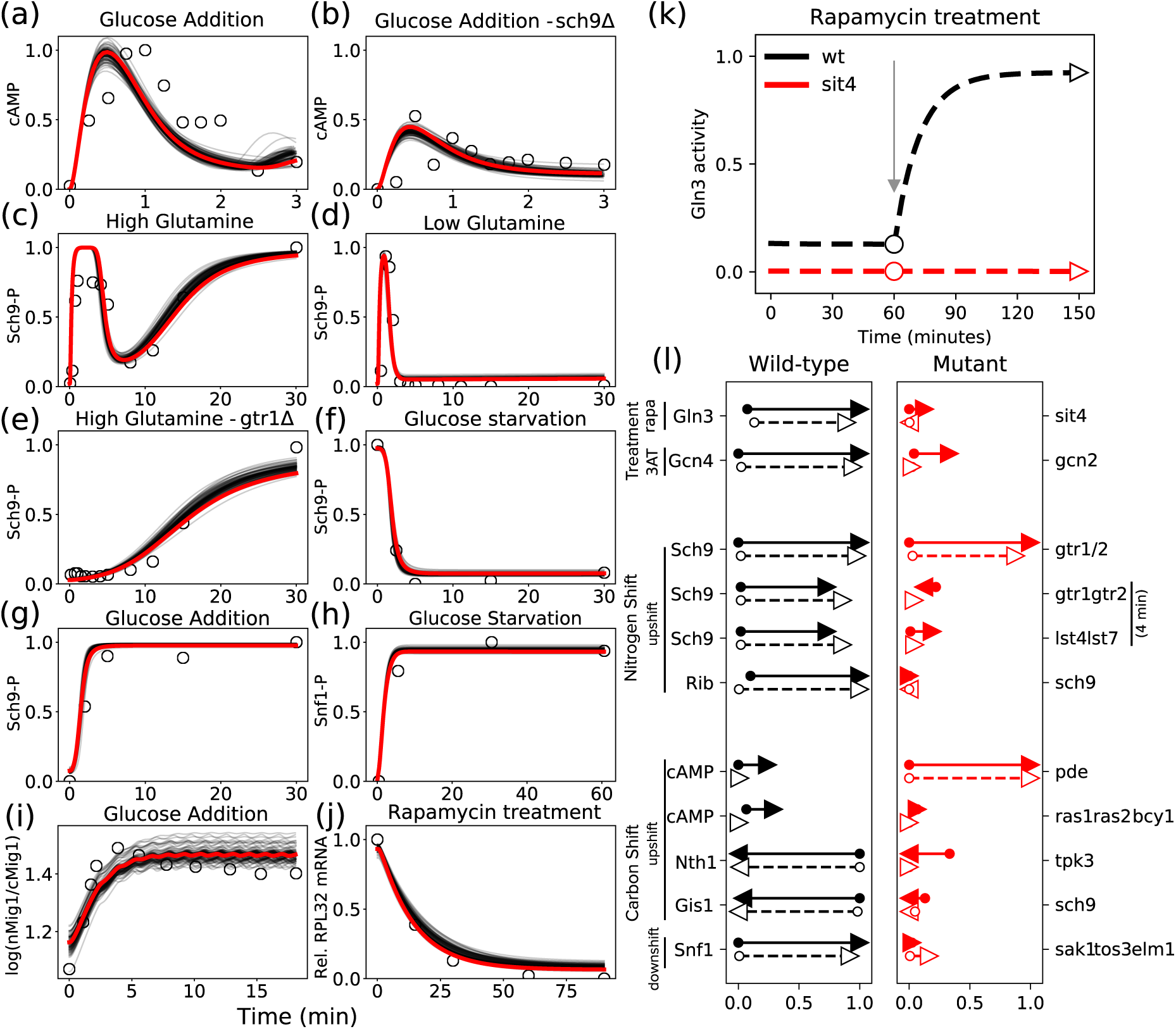
The model successfully explains experimental data. (a)-(j) Time series fits to literature-curated time-course data. The red lines represent simulated trajectories from the best-fit parameter set, while the gray lines represent simulations from 100 parameter sets with comparable sums of squared errors. The experimental measurements are shown as open circles. (a) and (b): relief from glucose starvation. (c), (d) and (e): relief from nitrogen starvation by glutamine addition. (f) and (h): glucose starvation. (g) and (i): relief from glucose starvation. (j) Rapamycin treatment of well-nourished cells. (k) Comparing model simulations with data from a shift experiment [76] measuring Gln3 phosphorylation in response to rapamycin treatment in well-nourished *wt* and *sit4*Δ strains. In the *wt* simulation (black dashed line), we calculate the steady state value of Gln3 in rich medium in the absence of rapamycin (black circle). Next, rapamycin is introduced (at the gray arrow), and the new, postshift steady state is recorded (black triangle). The same simulation was repeated for *sit4*Δ cells to yield preshift (red circle) and postshift (red triangle) steady states. (l) Visualization of perturbation analyses. The perturbations are drug treatments or nutrient shifts carried out in *wt* cells (left column, black arrows) and mutant strains (right column, red arrows). The labels on the left indicate the molecular species being assayed, and the labels on the far right indicate the gene(s) deleted in the mutant strains. Experimental results are represented by solid arrows, and model simulations by dashed arrows. For each molecular species under consideration, the arrow’s tail and head indicate (respectively) the pre- and post-perturbation steady-state values, as measured on the relative scale (0 to 1) at the bottom of the column. As an example, the rapamycin treatment in panel (k) is reproduced in the first line of panel (l). Note that the *y*-axis in panel (k) is now the *x*-axis in panel (l).

Our model includes molecular species that are common readouts in experiments investigating nutrient signaling. cAMP is a standard readout for the activity of the PKA pathway. cAMP shows rapid transient changes, on the time scale of minutes, upon relief from glucose starvation. Model simulations successfully capture the cAMP dynamics in *wt* cells as well as *sch9*Δ mutant cells (Figure 2(a), (b)). Sch9 phosphorylation is a widely accepted readout for TORC1 activity and displays changes on the time scale of tens of minutes upon relief from nitrogen starvation [30]. Model simulations successfully capture the transient as well as steady state behavior of Sch9 phosphorylation in *wt* and in *gtr1,2*Δ strains (Figure 2 (c), (d), and (e)). Moreover, we are able to capture changes in Sch9 phosphorylation in response to glucose starvation and readdition (Figure 2(f), (g)), as reported by Prouteau *et al*. [72]. Snf1 kinase activity is typically measured indirectly, by examining the phosphorylation of the SAMS peptide, a Snf1 target sequence [75]. The model successfully captures the activation of Snf1 during glucose downshift (Figure 2(h)) leading to the expression of glucose-repressed genes. Apart from assays of biochemical activities, subcellular localization measured by fluorescence microscopy provides indirect information about the activity of transcription factors. Mig1 nuclear localization upon relief from glucose starvation was recently reported by Welkenhuysen *et al*. [46] (Figure 2(i)). Model simulations successfully capture this response, if we identify Mig1 transcriptional activity with the experimentally observed translocation of Mig1 into the nucleus. Direct information about transcription factor activity can also be obtained by measuring mRNA levels of the transcription factor’s target genes. For example, *RPL*32 mRNA, encoding the large ribosomal subunit, is down-regulated in response to rapamycin treatment, and our model captures this effect nicely (Figure 2(j)).

#### Capturing phenotypes of *wt* and mutant strains in response to nutrient shifts

To investigate the roles of regulators in nutrient signaling, experimentalists typically characterize the phenotype of a strain in response to perturbations (nutrient shifts or drug treatments). In order to parametrize our model we collected data from direct biochemical assays of molecular responses to perturbations (see Supplementary Table 3).

An example of the type of perturbation data used to calibrate our model is illustrated in Figure 2(k), where model simulations are compared to data reported in Beck *et al*., 1999 [76], who studied the phosphorylation of Gln3 by TORC1 and Sit4. Using Western blots of Gln3 pull-downs, Beck *et al*. found that Gln3 is phosphorylated (inactive) in untreated *wt* cells (black dashed line), while treatment with rapamycin led to Gln3 dephosphorylation. In contrast, Gln3 remained phosphorylated (inactive) in a *sit4*Δ strain even after rapamycin treatment (red dashed line). The figure shows simulated time courses of active (dephosphorylated) Gln3 under these conditions, and the markers (empty circles and empty triangles) indicate the (pre- and post-shift) steady-state values of active Gln3 used to compare against the experimental measurements.

Figure 2(l) visualizes such comparisons for a representative sample of experimental perturbations. For example, the results in Figure 2(k) are summarized in the first line of Figure 2(l). The second line shows the results of 3-amino-1,2,4-triazole (3AT) treatment on Gcn4 activity in *wt* cells (black) and in a *gcn*2 deletion strain (red). 3AT inhibits histidine synthesis, resulting in induction of Gcn4 and an amino acid starvation response. There is no induction of Gcn4 expression in a *gcn2*Δ strain.

The second and third blocks of Figure 2(l) show the results of ‘nitrogen shifts’ and ‘carbon shifts’; either a downshift (nutrient starvation) or upshift (relief from starvation). The first two rows in the ‘Nitrogen Shift’ block (call them N1 and N2) repeat the results in panels (c) and (e), which report the phosphorylation of Sch9 in *wt* and *gtr1*Δ*gtr2*Δ strains in response to relief from starvation (i.e., addition of glutamine to nitrogen-starved cells). Row N1 compares the pre-shift steady state (at time 0) to the post-shift steady state (at time 30 min). Row N2 compares the pre-shift steady state to the transient state (at time 4 min). Row N3 makes a similar comparison of *wt* cells to *lst4*Δ*lst7*Δ cells. Row N4 compares expression levels of RPL25 (a ribosomal protein), as measured by [73] in *wt* and *sch9*Δ strains, with model simulations.

In the ‘Carbon Shift’ block, the first four rows report the results of readdition of glucose to carbon-starved cells, while the last row summarizes a glucose starvation experiment. Row C1, left column, repeats the results in panel (a). The *Pde1*Δ*pde2*Δ strain shows an increase in cAMP levels compared to *wt* [49], while the *ras1*Δ*ras2*Δ*bcy1*Δ strain shows a diminished amount of cAMP compared to *wt* [7]. Row C3 depicts trehalase (Nth1) levels after a glucose up-shift; trehalase levels remain low in a *tpk3*Δ mutant [8]. Row C4 demonstrates that a mutation in the nitrogen signaling pathway (*sch9*Δ) has a significant effect on the carbon stress response factor Gis1 [73]. Finally, row C5 shows that Snf1 activity increases in response to a carbon down-shift in *wt* cells, but this response is absent in *sak1*Δ*tos3*Δ*elm1*Δ cells 77].

#### Alternative sets of parameter values

In the previous subsections, quantitative data obtained from the literature was used to estimate the numerical values of the parameters in the model. Since the available data is sparse and the number of parameters is large, we expect that many parameter values will be poorly constrained by the data. Since any particular set of parameter estimates will be unreliable in this situation, we would like to obtain a representative ensemble of sets of parameter values that all give acceptable fits to the data. One approach to obtaining such an ensemble would be to randomly sample parameter values, evaluate the model for goodness-of-fit, and accept those parameter sets that meet some standard of acceptable fit. However, sampling parameter values in a completely random fashion is not an efficient way to find alternate parameter sets because small random perturbations will not effectively explore the parameter space, whereas large random perturbations are not likely to find an acceptable set of parameter values. Instead, we use the fact that the region of acceptable parameter values in a high-dimensional parameter space is characterized by ‘stiff’ and ‘sloppy’ directions [78], and a random parameter search has to respect these directions. In order to carry out this search we used a method of ‘model robustness analysis’ described by Tavassoly *et al*. [79].

The mathematical details of this method are provided in Section S2. In short, we first construct a quadratic cost function that quantifies the goodness-of-fit between model predictions and the curated data (Section S2.1). The cost is a function of the model’s parameter values. We then used a Markov-Chain Monte-Carlo (MCMC) sampling scheme to improve our initial ‘hand-tuned’ set of parameter values (Section S2.2). The resulting parameter set with the least cost we designate as the ‘optimal’ parameter set. In the neighborhood of this optimal parameter set, we expect the cost function to be bowl-shaped, such that any deviation from the optimal set will lead to an increase in cost. The curvature of the cost surface is described by the Hessian matrix of second derivatives of the cost function with respect to the parameter values. The eigenvectors of the Hessian matrix differentiate between the directions in parameter space along which the cost increases rapidly (i.e., the stiff directions) and the directions along which the cost function hardly changes (i.e., the sloppy directions). Knowledge of these directions helps us to constrain the search for alternative parameter sets.

Using this approach we obtained a large collection (18,000) of acceptable parameter sets (with cost less than twice the minimum). For this analysis, we varied 81 of 125 parameters, considering only parameters that affected the kinetics of the variables, fixing the rest of the parameters to their default values in the optimal parameter set (Section S2.5). We used this representative collection of parameter sets to investigate the robustness of model predictions, as described in the next section. We also used this collection to investigate the robustness of our estimates of parameter values as described in Section S2.8.

### Testing the model against observed phenotypes of mutant strains

In the previous section, we used quantitative data (time-courses and steady-state measurements) on specific molecular components of the nutrient signaling network to estimate parameter values in our mathematical model. In this section, we test the behavior of the parametrized model against qualitative experimental observations of the responses of *wt* and mutant strains to environmental perturbations. These ‘test case’ data were not used in the previous section to constrain the parameters.

From the literature we collected a list of 40 experiments (provided in Supplementary Table 4, Section S4) describing the qualitative phenotypes of mutant strains, typically colony growth phenotypes. We selected strains whose cellular states under nutrient shifts can be adequately characterized in terms of the transcription factors (TFs) in our model. To compare these experiments with our model predictions, we used the following strategy. For each colony growth experiment, we interpreted the observed phenotype in terms of the state (ON or OFF) of the nutrient responsive TFs in our model. Next we simulated each nutrient-shift or drug-treatment protocol for the specific mutant strain. We recorded the predicted steady-state values of the six TFs in our model, and converted these values into binary form (0 or 1) using thresholds corresponding to the half-maximal values in the *wt* simulation; thereby obtaining a Boolean vector of predicted TF states. Finally, we compared the predicted states of the relevant TFs to those interpreted from the experiment.

An important assay in the investigation of the nutrient signaling response is rapamycin treatment, constituting 22 of the 40 experiments in our collection. In *wt* cells, rapamycin treatment inhibits TORC1 activity, consequently activating nitrogen stress responses via TFs like Gln3, Gcn4, and Rtg1/3 (via the Tap42-Sit4 branch), inhibiting ribosome biogenesis via transcriptional repressors like Dot6/Tod6 (via the Sch9 branch), and indirectly causing a cell cycle arrest in the G1 phase. It appears that decoupling either one of these branches from TORC1 is sufficient to confer rapamycin resistance (cf. rapamycin treatment of *gln3*Δ*gcn4*Δ double mutant [80], and the Sch9^*DE*^ and *tap41-11* TORC1 insensitive strains [81]). Taking these observations into account, we attempted to carry out *in silico* experiments to explain our collection of experiments by examining different potential effects of rapamcyin treatment, including upregulation of nitrogen adaptation responses and downregulation of ribosome biogenesis. Using either of these definitions, the model could explain only about half of the rapamycin experiments that we have collected. We present this investigation in Section S3. Below, we focus on experiments not involving rapamycin treatment.

In order to characterize our confidence in model predictions, we recorded statistics on the number of times an experiment is predicted correctly across the representative collection of parameter sets (Table 2). Across all parameter sets, the model is able to correctly simulate up to 17 of 18 experiments. Five of these 18 experiments are explained by all of the parameter sets, and three of 18 are predicted by fewer than half of the parameter sets. The following paragraphs discuss some of the experimental observations that our model succeeds in explaining and some that our model fails to explain. The predictions described here are made using the optimal parameter set. Details of simulations are provided in Section S4.

**Table 2:**
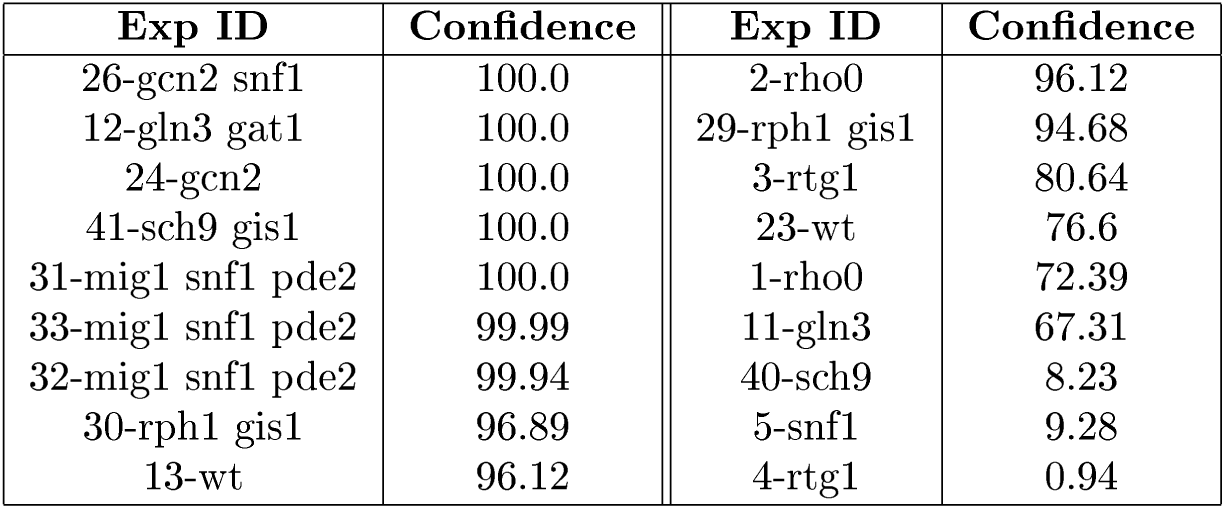
Confidence in qualitative predictions expressed as the percentage of parameter sets that make the correct prediction. The experiments are ordered in decreasing order of prediction confidence. The *in silico* experiments corresponding to the experiment IDs are presented in Section S4.

| Exp ID | Confidence | Exp ID | Confidence |
| --- | --- | --- | --- |
| 26-gcn2 snf1 | 100.0 | 2-rho0 | 96.12 |
| 12-gln3 gat1 | 100.0 | 29-rph1 gis1 | 94.68 |
| 24-gcn2 | 100.0 | 3-rtg1 | 80.64 |
| 41-sch9 gis1 | 100.0 | 23-wt | 76.6 |
| 31-mig1 snf1 pde2 | 100.0 | 1-rho0 | 72.39 |
| 33-mig1 snf1 pde2 | 99.99 | 11-gln3 | 67.31 |
| 32-mig1 snf1 pde2 | 99.94 | 40-sch9 | 8.23 |
| 30-rph1 gis1 | 96.89 | 5-snf1 | 9.28 |
| 13-wt | 96.12 | 4-rtg1 | 0.94 |

#### Nitrogen adaptation responses

An important function of the tricarboxylic acid cycle (TCA cycle) in central carbon metabolism is diverting the carbon backbone to amino acid biosynthesis pathways via *α*-ketoglutarate. Since the genes coding for the TCA cycle enzymes are present in the mitochondrial genome, mitochondrial dysfunction can severely affect respiration, resulting in the so-called *ρ*^0^ petite strains. Conceivably, these strains have a diminished capacity to divert carbon flux into amino acid biosynthesis. In yeast, the Rtg1/3 transcription factors control the expression of the *CIT2* gene, a TCA cycle enzyme encoded by the nuclear genome, which allows the diversion of carbon flux to amino acid biosynthesis pathways, under nuclear control (the so called retrograde pathway). Liu *et al*. studied *wt* (*ρ*^+^) and *ρ*^0^ strains to characterize the activity of the Rtg1/3 transcription factors by systematically varying the carbon and nitrogen content in the growth medium [82]. While the *ρ*^0^ strains can grow on media containing glucose only, raffinose only, and glucose supplemented with glutamate, the *rtg* strains in a *ρ*^0^ background only show growth on glucose + glutamate medium.

We simulated *wt* and *rtg* strains in such *ρ*^0^ petite strains. We assume that *ρ*^0^ strains have a lower level of glutamine pool replenishment than *ρ*^+^ strains. The *rtg1*Δ, *rtg3*Δ and *rtg1*Δ*rtg3*Δ strains in *ρ*^0^ backgrounds do not grow on media lacking glutamate, while supplementing the medium with glutamate results in *wt* growth [83]. Model predictions recapitulate the observations made by Liu *et al*. in media with and without glutamate (Sections S4.1 to S4.3).

#### The Snf1 pathway

While investigating the regulation of Gis1, Balciunas *et al*. studied the *mig1*Δ*snf1*Δ*pde2*Δ triple-mutant strain in various carbon sources and found that this strain is viable when grown on glucose and is inviable when grown on raffinose [84]. In order for this strain to grow on raffinose, Gis1 will have to be expressed. Our model predicts that Gis1 will be inactive during growth on both glucose and raffinose in this strain, indicating that the strain will show no growth on raffinose (Sections S4.31 and S4.32).

#### Model mismatches

The majority of parameter sets (> 50%) in our model succeed in explaining 15 of the 18 ‘test-case’ experiments (Table 2). We discuss the reasons for the three model mismatches below.

Gasmi *et al*. [85] reported that a *snf1*Δ strain cultured on minimal medium supplemented with ethanol grows slowly. In our model the glucose repression factor Mig1 will remain active in a *snf1*Δ strain, inhibiting any carbon adaptation responses, indicating no growth (Section S4.5). Liu *et al*. [82] showed that an *rtg1*Δ single deletion strain does not grow on medium containing a low amount of glutamate (0.02%). Our model predicts that this strain will still mount an Rtg1/3 response (Section S4.4). This might indicate a problem with the representation of nitrogen sufficiency in the model. Roosen *et al*. [19] showed that an *sch9*Δ strain grows when cultured on medium containing glycerol as a carbon source. In order to grow on glycerol, the Gis1 transcription factor must be activated via inhibition of PKA. Since our model assumes an inhibition of PKA via Sch9, an *sch9*Δ strain, with hyperactive PKA, will result in repression of Gis1 (Section S4.40).

### Predictions of global cellular responses to nutrient states

Motivated by our model’s success in predicting the qualitative phenotypes of 15 out of 18 mutant yeast strains, we used it to predict global cellular responses to nutrient conditions in a variety of mutant strains. We chose a list 16 mutant strains affecting all three nutrient signaling pathways: thirteen of these strains are gene deletion mutants including *lst4*Δ*lst7*Δ, *sch9*Δ, *gcn2*Δ, *snf1*Δ, *tpk1*Δ*tpk2*Δ*tpk3*Δ, *cyr1*Δ, *pde1*Δ*pde2*Δ*pde3*Δ, *sak1*Δ, *tor1*Δ, *ras2*Δ, *gtr1*Δ*gtr2*Δ, *bcy1*Δ, *ira1*Δ*ira2*Δ. and three are protein-sequence mutants: *GCN*2-*S*577*A* has lost the phosphorylation site for TORC1, and the two hypothetical strains *GLN* 3-Δ*ST* and *GLN* 3-Δ*TT* lack the phosphorylation sites for Snf1 and TORC1, respectively (see Figure 4).

**Figure 3:**
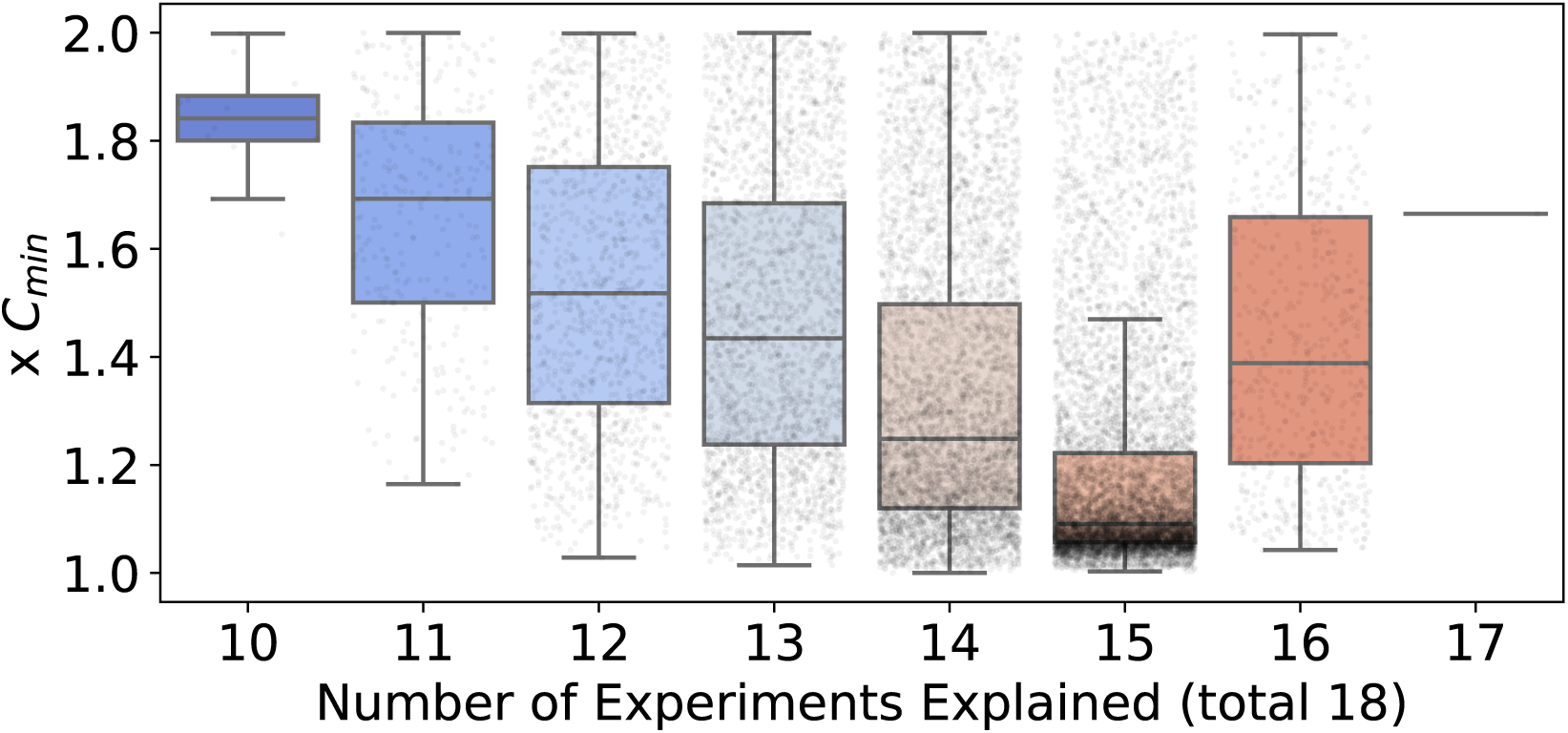
Dependence of model cost on explanatory capacity, across the entire collection of 18,000 alternative sets of parameter values. Model predictions of the 40 qualitative phenotype experiments were repeated for the collection of acceptable parameter sets. Each boxplot shows the distribution of cost-of-fit (to the quantitative measurements) for a given level of explanatory capacity (i.e. the number of qualitative phenotypes explained). Each point represents a parameter set. The cost, on the *y*-axis, is reported as a multiple of *C*_min_, the best observed cost across all parameter sets. The boxplot shows the median cost, and the whiskers extend to 1.5 times the interquartile range (IQR), or the last data point less than 1.5xIQR. Note that only parameter sets with a cost less than or equal to twice the *C*_min_ are reported.

**Figure 4:**
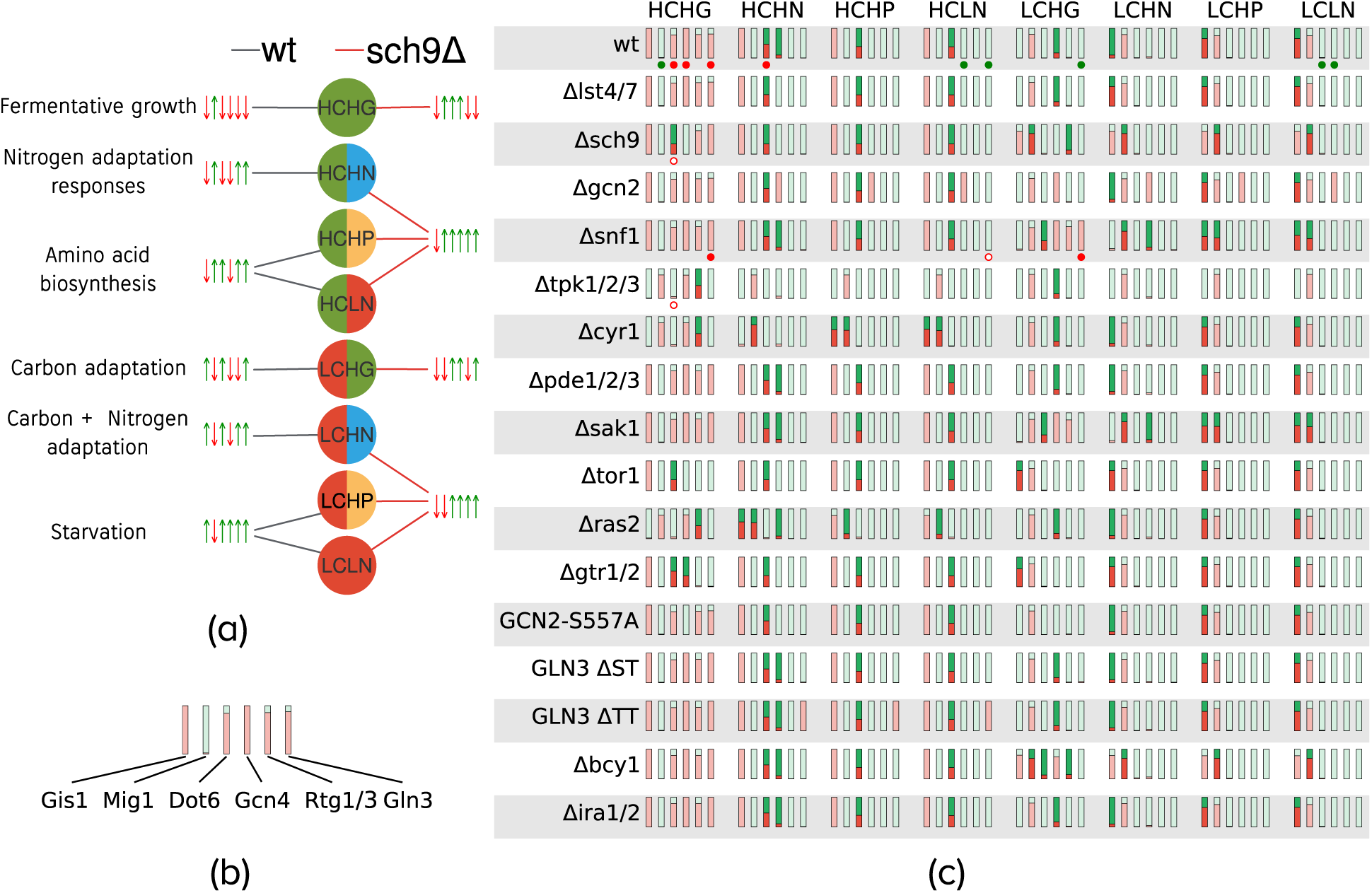
Global cellular responses to nutrient conditions. (a) Illustration of cellular states in response to various nutrient environments. The nutrient input is shown in the circles: L, H indicate low and high, respectively; C, G, N, P represent carbon, glutamine, ammonia and proline, respectively. The cellular state, defined by the set of transcription factor (TF) readouts, is shown by the upward pointing green (on) and downward pointing red arrows (off). The order of TFs is: Gis1, Mig1, Dot6, Rtg1/3, Gcn4, Gln3. Each nutrient input is connected to a cellular state by an edge. The figure shows a comparison between the predicted states for two strains, *wt* (black edges) and *sch9*Δ (red edges), using the optimal parameter set. Qualitative descriptions of the cellular states in the *wt* simulations are provided on the far left. (b) Representation of the consensus prediction of cellular state across 18,000 alternative parameter sets. The example shows the prediction for *wt* cells under HCHG condition. The height of a red/green bar represents the fraction of parameter sets that predicted the corresponding state to be off and on, respectively. We consider the prediction to be robust if greater than 90% of parameter sets are in agreement, shown using light green or red. As shown, most parameter sets are in agreement regarding the states of the readouts for *wt* cells under high carbon, high glutamine. (c) The robustness of global state predictions across 17 strains and 8 nutrient conditions. Fragile predictions are indicated in bright green and red. Additionally, direct measurements of TF activities obtained from the literature are indicated using filled and empty circles below the corresponding readouts; a filled circle indicates that model prediction agrees with experimental data while empty circles represent a model mismatch. Green and red circles indicate that the readout is on or off, respectively.

We characterized each strain’s response to eight nutrient conditions (qualitatively low and high inputs for ‘Carbon’, ‘Glutamine’, ‘Ammonia’, and ‘Proline’). In Figure 4, these qualitative states are denoted using ‘H’ and ‘L’ respectively for high and low, and the nutrient input is denoted using ‘C’, ‘G’, ‘N’, and ‘P’ for Carbon, Glutamine, Ammonia and Proline, respectively. (Note that ‘LN’ represents nitrogen starvation.) For each simulation, we recorded the steady state values of the six transcription factors (TFs) in our model and thresholded the TF activities using their half-maximal values in the *wt* simulations. We thus obtained a Boolean vector describing the phenotypic state of a cell under different nutrient conditions, with each TF taking a value 0 (off) or 1 (on). The predicted states for the 17 strains (including *wt*) over all nutrient conditions (as predicted by the model using the optimal parameter set) are provided in Supplementary Table S5.

Figure 4(a) provides an example of the predicted cellular states for *wt* and *sch9*Δ cells for each of the eight nutrient conditions. We observed that, for each nutrient input, the state of the *wt* strain was different from that of the *sch9*Δ strain. Repeating these simulations across the 17 strains and the 8 nutrient conditions yielded a total of 136 predicted cellular states. In order to gain confidence regarding model predictions, we repeated these *in silico* experiments for each of the 18,000 alternative parameter sets. Using two parameter sets, if the predicted activity of a particular TF in a given experiment is identical, the two sets are considered to agree on the prediction. We define fragile predictions to be the experiments where fewer than 90% of the parameter sets were in agreement about the outcome of a simulation. The results of this analysis are summarized in panel (c) of Figure 4, where the consensus of the predictions across all tested parameter sets is indicated by the brightness of the bars. We observed that, in the HCHG simulations, the predictions exhibited consensus across most strains for all TFs. Note that in Figure 4(c) there is significant disagreement between the predictions made by the parameter sets regarding the state of Dot6, the repressor of RIBI, in the case of high carbon (HC) irrespective of the nitrogen sufficiency. This is likely due to the lack of experimental data constraining Dot6 dynamics. Furthermore, we observe that the HP and LN columns are identical for both HC and LC, implying that the model fails to distinguish between the high proline and nitrogen starvation conditions, whereas proline, a poor source of nutrients would still be expected to serve as a substrate in order to maintain intracellular amino acid reserves. We believe that this failure is a consequence of the fact that our model does not include the effects of metabolic feedback on nutrient signaling, thus rendering the two poor-nitrogen conditions indistinguishable.

Finally we observe that the alternative parameter sets disagree about the state of Gis1 across most strains during nitrogen stress under carbon starvation (LCHN, LCHP and LCLN columns in Figure 4(c)). Gis1 is jointly regulated by Sch9 and PKA. However, the data used to calibrate the model come from experiments related to the PKA pathway only [73]. In order to refine our predictions for Gis1, the model will need to be recalibrated with Gis1 data obtained from nitrogen-shift experiments, when they become available.

## Discussion

Since the default state of yeast cells in the presence of nutrients is exponential growth, the coordination of various nutrient levels within the cell is of paramount importance. The integration of extracellular nutrient signals is distributed broadly across the cAMP/PKA (glucose signaling), Snf1 (carbon adaptation) and TORC1 (nitrogen signaling) pathways, along with many crosstalk interactions among members of these pathways and their downstream targets. These interactions comprise the nutrient-signaling and decision-making system in budding yeast. In this paper we have proposed a mathematical model of this system, based on well-studied interactions of the individual pathways and less-studied crosstalk between the pathways (Figure 1). Our results demonstrate that the proposed mechanism can account for many aspects of nutrient signaling in budding yeast, as observed in diverse nutrient-shift experiments (Figure 2).

### Global cellular responses to nutrient shifts

Two experimental observations highlight the crosstalk between nutrient signaling pathways: (1) cells facing nitrogen stress in the context of carbon abundance can redirect carbon flux from the TCA cycle into amino acid biosynthesis [86], and (2) acute nitrogen starvation results in down-regulation of glucose fermentation [87]. These interactions are regulated by metabolic enzymes that are under the regulation of various transcription factors (TFs) included in our model. To get a sense of the cell’s response to complex nutrient shifts, we thus need to examine the activities of the TFs spanning carbon and nitrogen responses.

In our model, we use the Boolean states of the TFs to define the state of a cell. Each of these predicted states is deemed ‘robust’ or ‘fragile’ based on the consensus among predictions from a representative collection of 18,000 alternative sets of parameter values (see Figure 4 (c)). Robust predictions that are subsequently shown to be incorrect indicate parts of the model that need further refinement. Fragile predictions, on the other hand, indicate those parts of the model that are poorly constrained by available data. Experimental tests of a fragile prediction can invalidate those ‘alternative’ parameter sets that make the incorrect prediction. Hence, both robust and fragile predictions suggest potentially informative experiments that can be used to further constrain the model. In the following paragraphs, we propose experiments that we deem informative based on some fragile predictions.

The Rtg1/3 transcription factors play the crucial role of diverting carbon flux into amino acid biosynthesis. Rtg1/3 are expected to be activated during nitrogen stress in carbon-rich conditions. Figure 4 (c) indicates that the model makes fragile predictions for Rtg1/3 activity in the HCHG condition in three mutant strains, namely *tpk1/2/3*Δ, *cyr1*Δ, and *ras2*Δ. In these three strains PKA is inactive. Because, in our model, PKA activates TORC1 (via Sak1 and Snf1), TORC1 activity should be low in these mutants. However, since high glutamine activates TORC1, this protein receives contradictory signals from the carbon and nitrogen pathways. As a result, Rtg1/3 may be either active or inactive, depending on the values of the parameters in our model simulations. Hence, data pertaining to Rtg1/3 activity in these strains will help to narrow down the acceptable sets of parameter values.

The Mig1 transcriptional repressor implements glucose repression, i.e., in the presence of glucose, Mig1 inhibits expression of genes involved in carbon adaptation responses. During glucose starvation, Snf1 inhibits Mig1 by localizing it to the cytoplasm. Interestingly, our model predictions for the state of Mig1 are fragile in LCHN, LCHP, and LCLN conditions in two separate strains, *sch9*Δ and *bcy1*Δ. (Note: Sch9 inhibits PKA via Bcy1.) In this case, the fragile predictions of Mig1 activity are consequences of contradictory signals received by Snf1. In LC, Snf1 should be active and Mig1 inactive. However, in these mutants, despite LC conditions, PKA is active, Sak1 is inactive, and there is little or no activation of Snf1. Hence, whether Snf1 is active or inactive depends sensitively on precise parameter values. Therefore, measurements of Mig1 activity under glucose starvation conditions in the *bcy1*Δ strain will also help to narrow down the acceptable sets of parameter values.

An important determinant of cell growth under diverse nutrient conditions is ribosome biogenesis, which in our model is regulated by the Dot6/Tod6 repressors. Interestingly, Figure 4 (c) shows that Dot6 predictions are fragile under HCHN, HCHP, and HCLP conditions across most mutant strains, indicating that the strength of crosstalk between carbon and nitrogen signaling pathways is crucial in determining Dot6 activity. Measurements of Dot6 activity in the mutant strains identified by Figure 4 (c) will provide important constraints on these crosstalk interactions. However, we must bear in mind that other factors controlling ribosome biogenesis, such as Sfp1, are not yet included in our model.

### Nutrient adaptation responses and global metabolic feedback

Our proposed mechanism successfully recapitulates many features of the nutrient response system in budding yeast over short timescales (30 minutes). During nutrient stress, the cell mounts a variety of adaptation responses in order to maintain intracellular nutrient pools. Our model captures the activation of stress TFs immediately after a nutrient shift. However, on longer time scales of about 50 minutes, data from Granados *et al*. reveal that many of these stress TFs are eventually turned off [88]. Our model simulations do not capture this feature. How can this inactivation of TFs be explained? The stress TFs activate various biosynthetic pathway enzymes which restore the flux of metabolites through anabolic processes. Consequently, the starvation signals are turned down, inactivating the stress TFs themselves. Metabolic processes are known to play important roles in cell growth and division [89]. Because the mechanisms by which metabolic responses feedback on the nutrient signaling network are complex and poorly understood, we have not yet tried to include these effects in our model (Figure 5).

**Figure 5:**
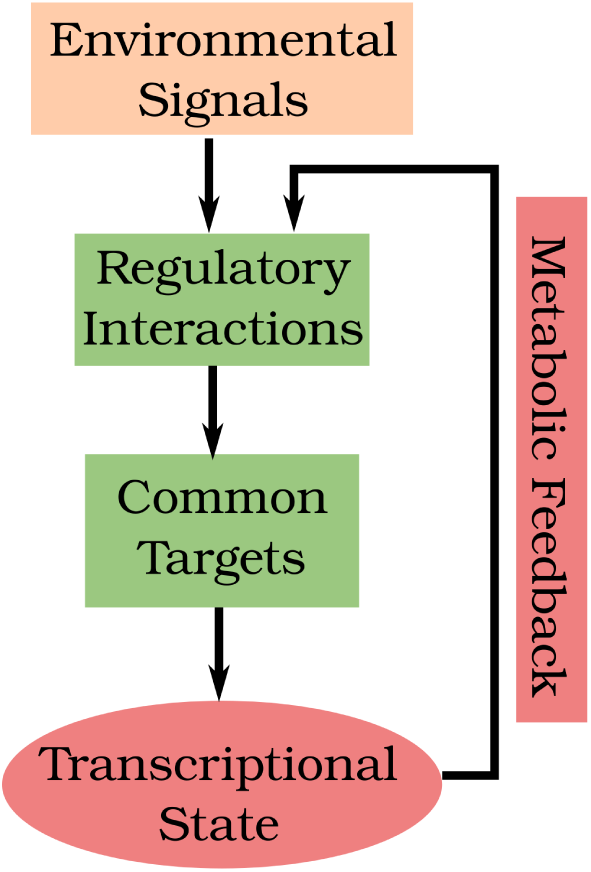
During nutrient stress, upstream sensing mechanisms transduce stress signals to downstream transcription factors. Stress signals can originate from both the extracellular environment and intracellular nutrient reserves. The latter are sensed indirectly via the levels of uncharged tRNAs (in the case of amino acids) or metabolic intermediates that are directly sensed by signaling molecules. The drops in flux through metabolic pathways result in upregulation of specific biosynthetic pathways and other adaptation responses, indicated in red at the bottom of the figure. Subsequently, metabolic fluxes are remodeled, leading to inactivation of stress responses. This ‘metabolic feedback’, which is currently not included in our model, determines the long-term responses of the nutrient signaling network.

In general, how to incorporate metabolic fluxes in nutrient signaling networks is a difficult challenge. Three aspects stand out: (1) the accurate representation of intracellular nutrient pools, (2) the metabolic fluxes spanning carbon and nitrogen metabolism, and (3) the dynamics of regulation of metabolic enzymes by the upstream signaling network. By providing a mechanistic model of the nutrient signaling system in yeast, our model is well poised to serve as a bridge between global metabolism and a wide range of cellular adaptation responses ranging from ribosome biogenesis to autophagy.

## Methods

Ideally, dynamical models of biological regulatory systems are based on biochemical details of the system under consideration, but oftentimes these details are obscure. Furthermore, the rate constants of enzyme-catalyzed reactions in a detailed mechanism are difficult to measure experimentally, posing a major hurdle for model parametrization. In this paper, we employ a general framework that circumvents these problems by using generic functions to describe biochemical reactions that occur on three different timescales: slow (e.g., gene expression, protein synthesis and degradation), intermediate (e.g., post-translational modifications) and fast (e.g., complex formation and dissociation). This approach is called the ‘standard component’ modeling strategy [90, 91]. We provide a brief description of this strategy in Section S1 and a full list of the model equations in Section S1.1.

In order to estimate numerical values of the parameter in the model, we need quantitative experimental data from the literature. We focused on published measurements of molecular species as functions of time, which we digitized using WebPlotDigitizer [92], and on data from electrophoretic gels, which we digitized using the ‘Analyze Gels’ option in ImageJ [93]. As described in Results, all values obtained from a given publication are scaled between 0 and 1 with respect to the lowest and highest reported values in a particular experiment.

In order to represent a nutrient shift, we first simulate the model to steady state using parameters representing the pre-shift condition. For example, in order to represent an amino acid-replete but carbon-poor pre-shift condition, we first set the ‘Carbon’ parameter to 0 and the ‘Glutamine’ parameter to 1, simulate the model, and record the steady state. Then, in order to simulate the shift to a carbon-rich environment, we set the ‘Carbon’ parameter to 1 while maintaining ‘Glutamine’ parameter at 1, continuing the simulation from the previous steady state.

### Parameter estimation and robustness analysis

During the model construction phase, we parametrized individual parts of the network corresponding to each signaling pathway with time-course data relevant to that pathway. We defined a quadratic cost function, as described in Section S2.1, and used the Levenberg-Marquardt Least Squares Optimization method (the `leastsq` function in the Scipy Python library) to minimize the cost (i.e., to improve the goodness-of-fit of the simulated trajectories to the experimental data). Next, we combined the subnetworks and supplemented the cost function with additional time-course and ‘perturbation’ data (see Figure 2) relevant to regulators across the signaling pathways. We observed that the quality of the least-squares fit declined as the complexity of the model and the data increased. Hence, after an initial manual tuning of the full set of 128 parameters, we used a Markov-Chain Monte Carlo (MCMC) sampling procedure (described in Section S2.2) to find an ‘optimal’ (least-cost) parameter set at the end of 10,000 steps of MCMC sampling. The parameter values constituting the optimal set are presented in Table S2.

In order to better understand the reliability of the optimal parameter values, we performed a global parameter-robustness analysis, as described in Section S2. The goodness-of-fit cost function defined in the previous paragraph is known to have a characteristic ‘stiff’ and ‘sloppy’ shape in the neighborhood of the best fitting parameter set. Our goal was to characterize the shape of the cost function and, in the process, obtain a large collection of alternative parameter sets that provide a ‘good’ fit (if not the ‘best’ fit) to the data. To this end, we started by calculating the cost function on a Latin Hypercube sample around the optimal set of parameter values obtained from the MCMC procedure. Then we constructed a quadratic approximation (the Hessian matrix) of the cost function around the local minimum. Using this approximate Hessian, we proposed new sets of parameter values constrained by the directions in parameter space corresponding to the largest eigenvalues of the Hessian, namely the ‘stiff’ directions. By limiting parameter variations in the stiff directions and permitting large variations in the sloppy directions, we were able to generate thousands of potential parameter sets that differed significantly from the optimal set but still might provide an acceptable fit to the experimental data. For each of these proposed parameter sets, we evaluated the model cost, accepting the set if its associated cost was less than three times the lowest cost. Nest, we recomputed the Hessian matrix using the cost evaluations over the ‘accepted’ sets of parameter values, and we repeated the procedure of proposing new parameter sets based on the stiff and sloppy directions of the Hessian. At the end of four iterations of this process, we had collected 24,000 parameter sets with cost < 3 × C_*min*_. We winnowed this collection down to 18,000 parameter sets with cost < 2 × C_*min*_. Using this large collection of ‘acceptable’ parameter sets, we studied the robustness of our parameter estimates, and the results of this analysis are presented in Section S2.

## Acknowledgements

A grant from the National Science Foundation (DBI-1759858) supported this work. APJ would additionally like to acknowledge the Genetics, Bioinformatics, and Computational Biology (GBCB) and (Computational Tissue Engineering) interdisciplinary graduate education programs at Virginia Tech for funding. The authors would like to thank Dr. William Baumann for providing the derivation of the Hessian-guided search method, and for helpful discussions.

## Availability

The data curated from the literature is provided in additional Microsoft Excel files.

**Supplementary-Table-1**.**xlsx** provides parameter values from the optimal parameter set.

**Supplementary-Table-2**.**xlsx** provides literature curated timecourse data.

**Supplementary-Table-3**.**xlsx** provides literature curated perturbation data.

**Supplementary-Table-4**.**xlsx** provides literature curated qualitative shift experiments and their representations in the model. The ‘Gln3Gcn4’ sheet specifies rapamycin experiments using Gln3 and Gcn4 as readouts. Similarly the ‘Dot6’ sheet specifies rapamycin experiments using Dot6 as the readout. The non-rapamycin treatment experiments are identically defined in both sheets.

**Supplementary-Table-5**.**xlsx** provides global TF state predictions using the optimal parameter set.

The code and data used in the construction, calibration and analysis of the model is available at https://github.com/amoghpj/nutrient-signaling. The model is additionally provided in the SBML format. To enable ease of interaction with code, the datasets mentioned above are also provided as YAML files in this repository.

## S1 Model Construction

The Standard Component Modeling (SCM) framework used to create our model of nutrient signaling is described here in brief. SCM classifies biochemical reactions based on their time scales. The slow timescale reactions (*Class I*) are represented by mass-action rate laws

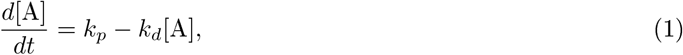

where *k*_*p*_ is the rate of production and *k*_*d*_ is the rate of degradation of species A.

The intermediate timescale reactions (*Class II*) are represented by a generic, sigmoidal, ‘soft-Heaviside’ function. For example, a protein P undergoing a post-translational modification would be described by the differential equation

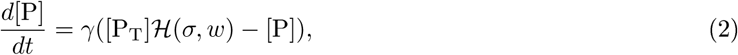

where [P_T_] represents the total amount of species P, and ℋ is the soft-Heaviside function, given by

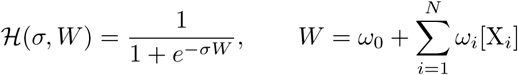

ℋ is a sigmoid function used to represent a switch-like biochemical mechanism. Here the *W* term is composed of a linear combination of the concentrations of the regulators X_*i*_ of P with appropriate signed coefficients *ω*_*i*_. The *γ* parameter governs the time scale of this reaction. In this work, [P_T_] = 1 for every *Class II* variable in a wild-type cell, and we set [P_T_] = 0 to simulate a cell deleted for the gene encoding P. Thus, the value of a *Class II* variable, which lies between 0 and 1, denotes the fraction of the given signaling component in a particular post-translational modification state.

Finally the reactions occurring on fast time scales, such as the formation of protein complexes, are represented by the *Class III* equations governed by the stoichiometric relationships between the constituents of the protein complexes. Thus, the stoichiometric association of components *X* and *Y* to produce a complex *C* would be represented as

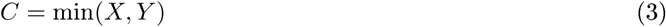

The kinetic expressions defining the ODE model are presented in the next section. A majority of the expressions (21 out of 25) are *Class II* equations. The model has four *Class I* equations, describing the dynamics of Glutamine and cAMP, Ribosomal components (Rib), and Protein accumulation. Finally the model has two *Class III* equations: First, the charging of tRNAs as a fast association between total uncharged tRNAs and intracellular amino acids, represented by Glutamine. Second, the assembly of active Ribosomes (aRib) is represented as the fast association of Ribosomal components (Rib) and unphosphorylated initiation factor eIF2*α*.

### S1.1 Kinetic expressions

#### Nutrient signal sensing and transduction

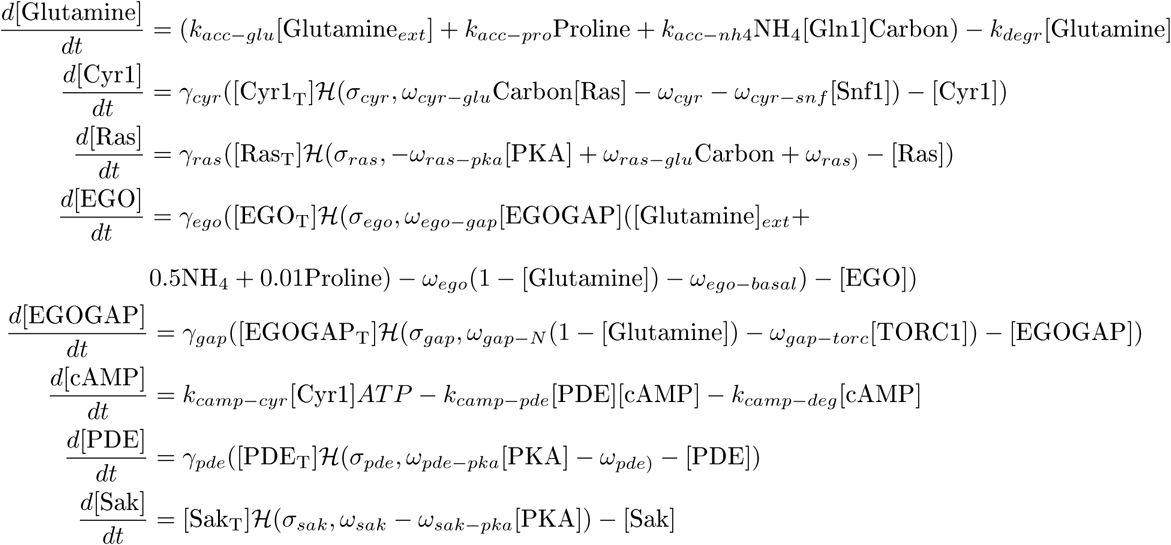

#### Master regulators

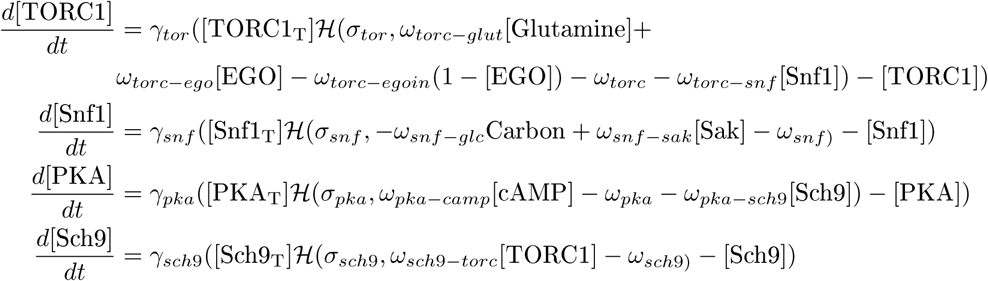

#### Downstream responses

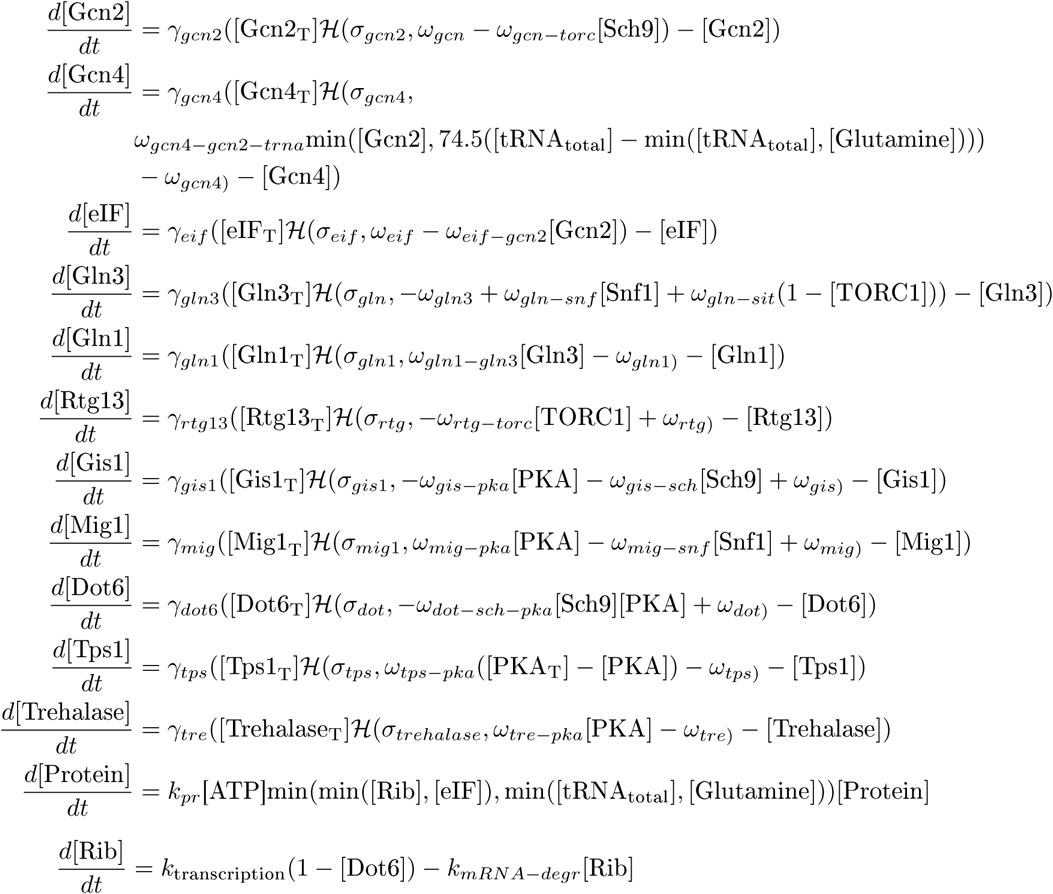

Some heuristics have been used in order to model the following components.

##### Amino acid sensing

The detailed mechanism of amino acid sensing is beyond the scope of this model. In order to simplify this pathway, it is assumed that the EGO complex integrates all nitrogen signals. Thus the activation of the EGO complex is determined by the strength of the activating signal, which is represented by

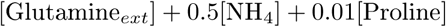

with the coefficients being derived from best fits to Sch9 activation data from [30].

##### Ribosome Biogenesis

Here we abstract away the complexities of the synthesis of ribosome components and represent the ‘synthesis’ of ribosomes by a mass action law, regulated by the RIBI repressor Dot6.

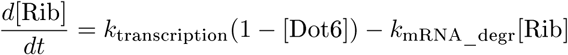

##### Protein Translation

We abstract away the complexities of ribosome assembly and assume that a ribosome is functional when a stoichiometric ratio of the ribosomal precursor (Rib), charged tRNA, and unphosphorylated initiation factor eIF2*α* is present.

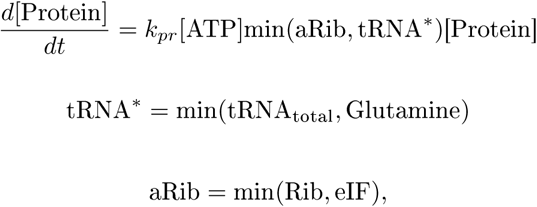

where tRNA^*^ represents the charged tRNA pool, and aRib represents the actively translating ribosomes.

## S2 Robustness analysis

This section describes the details of the parameter robustness analysis. The motivation for this analysis is the concept of model sloppiness developed by Gutenkunst *et al*., 2007 [78] who investigated the characteristics of a cost function quantifying the deviation of model predictions from a set of experimental data. The value of the cost function depends on the values of the parameters in the model. In their investigations Gutekunst *et al*. found that the cost as a function of model parameters typically exhibits a ‘stiff/sloppy’ structure, i.e., while one can identify a few parameter combinations that tightly constrain the cost function, a majority of parameter combinations do not significantly constrain the cost function. Here, we are interested in the so-called stiff parameter directions, which are highly constrained by data. We carry out this investigation in the following stages:

1. We define a cost function that measures the fit between the experimental data and model predictions, described in Section S2.1.
2. We use the cost function to improve the global fit of the model to the data using an MCMC sampling strategy described in Section S2.2.
3. We approximate the structure of this cost function in parameter space by computing the Hessian of this surface around the optimal parameter set using the method described in [79]. Details of this method are provided in Section S2.3.
4. We generate a sample of parameter sets constrained by the eigenvectors of the Hessian matrix, and iteratively refine the Hessian, as described in Sections S2.4 and S2.5.
5. Finally, we study the properties of this refined Hessian to identify the stiff and sloppy parameter directions in our model, given the set of curated experimental data used to constrain the model. This is described in Sections S2.6 to S2.9.

### S2.1 The goodness-of-fit cost function

This quadratic cost function *C*(**p**), which includes both the time series data and the steady-state pertubation data as described in ‘Results’, is defined as follows:

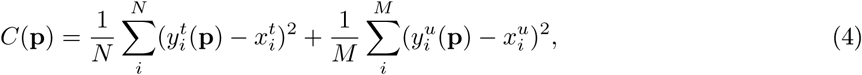

where *N* is the number of time points (*t*) and *M* the number of perturbations (*u*), **p** is the candidate parameter vector, *y*(**p**) is the model prediction and *x* is the literature-derived activity of the variable under consideration. While the number of time points exceed the number of perturbation data points, we do not preferentially weight one type of data over the other.

### S2.2 MCMC sampling to improve estimate of parameter values

**Figure S1:**
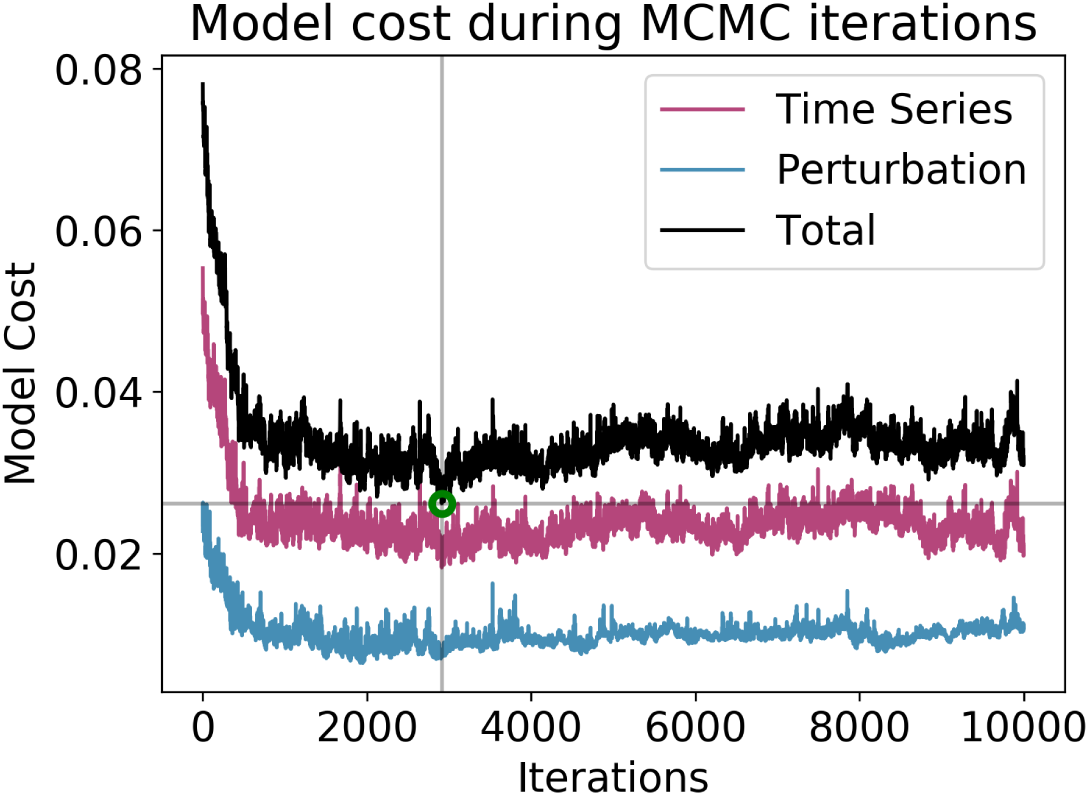
Results of MCMC sampling. The black line shows the total model cost. The contributions from the time course term and from the perturbation term are shown in pink and blue, respectively. The lowest cost, designated by the green circle at iteration 2900, defines the ‘optimal’ parameter set.

Having defined the quadratic cost function, we used a Markov Chain Monte Carlo (MCMC) sampling strategy to improve the fit to the data. Briefly, in every MCMC iteration, the last accepted parameter set is perturbed as follows: for each parameter with value *p* in the last accepted set, a new value *p*′ is sampled from a normal distribution 𝒩(*μ* = *p, σ* = 0.025*p*). The cost function is evaluated for this new parameter set and this set is accepted with probability *e*^−*β*Δ*C*^, where Δ*C* = *C*(*p*′) −*C*(*p*). We chose *β* = 3.6 based on the magnitude of the change in cost that we observed in each iteration. Starting from the hand-tuned parameter set, we repeat MCMC sampling 10,000 times. The change in cost across the iterations is shown in Figure S1.

### S2.3 Defining a Hessian approximation based on a sample of parameter sets

In this section, we derive the expressions used to compute the Hessian.

Let **p**^*^ denote the parameter set that minimizes *C*(**p**), i.e., *C*(**p**^*^) = *C*_min_, then

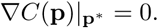

Thus, for every **p** in the neighborhood of **p**^*^, we can carry out a Taylor series expansion around **p**^*^. Omitting the higher order terms:

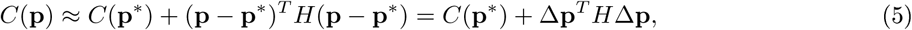

where 2*H* = ∇^2^*C* is the Hessian of the cost function.

Next, we define some notation as introduced in Magnus and Neudecker [94]. For an *m* × *n* matrix *A*, the vectorization operation vec(*A*) results in a *mn* × 1 column vector that stacks the columns of *A*. If *A* is an *n* × *n* symmetric matrix, then the operation vech(*A*) stacks the lower triangular columns, yielding an 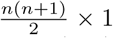 column vector. There exists a unique matrix *D*, called the duplicator matrix, with dimensions 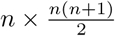, and a unique matrix *L* called the eliminator matrix with dimensions 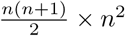, such that

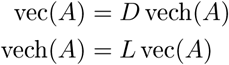

The Kronecker product (denoted by ⊗) of an *m* × *n* matrix A and an *s* × *t* matrix B is an *mn* × *st* matrix

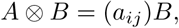

where *a*_*ij*_ is the *ij*^th^ entry of *A*. For any three matrices *A, B* and *C* the following holds true

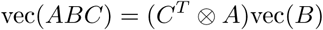

Suppose we choose *S* parameter sets **p**_*i*_, ≤ 1 *i* ≤ *S* in the neighborhood of **p**^*^. We can construct a quadratic error function *E*_*H*_ which will be minimized when *H* approximates the true Hessian of the function.

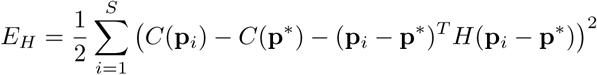

Using Δ*C*_*i*_ = *C*(**p**_*i*_) − *C*(**p**^*^) and Δ**p**_*i*_ = (**p**_*i*_ − **p**^*^), we have

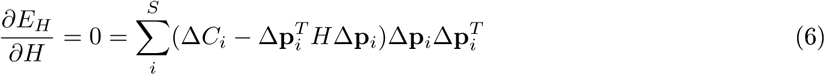

We wish to solve for *H* given a set of parameter vectors **p**. For a model with *k* parameters, we will have to solve for *k*^2^ terms in *H*. However, we can decrease the size of this problem by considering the fact that the Hessian should be a symmetric matrix. Thus, we need to solve only for the terms in the lower triangle.

Simplifying Equation (6), we have

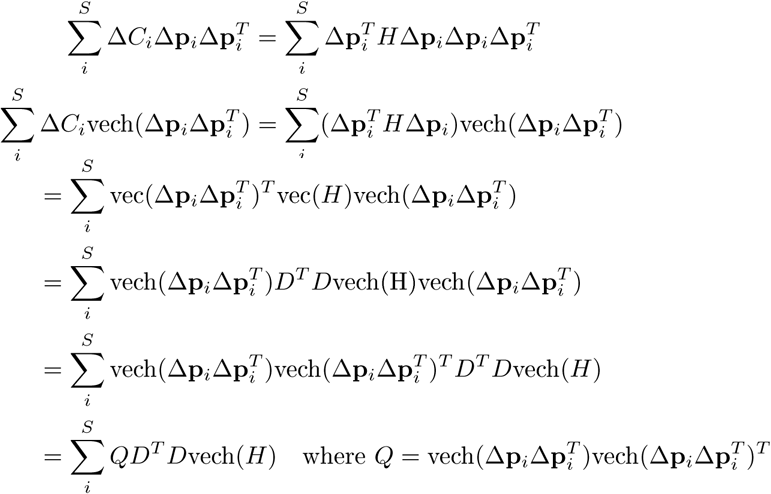

If we define *R* =Σ_*i*_ Δ*C*_*i*_ vech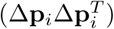, we have that

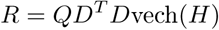

In other words,

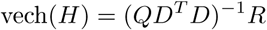

and

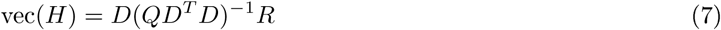

Here, *R* is symmetric and *D* is the appropriate duplicator matrix.

### S2.4 Sampling new parameter sets constrained by the approximate Hessian

Using the approximate Hessian matrix computed as described in Equation (7), we next wish to use the eigenvectors of this matrix to contrain the search for new parameter vectors. Intuitively we wish to avoid the eigenvector directions corresponding to ‘large’ eigenvalues which are the stiff directions. We know that the dimensions of the cost ellipsoids are proportional to 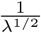 where *λ* is an eigenvalue of the Hessian. Thus we can weight the eigenvectors by a factor of 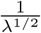, which favors the sloppiest eigenvector directions. In practice we want a candidate parameter vector to respect all the stiff and sloppy directions. We first generate a random vector *α*, which we then transform to respect the stiff and sloppy directions. We describe the transformation matrix below.

We start by translating the frame of reference our system to **p**^*^ so that we can generate vectors Δ**p** = **p**−**p**^*^which produce a relative increase in model cost Δ*C* = *C* − *C*_min_. Rewriting Equation (5),

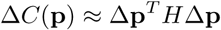

To avoid an ill-conditioned Hessian, where the eigenvalues span many orders of magnitude, we choose 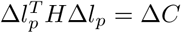 where Δ*l*_*p*_ = log(**p**) − log(**p**^*^), with the logarithm of a vector being taken elementwise.

To sample from this new ellipsoid, we first create a random vector *α* which will lie inside the cost ellipsoid. we sample a vector 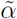 of random numbers drawn from 𝒩(0, 1). Next, we compute *α*

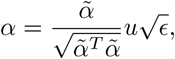

where *u* is a scalar drawn from the uniform distribution on [0, 1] and *ϵ* = 2*C*_*min*_ = 2 × 0.026 is the maximum value of Δ*C* we wish to consider. The factor 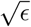 scales the magnitude of the unit vector 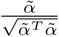to the edge of the ellipsoid. The final factor *u* ensures that the vector lies inside the ellipsoid, as we want to sample the volume, not just the surface of the ellipsoid.

The final step needed to generate a candidate parameter vector is to transform it to respect the stiff and sloppy directions. Using an eigenvalue decomposition, we write *H* = *V* Λ*V*^*T*^ (where *V* is the eigenvalue matrix and Λ is the eigenvector matrix). (Note that we compute the absolute values of the eigenvalues and replace every eigenvalue that is less than 0.1 by 0.1. This step ensures that *H* is positive definite, but limits the length of the longest ellipsoid axes.) Our constrained parameter vector should take the form Δ*l*_*p*_ = *V* Λ^−1*/*2^*α*. This will satisfy the cost constraint as follows

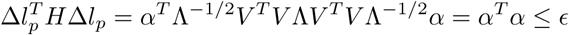

Reordering terms, we have

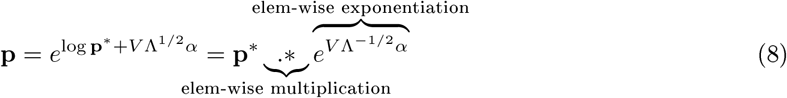

### S2.5 Iterative refinement of the approximate Hessian

The following steps describe our methodology to initialize a approximation to the Hessian and then refine this approximation iteratively.

1. To compute an initial approximation of the Hessian, we used Latin Hypercube Sampling around 2.5%ranges around **p**^*^ in order to sample parameter vectors close to **p**^*^. We evaluated the cost for each parameter set. For each range explored, we recorded the parameter sets with cost less than 2*C*_min_.
2. Next, using Equation (7), an approximate Hessian was constructed using the accepted parameter sets from the Latin Hypercube sample. This step was designated as iteration 0.
3. Using the approximate Hessian generated in the previous step, 30,000 parameter sets were generated using Equation (8). The goodness-of-fit cost was evaluated for each parameter set and any parameter set satisfying a cost cutoff of 3*C*_min_ was accepted.
4. Using these accepted parameter sets, the approximate Hessian was recomputed, a new ensemble of 30,000 parameter sets was generated, and the cost evaluation and parameter set acceptance procedure was repeated, while continually expanding the ensemble of accepted parameter sets.
5. The previous step was repeated four times.

In our parameter search we fix the values of the total amounts of protein, the P_T_ parameters to 1.0 since we currently do not have accurate abundances of the regulators in the model, accounting for 21 parameters. We also set the *sigma* parameters in each *Class II* equation to their nominal values presented in Table S2, accounting for another 21 parameters. Lastly, 5 parameters serve as model inputs, namely ATP, Carbon, Glutamine_*textext*_, Ammonia, and Proline. Thus we fix the values of 49 of the 128 kinetic parameters, The remaining 81 parameters were varied in this analysis. At the end of the four iterations, we obtained a total of 24,066 parameter sets.

### S2.6 Fewer than 16% of the eigenvectors are required to capture all the stiff directions

We studied the eigenvector of the refined-approximate Hessian *H* in order to identify the parameters that contribute to the stiff directions. We deemed a parameter as making a substantial contribution to an eigenvector if the absolute value of its ‘loading’ (i.e., its weight in the eigenvector) was greater than one standard deviation of all the loadings across all eigenvectors. Figure S2(a) shows the number of unique parameters with substantial contributions to the ordered eigenvectors of *H*. Since we vary 81 parameters, the matrix *H* has 81 eigenvectors. We observe that 90% (72 of 81) of the parameters make substantial contributions to the first 13 eigenvectors. Thus, all parameters contribute substantially in the first 16% (13 of 81) of the stiff directions.

Finally, we observe an initial slump in the number of unique parameters, implying that only 28 parameters contribute substantially to the first seven stiff directions, i.e., around 34% of the parameters contribute to the stiffest directions, indicated by the red lines in Figure S2(a).

**Figure S2:**
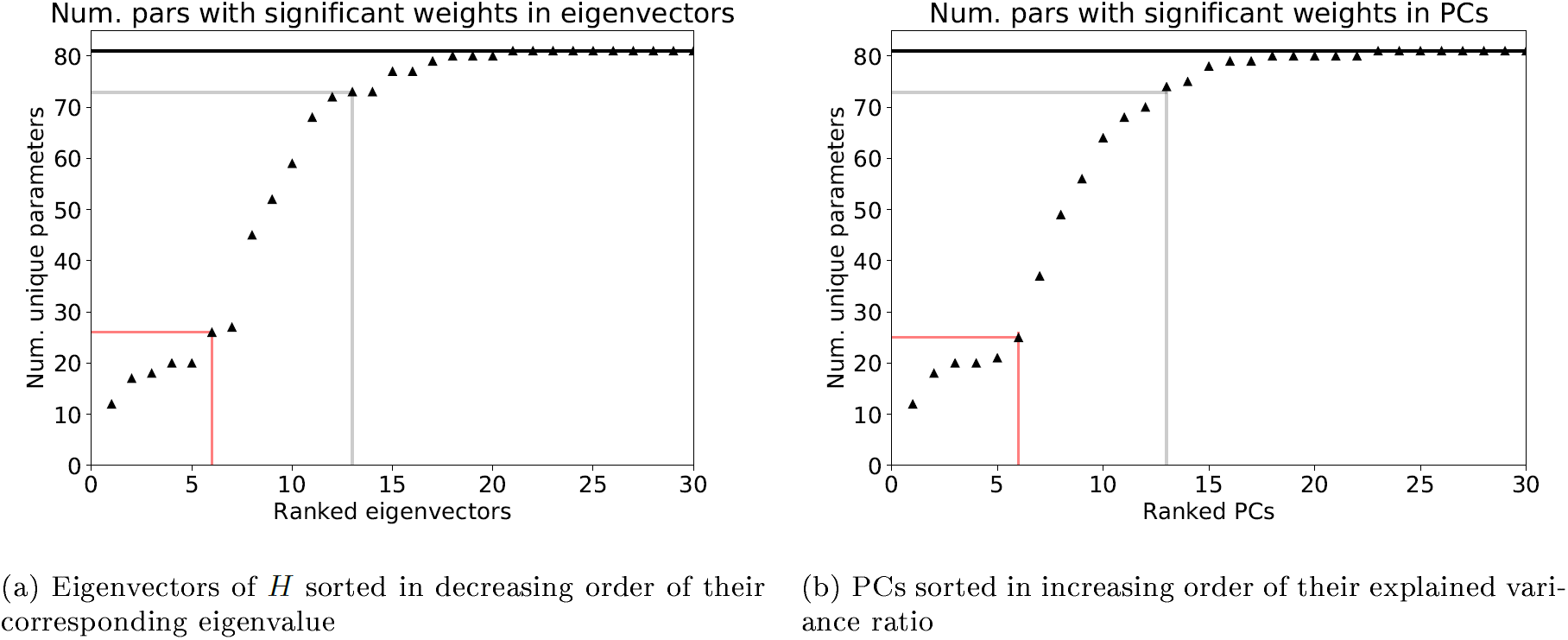
Comparison between the parameters with high weights in eigenvectors. On the *x*-axis are the sorted eigenvectors, such that the eigenvector corresponding to the largest eigenvalue gets a rank of 1. Each plot shows the number of unique parameters with substantial coefficients in the each eigenvector. The gray lines mark the number of eigenvectors required to capture 90% of the parameters. The black horizontal line marks the total number of parameters varied in our analysis, namely 81. The plot on the left is derived from the eigenvectors from the approximate Hessian, whereas the plot on the right represents the results from the principal components resulting from carrying out PCA on the collection of parameter sets, i.e., the eigenvectors of the covariance matrix sorted by the inverse of their eigenvalues. The red lines indicate a characteristic inflection point in the trend of parameters contributing substantially to eigenvectors which we interpret as the stiff directions.

**Figure S3:**
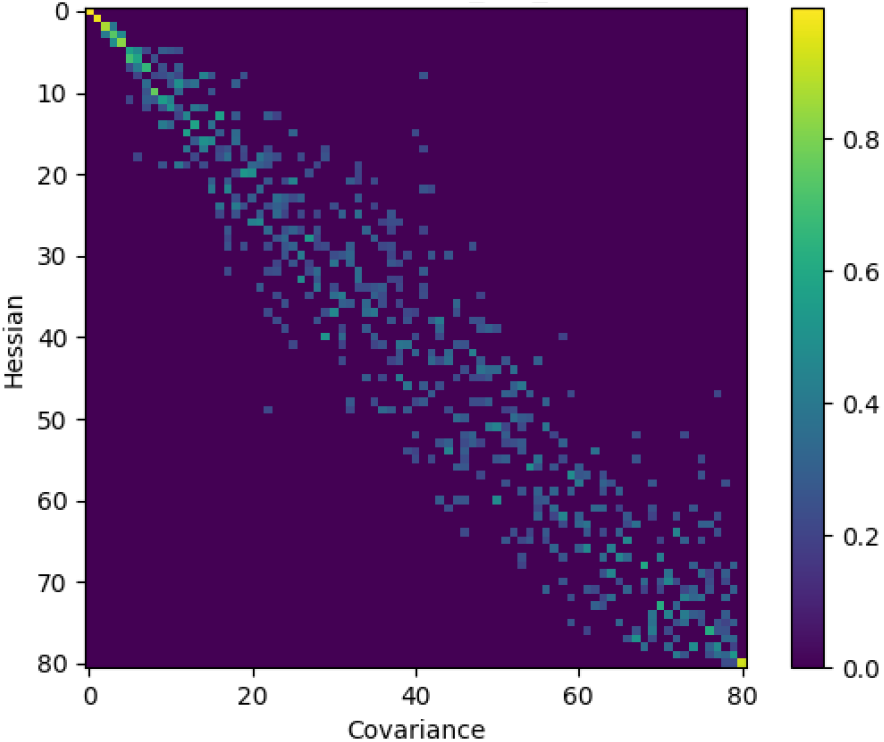
We compared the eigenvectors of the approximate Hessian with the principal components of the ensemble of parameter sets. The eigenvectors were sorted in descending order of their corresponding eigenvalue. The PCs were sorted in ascending order of their corresponding explained variance ratio (which is proportional to their eigenvalue). The heatmap shows the dot product of the eigenvectors and the PCs, with a brighter color implying a higher value.

### S2.7 Comparison between the eigenvectors of the Hessian and the inverse covariance matrix

We also compared the quality of the approximate Hessian derived from our iterative scheme, with the inverse of the covariance matrix (Σ^−1^) of the ensemble of parameter sets [95]. Since the eigenvectors of the Σ^−1^ matrix are identical to those of Σ, we obtained the principal components (PCs) of the ensemble of parameter sets (i.e. the eigenvectors of the covariance matrix). We then sorted the PCs in increasing order of the explained variance such that the PC with the smallest explained variance (proportional to its eigenvalue) received a rank of 1. Figure S2 (b) shows the parameters with substantial weights in these sorted eigenvectors. We note that the trends displayed by the PCs are qualitatively similar to those of the eigenvectors of the approximate Hessian in Figure S2 (b). We next examined if the eigenvectors and PCs were actually identical by studying their pairwise dot products. Figure S3 presents a heatmap of the pairwise dot products. A brighter color indicates a number closer to 1, indicating greater similarity. There is a very good correspondence between the first 8-10 eigenvectors and PCs, indicating that our iterative scheme is able to confidently estimate the stiffest directions in parameter space.

### S2.8 The relative ranges explored agree with the amount of data used to constrain the corresponding variable

Motivated by the success of our iterative Hessian-directed search in identifying the stiff directions of the cost function, we next examined the relationship between the stiff parameters and the data constraining the model. We first studied the relative ranges of parameter values explored for each parameter. These are visualized as ratios of parameter values with respect to **p**^*^ on a log10 scale in Figure S4. The parameters are sorted according to the relative ranges explored. The ensemble of parameter sets was further analyzed. The parameter ranges vary from< 1 to 2 orders of magnitude. Figure S4 summarizes these parameter ranges. One feature that stands out from this sorting is that the gamma parameters which govern the time scales are mostly found in the top half of the plot, with broad ranges. This is likely a consequence of the lack of time series data used to constrain most of the variables in the model. Examining the parameters with narrower ranges, at the bottom of the plot, we notice that while many parameters do occur in the equations of variables that are constrained by data, this is not the case for other parameters. Examples include *ω*_cyr_ (regulates basal dynamics Cyr1), *ω*_torc_ego_ (regulates the stimulation of TORC1 by Gtr1/2) and *ω*_gis_ (regulates basal dynamics of Gis1) (Figure S4) While the ranges in the figure indicate that the model is very sloppy in general, the occurrence of these ‘unconstrained’ parameters at the bottom of the figure was surprising. In order to investigate the influence these parameters had on the model, we decided to carry out a detailed parameter perturbation analysis.

### S2.9 Model structure exerts an important influence on the stiff/sloppy classification of parameters

In order to study the relationships among parameters, the model structure and the model constraints, we first ranked the parameters in the model by their contribution to the stiff directions. For this, an arbitrary cutoff of one standard deviation of the distribution of weights for parameters across all eigenvectors was chosen. Then, the parameters with absolute weights greater than the chosen cutoff were designated to contribute substantially to a given eigenvector. Finally, a cumulative list of parameters was constructed, where the rank of a given parameter is the eigenvector number in which it first appears with signficant weight.

From the comparison of the eigenvectors of the covariance matrix and the approximate Hessian, it can be observed that there is a good correspondence between the first 8 or 9 eigenvectors (Figure S3). These were designated the high confidence directions, and the parameters occurring in these directions are marked in bold in Table S3 respectively. A striking finding from this table is that among the top-ranked parameters, many are not constrained by data, i.e., they do not appear in equations whose dynamics are constrained by data.

While the ranks of parameters in Table S3 indicate a complex relationship between model constraints and model structure influencing the structure of the cost surface, we were interested in the distribution of parameters that do appear in equations constrained by data. To investigate this distribution, starting from **p**^*^ we picked each parameter one at a time and perturbed its value in a ±2.5%, ±10%, and ±10-fold range and obtained the fitting cost in each instance. We also measured the model cost when the parameter was set to 0. Figure S5 shows the results of this analysis. The color of the heatmap is the log 10 fold increase in cost over *C*_min_. The cost values are truncated to a 10-fold increase so that smaller costs are visible. We observe that while a perturbation of up to ±10% has little effect on model cost, a 10-fold change produces a dramatic increase in model cost for the parameters on top of the ranked list, while those at the bottom of the list show a decreased effect, in agreement with the ranking of stiffness. The arrows in Figure S5 indicate the parameters which occur in equations constrained by data. We observe that these parameters do not exhibit any type of clustering, and are distributed across the entire list.

**Figure S4:**
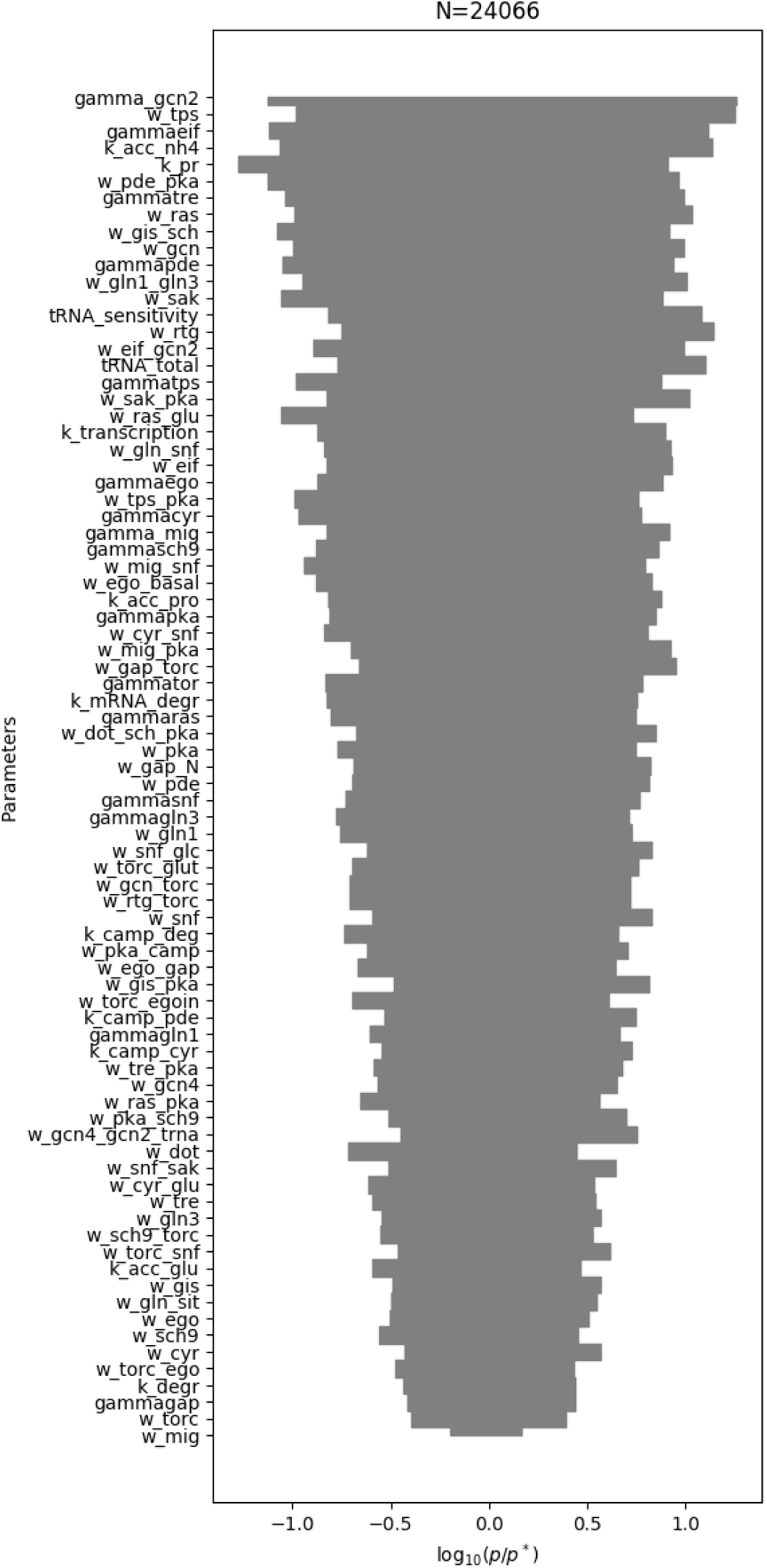
Ranges of parameters explored over all parameter sets at the end of four iterations. The smallest and largest value of each parameter over the ensemble were chosen, and the log10 value of the ratio with respect to the p* value was used to define the range.

These observations indicate that, despite a small amount of data available to constrain the model, the model structure (in particular the pathway crosstalk and feedback interactions) might play an important role in indirectly constraining other parts of the model that are not directly constrained by data.

**Figure S5:**
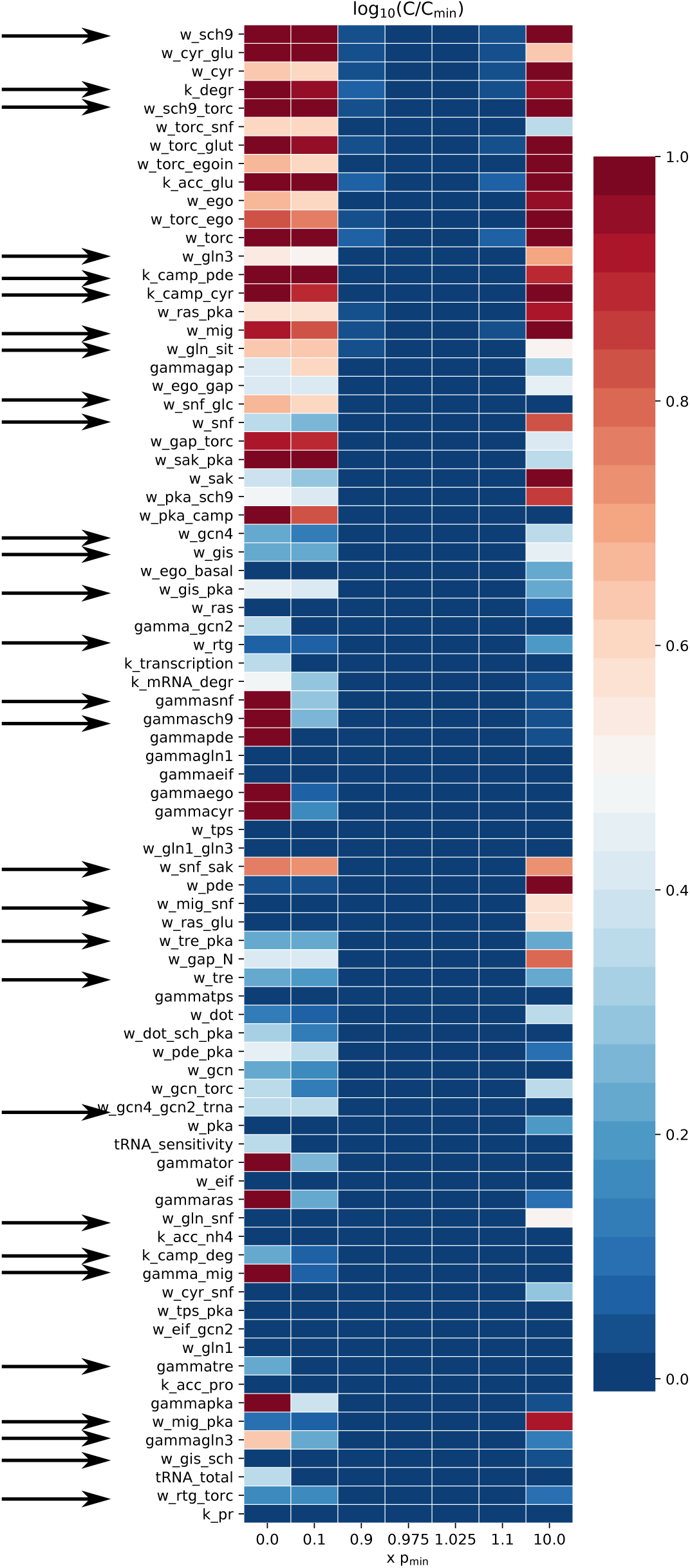
Parameter robustness is governed by model topology and experimental data.

## S3 Alternative interpretations of rapamycin treatment

As discussed in the section “Testing the model against observed phenotypes of mutant strains”, 22 experiments that we collected from the literature relate to rapamycin treatment in various mutant strains. There are three ways of interpreting the immediate effect of rapamycin on a strain grown in rich medium. First, via the Sch9 branch of TORC1 signaling, rapamycin can lead to an inhibition of ribosome biogenesis. In our model, this would be represented as an upregulation of Dot6 activity. Second, via the Tap42-Sit4 branch of TORC1 signaling, the activities of Gln3 and Gcn4 can result in an upregulation of nitrogen starvation and adaptation responses. Finally, via mechanisms not currently present in the model, TORC1 can directly or indirectly impinge on the cell cycle machinery to cause a G1 arrest [96]. We considered the first two definitions of the effect of rapamycin on cells. As shown in Figure S6, both the Dot6 and Gln3Gcn4 definitions show a trend of decreasing median cost until 11 of 22 experiments are explained. However, using the Dot6 definition, a large number of high cost parameter sets succeed in explaining up to 16 of 22 experiments.

We first describe the causes of model mismatch across both definitions of rapamycin. Finally, we investigate the basis of the 2000 parameter sets that explain up to 16 experiments using the Dot6 definition.

### Model mismatches

Eleven of the 22 experiments are explained by less than 50% of the parameter sets using either definition of rapamycin treatment (Table S1). Four of the 11 rapamycin treatment experiments involve strains carrying mutations downstream of TORC1. *SCH9*^*DE*^ encodes a constitutively active Sch9 kinase. *tap42-11* is a temperature senstitive allele of TAP42, which encodes a protein involved in transmitting the TORC1 signal to Sit4 and other stress response TFs. The single mutant strains *SCH9*^*DE*^ and *tap42-11* are slightly resistant to rapamycin treatment, whereas the double mutant strain *SCH9*^*DE*^*tap42-11* is fully resistant [97]. Using our definition of rapamycin treatment based on Gln3 and Gcn4 activities, strains involving *tap42-11* are predicted to be rapamycin resistant. However, since neither of these TFs are regulated by Sch9, the *SCH9*^*DE*^ strain is predicted to be rapamycin sensitive. Additionally, three strains, namely *gln3*Δ*gat1*Δ, *gcn4*Δ, and an overexpression mutant 2*μ* URE2 are all predicted to be rapamycin sensitive as a consequence of our chosen definition (Sections S4.6, S4.7 and S4.28).

Seven of the 11 rapamycin treatment experiments that we have curated include mutants of the carbon signaling pathway. These experiments clearly indicate that mutations affecting the carbon signaling pathways influence the nitrogen adaptation response, and the model’s failure to explain these results give us insight into the crosstalk between carbon and nitrogen pathways. As mentioned in the description of the PKA pathway in the Results section of the main text, our model supports some results from Schmelzle *et al*., but not from Zurita-Martinez *et al*., originating from strain specific differences. These observations account for three of the eight mismatches (‘14-bcy1’, ‘15-ira1’, and ‘16-ira1ira2’ described in Sections S4.14 to S4.16). Schmelzle *et al*. examined three hyperactivating PKA strains in a *gln3*Δ*gat1*Δ background, (‘19-RAS2v19gln3gat1’, ‘20-TPK1gln3gat1’, and ‘22-bcy1gln3gat’ described in Sections S4.19, S4.20 and S4.22). These strains were shown to be rapamycin resistant. However, our model predicts that these strains are sensitive to rapamycin. Our model does not currently include a direct interaction between PKA and TORC1. Similarly, the last rapamycin treatment mismatch relates to a *snf1*Δ strain which was observed to show rapamycin resistance [58]. While our model assumes that Snf1 inhibits TORC1, Snf1 will be inactive during growth on YPD, hence the model predicts that a *snf1* deletion will not affect Gln3 activity in this nutrient condition. Further mechanistic details of crosstalk between PKA, Snf1 and TORC1 will be needed in order to resolve these mismatches.

### The mechanism of Dot6 regulation explains the distrbution of number of experiments explained

We investigated the cause of the spread of number of experiments using the Dot6 definition, shown in Figure S6. We termed the parameter sets that explained 11 or fewer experiments as the “low” set, and those that explained more than 11 as the “high” set. We observed that the “low” set failed to explain the experiments starting from row 3 (‘20-TPK1 gln3 gat1’) in the first column of Table S1. These experiments constitute the model mismatches pertaining to claims by Schmelzle *et al*. and Zurita-Martinez *et al*. regarding the role of the PKA pathway in rescuing the rapamycin treament induced growth arrest phenotype. We hypothesized that the cause for this bimodal distibution of experiments explained was related to the representation of Dot6 in the model. Dot6 is regulated by both Sch9 and PKA. In order for rapamycin treatment to be sufficient for Dot6 activation, the Sch9 and PKA must regulate Dot6 via an AND gate. If this condition is not satisfied, the fact that rapamycin treament experiments are carried out in glucose medium will imply that PKA can independently repress Dot6. The parameters that regulate Dot6 are w_dot, the basal activation term, and w_dot_sch_pka, the strength of inhibition by the sum of PKA and Sch9 activities. Since PKA and Sch9 take a maximum value of 1.0, Dot6 is maximally repressed for any combination of parameters satisfying the following relation

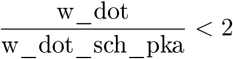

Figure S7 shows that qualitatively a majority of the parameter sets in the “high” set indeed satisfy this relationship.

**Figure S6:**
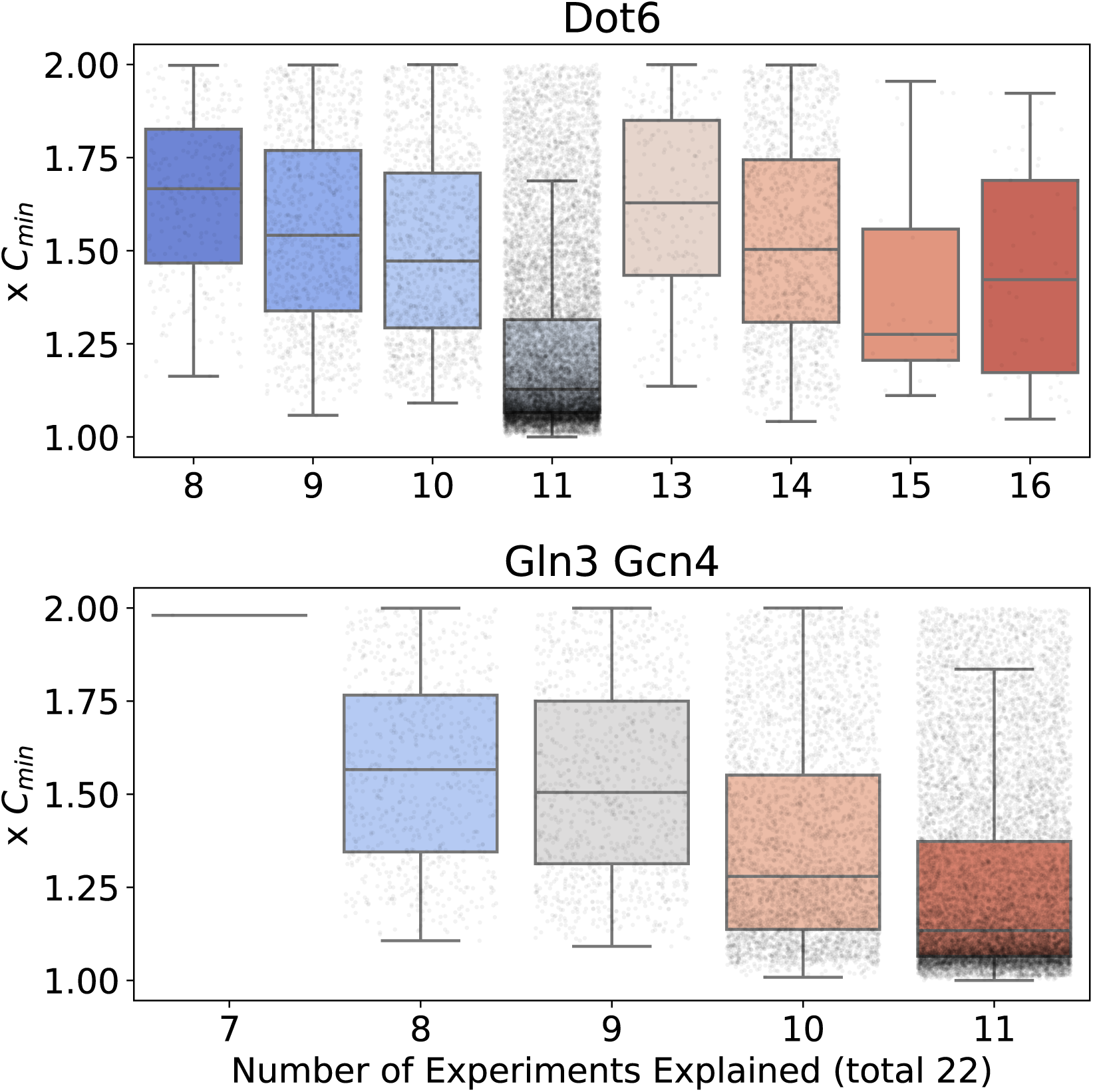
Dependence of model cost on explanatory capacity, across the entire collection of alternative sets of parameter values. Rapamycin experiments are defined using Dot6 as a model readout. The collection of parameter sets explaining less than or equal to 11 experiments are termed the “low” set, and those explaining more than 11 are termed the “high” set.

**Table S1:** List of rapamycin experiments, with confidence in model predictions. Confidence in qualitative predictions is expressed as the percentage of parameter sets that make the correct prediction. The *in silico* experiments corresponding to the experiment IDs are presented in Section S4.

| Exp ID | Dot6 | Gln3Gcn4 |
| --- | --- | --- |
| 20-TPK1 gln3 gat1 | 13.13 | 0.74 |
| 6-gln3 gat1 | 13.63 | 0.74 |
| 7-2μ URE2 | 13.63 | 0.7 |
| 37-tap42-11 | 13.63 | 71.41 |
| 34-snf1 | 13.63 | 0.0 |
| 27-gln3 gcn4 | 13.63 | 100.0 |
| 14-bcy1 | 13.64 | 0.0 |
| 22-bcy1 gln3 gat1 | 13.65 | 0.74 |
| 19-RAS2v19 gln3 gat1 | 13.65 | 0.74 |
| 16-ira1 ira2 | 13.65 | 0.0 |
| 15-ira1 | 13.65 | 0.0 |
| 10-gat1 | 85.44 | 84.81 |
| 28-gcn4 | 86.37 | 0.0 |
| 9-gln3 | 86.37 | 84.81 |
| 35-reg1 | 86.37 | 99.26 |
| 36-ure2 | 86.37 | 99.26 |
| 8-wt | 86.37 | 98.91 |
| 18-tpk1 | 86.67 | 99.06 |
| 17-ras2 | 86.78 | 99.17 |
| 21-bcy1 | 86.87 | 99.22 |
| 38-SCH9<sup>DE</sup> | 88.18 | 0.18 |
| 39-SCH9<sup>DE</sup> tap42-11 | 88.18 | 100.0 |

**Figure S7:**
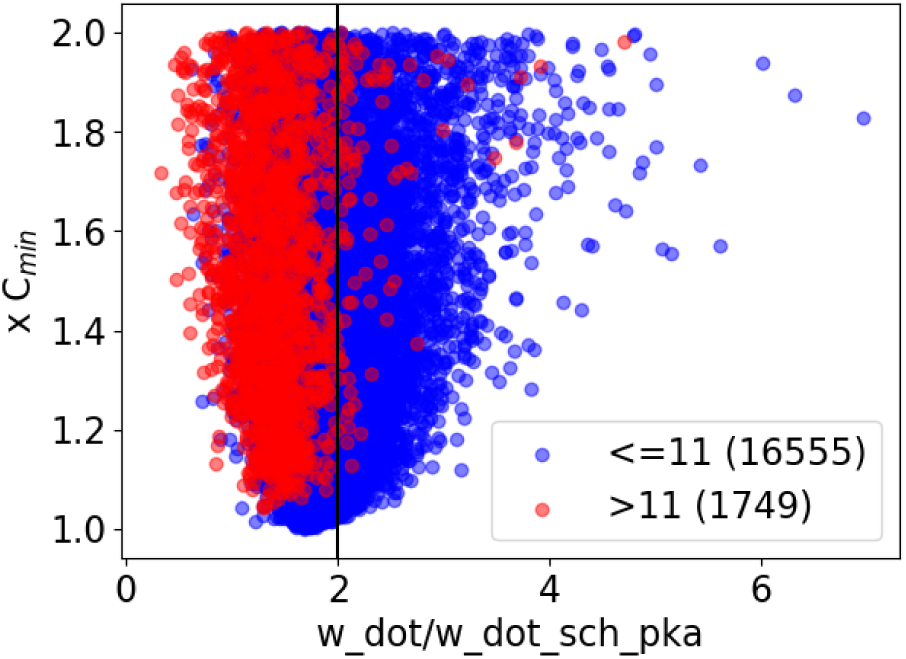
Distribution of ratios of kinetic parameters of Dot6. The “high” set is represented by the red points, and the “low” set is represented by the blue points. The scatter plot shows the ratio of the Dot6 parameters on the *x*-axis, and the cost associated with the parameter set on the *y*-axis

**Table S2:** Parameter values constituting the optimal parameter set, obtained from 10,000 steps of MCMC sampling. This set of values is used to define the simulation of a *wt* strain under HCHN conditions. Four time scale parameters describing the activation of transcription factors namely *γ*_*gcn*4_, *γ*_*rtg*13_, *γ*_*gis*1_, *γ*_*dot*6_ are set to 1, since there is no short time scale data available to constrain their dynamics.

| Name | Value | Name | Value | Name | Value |
| --- | --- | --- | --- | --- | --- |
| Cyr1_T | 1.00e+00 | k_degr | 8.98e-02 | w_gis | 1.30e+00 |
| Dot6_T | 1.00e+00 | k_mRNA_degr | 7.30e-02 | w_gis_pka | 3.30e+00 |
| EGO_T | 1.00e+00 | k_pr | 2.02e-02 | w_gis_sch | 8.42e-01 |
| EGOGAP_T | 1.00e+00 | k_transcription | 2.36e-01 | w_gln_sit | 8.61e-01 |
| Gcn2_T | 1.00e+00 | sigma_cyr | 3.50e+00 | w_gln_snf | 3.90e+00 |
| Gcn4_T | 1.00e+00 | sigma_dot | 2.00e+01 | w_gln1 | 2.22e-01 |
| Gis1_T | 1.00e+00 | sigma_ego | 5.00e+00 | w_gln1_gln3 | 5.20e-01 |
| Gln1_T | 1.00e+00 | sigma_eif | 1.00e+00 | w_gln3 | 6.39e-01 |
| Gln3_T | 1.00e+00 | sigma_gap | 1.00e+00 | w_mig | 1.06e+01 |
| Mig1_T | 1.00e+00 | sigma_gcn2 | 2.00e+01 | w_mig_pka | 2.31e+00 |
| PDE_T | 1.00e+00 | sigma_gcn4 | 5.00e+00 | w_mig_snf | 1.21e+00 |
| PKA_T | 1.00e+00 | sigma_gis1 | 1.00e+01 | w_pde | 3.83e-01 |
| Ras_T | 1.00e+00 | sigma_gln | 1.00e+01 | w_pde_pka | 2.89e+00 |
| Rtg13_T | 1.00e+00 | sigma_gln1 | 1.00e+00 | w_pka | 5.81e-02 |
| Sak_T | 1.00e+00 | sigma_mig1 | 2.70e-01 | w_pka_camp | 1.02e+02 |
| Sch9_T | 1.00e+00 | sigma_pde | 1.90e+00 | w_pka_sch9 | 1.75e+01 |
| Snf1_T | 1.00e+00 | sigma_pka | 1.00e+00 | w_ras | 2.08e-02 |
| TORC1_T | 1.00e+00 | sigma_ras | 1.00e+00 | w_ras_glu | 2.07e-01 |
| Tps1_T | 1.00e+00 | sigma_rtg | 1.00e+01 | w_ras_pka | 1.87e+00 |
| Trehalase_T | 1.00e+00 | sigma_sak | 2.00e+01 | w_rtg | 1.86e-01 |
| eIF_T | 1.00e+00 | sigma_sch9 | 8.00e+00 | w_rtg_torc | 8.77e-01 |
| gamma_gcn2 | 4.71e+00 | sigma_snf | 3.00e+00 | w_sak | 2.05e-01 |
| gamma_mig | 6.56e-01 | sigma_tor | 5.00e+00 | w_sak_pka | 3.75e-01 |
| gamma_cyr | 8.96e+00 | sigma_tps | 5.00e+00 | w_sch9 | 5.65e-01 |
| gamma_ego | 5.07e+01 | sigma_trehalase | 1.00e+01 | w_sch9_torc | 1.96e+00 |
| gamma_eif | 4.71e-01 | tRNA_scale | 7.45e+01 | w_snf | 5.38e-01 |
| gamma_gap | 5.62e-01 | tRNA_total | 2.47e+00 | w_snf_glc | 1.15e+00 |
| gamma_gln1 | 6.35e-02 | w_cyr | 1.35e+00 | w_snf_sak | 1.52e+00 |
| gamma_gln3 | 8.09e-02 | w_cyr_glu | 5.13e+00 | w_torc | 5.39e-01 |
| gamma_pde | 2.82e-01 | w_cyr_snf | 1.19e-01 | w_torc_ego | 8.77e-01 |
| gamma_pka | 2.68e+00 | w_dot | 2.93e-01 | w_torc_egoin | 3.03e-01 |
| gamma_ras | 1.82e+00 | w_dot_sch_pka | 1.63e-01 | w_torc_glut | 8.63e-01 |
| gamma_sch9 | 4.63e+00 | w_ego | 2.84e-01 | w_torc_snf | 4.37e-01 |
| gamma_snf | 8.20e-01 | w_ego_basal | 1.10e-02 | w_tps | 5.30e-02 |
| gamma_tor | 7.55e+00 | w_ego_gap | 2.21e+00 | w_tps_pka | 5.74e-01 |
| gamma_tps | 4.71e-01 | w_eif | 3.73e+00 | w_tre | 1.07e+00 |
| gamma_tre | 3.42e-01 | w_eif_gcn2 | 2.76e-01 | w_tre_pka | 3.07e+00 |
| k_acc_glu | 4.92e-02 | w_gap_N | 7.76e+00 | ATP | 1.0 |
| k_acc_nh4 | 1.47e-03 | w_gap_torc | 8.83e+01 | Carbon | 1.0 |
| k_acc_pro | 2.15e-04 | w_gcn | 1.15e-01 | Glutamine_ext | 1.0 |
| k_camp_cyr | 1.09e+01 | w_gcn_torc | 1.29e+00 | NH4 | 0.00e+00 |
| k_camp_deg | 8.38e-02 | w_gcn4 | 7.43e-01 | Proline | 0.00e+00 |
| k_camp_pde | 1.41e+01 | w_gcn4_gcn2_trna | 1.53e+00 |  |  |

**Table S3:**
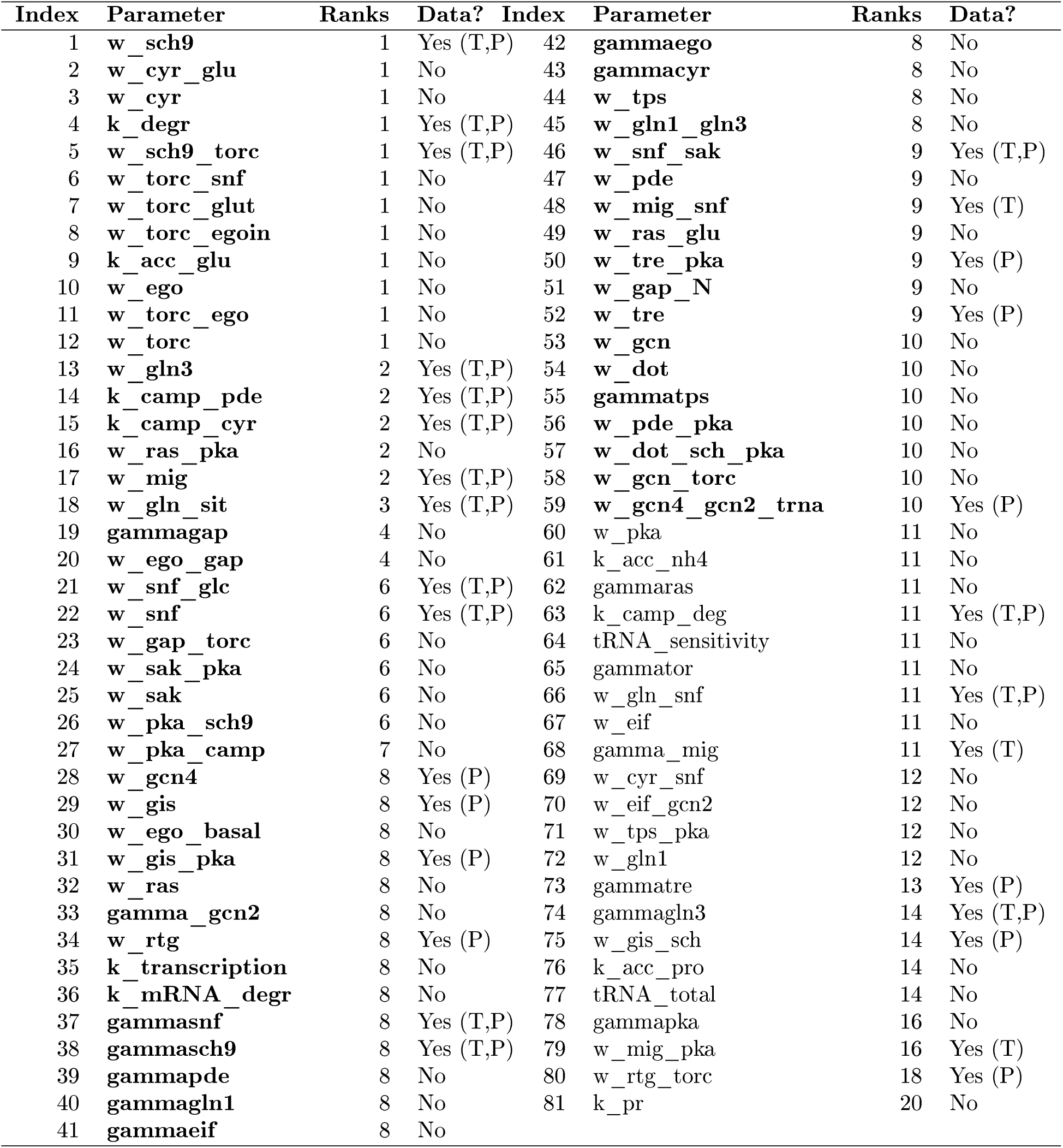
Ranked list of parameters. The ‘Data?’ column indicates whether or not the equation in which a parameter appears is contrained by any type of data. If it is constrained, the column further indicates the type of data, ‘T’ for timecourses or ‘P’ for perturbations. The parameters highlighted in bold indicate those appearing in the first 10 stiff directions.

## S4 Comparison of model predictions with qualitative experimental data

This section records the curated qualitative experiments, along with the model simulations using the optimal parameter set. Each experiment has a unique experiment ID, a table containing the interpreted and predicted states of transcription factors, along with the simulated values at steady state, a description of the experiment containing a reference to the original publication, the strain used, and the experiment performed, and the parameters used to represent the strain and the shift experiment in the model. Finally, the simulated time courses of the six model readouts and the interpretation of the simulation are described.

As described in Section S3, we use two definitions of rapamycin experiments. The sample simulations here use the Gln3Gcn4 definition of rapamycin treatment. Experiments involving rapamycin treatment are thus indicated using the text **Readout used is Gln3 Gcn4**. For the general trend of predictions using the alternate definition using Dot6, please see Table S1. (Note that the results shown here only correspond to one parameter set of our collection of 18,000 alternate sets of parameter values.)

### S4.1 1-rho0

| TF | Interpreted | Simulated | Simulation |
| --- | --- | --- | --- |
| *Gis1* | - | - | 0.000 |
| *Mig1* | - | - | 1.487 |
| *Dot6* | - | - | 0.312 |
| *Gcn4* | - | - | 0.024 |
| **Rtg13** | Off | Off | 0.274 |
| *Gln3* | - | - | 0.983 |

#### Description

Liu et al, 1999 studied a *rho0* strain (PSY142 *ρ*^0^) grown in YP + 5% glucose.

#### Representation

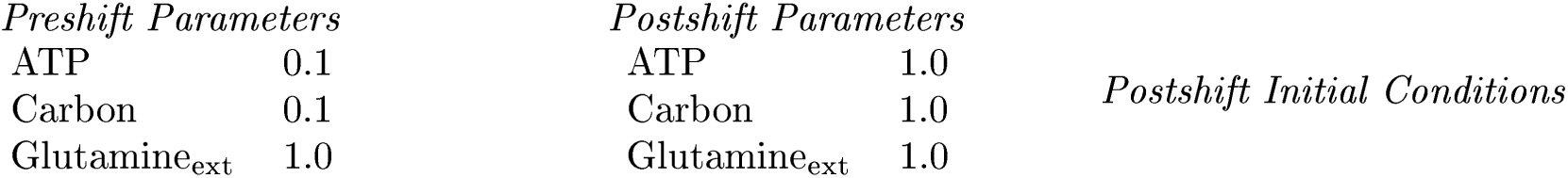

**Representation:**
| *Preshift Parameters* |  | *Postshift Parameters* |  | *Postshift Initial Conditions* |
| --- | --- | --- | --- | --- |
| ATP | 0.1 | ATP | 1.0 |  |
| Carbon | 0.1 | Carbon | 1.0 |  |
| Glutamine<sub>ext</sub> | 1.0 | Glutamine<sub>ext</sub> | 1.0 |  |

#### Mutant definition

| *Parameters* |  | *Initial conditions* |
| --- | --- | --- |
| k<sub>accglu</sub> | 0.0369 |  |
| k<sub>accpro</sub> | 0.0002 |  |
| k<sub>accnh4</sub> | 0.0011 |  |

#### Model agrees with experiment

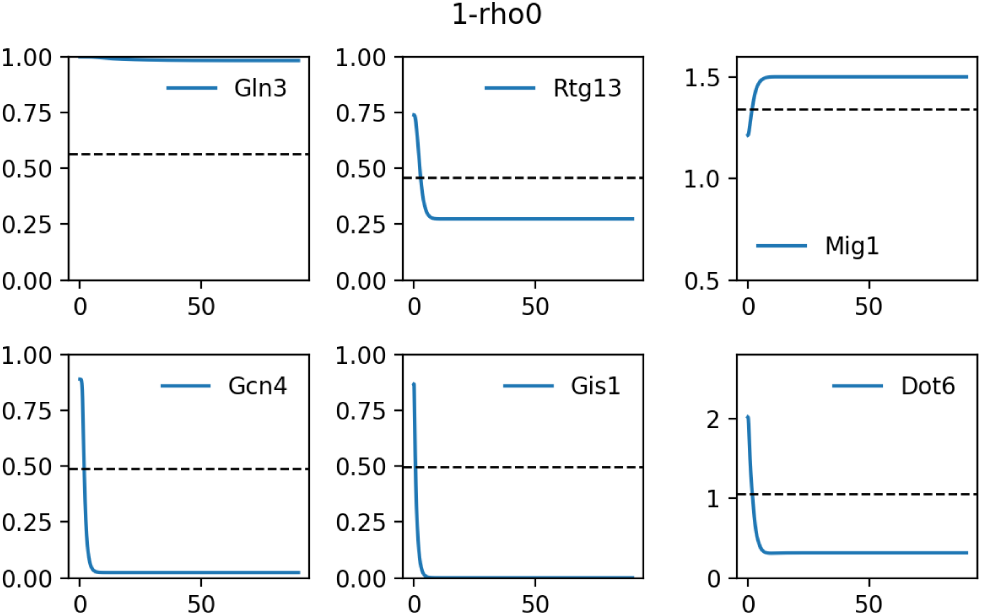

### S4.2 2-rho0

| TF | Interpreted | Simulated | Simulation |
| --- | --- | --- | --- |
| *Gis1* | - | - | 0.000 |
| *Mig1* | - | - | 1.482 |
| *Dot6* | - | - | 1.121 |
| *Gcn4* | - | - | 0.722 |
| **Rtg13** | On | On | 0.683 |
| *Gln3* | - | - | 0.999 |

#### Description

Liu et al, 1999 studied a *rho0* strain (PSY142 *ρ*^0^) grown in YP + 2% raffinose.

#### Representation

**Representation:**
| *Preshift Parameters* |  | *Postshift Parameters* |  | *Postshift Initial Conditions* |
| --- | --- | --- | --- | --- |
| ATP | 0.1 | ATP | 0.7 |  |
| Carbon | 0.1 | Carbon | 0.7 |  |
| Glutamine<sub>ext</sub> | 1.0 | Glutamine<sub>ext</sub> | 0.5 |  |

#### Mutant definition

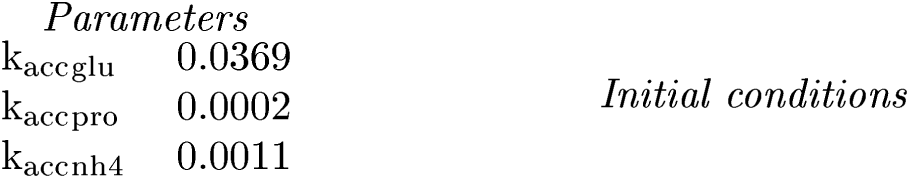

#### Model agrees with experiment

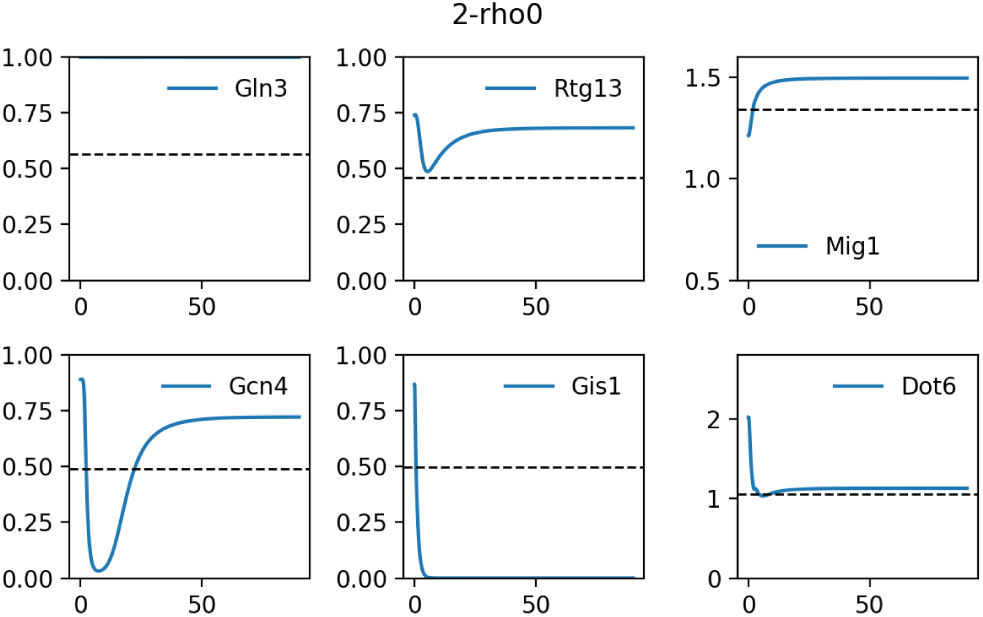

### S4.3 3-rtg1

| TF | Interpreted | Simulated | Simulation |
| --- | --- | --- | --- |
| *Gis1* | - | - | 0.000 |
| *Mig1* | - | - | 1.498 |
| *Dot6* | - | - | 0.532 |
| *Gcn4* | - | - | 0.024 |
| **Rtg13** | Off | Off | 0.279 |
| *Gln3* | - | - | 0.992 |

#### Description

Liu et al, 1999 studied a *rtg1* strain (PSY142 *ρ*^0^) grown in YNBD + 0.02% Glutamate.

#### Representation

**Representation:**
| *Preshift Parameters* |  | *Postshift Parameters* |  | *Postshift Initial Conditions* |
| --- | --- | --- | --- | --- |
| ATP | 1.0 | ATP | 1.0 |  |
| Carbon | 1.0 | Carbon | 1.0 |  |
| Glutamine<sub>ext</sub> | 1.0 | Glutamine<sub>ext</sub> | 0.9 |  |

#### Mutant definition

| *Parameters* |  |
| --- | --- |
| Rtg13<sub>T</sub> | 0.75 |
| k<sub>accglu</sub> | 0.0369 |
| k<sub>accpro</sub> | 0.0002 |
| k<sub>accnh4</sub> | 0.0011 |

#### Model agrees with experiment

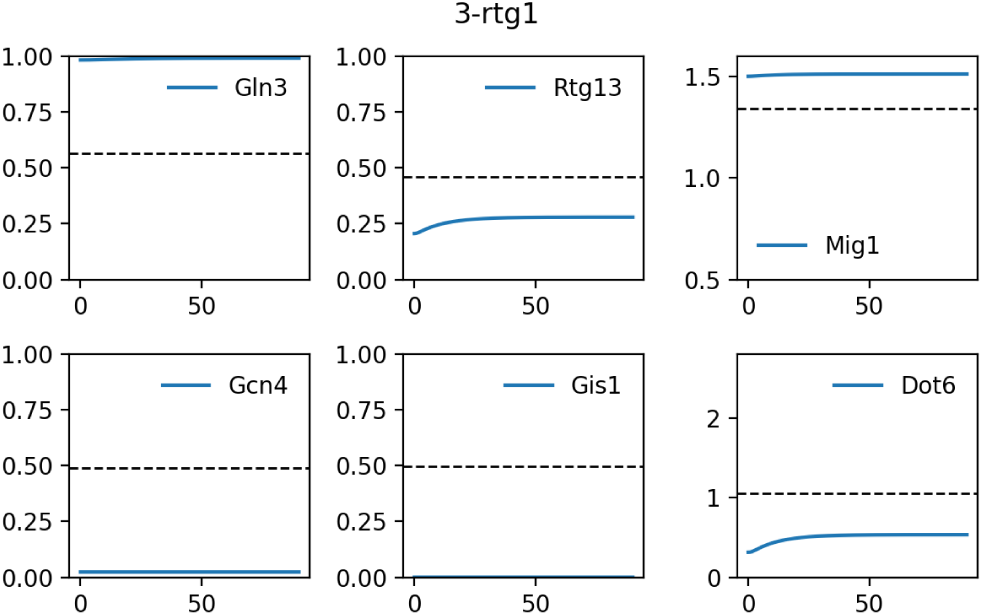

### S4.4 4-rtg1

| TF | Interpreted | Simulated | Simulation |
| --- | --- | --- | --- |
| *Gis1* | - | - | 0.000 |
| *Mig1* | - | - | 1.503 |
| *Dot6* | - | - | 1.094 |
| *Gcn4* | - | - | 0.945 |
| **Rtg13** | Off | On | 0.613 |
| *Gln3* | - | - | 1.000 |

#### Description

Liu et al, 1999 studied a *rtg1* strain (PSY142 *ρ*^0^) grown in YNBD.

#### Representation

**Representation:**
| *Preshift Parameters* |  | *Postshift Parameters* |  | *Postshift Initial Conditions* |
| --- | --- | --- | --- | --- |
| ATP | 1.0 | ATP | 1.0 |  |
| Carbon | 1.0 | Carbon | 1.0 |  |
| Glutamine<sub>ext</sub> | 1.0 | Glutamine<sub>ext</sub> | 0.0 |  |

#### Mutant definition

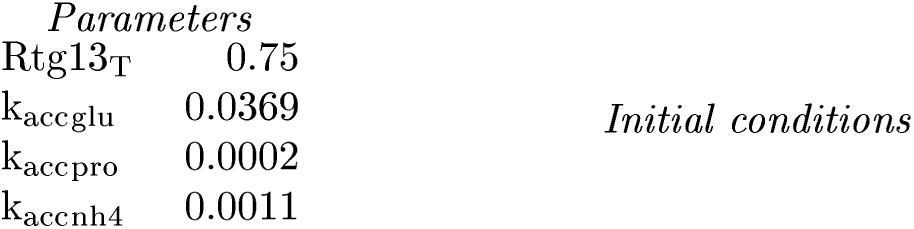

#### Model does not agree with experiment

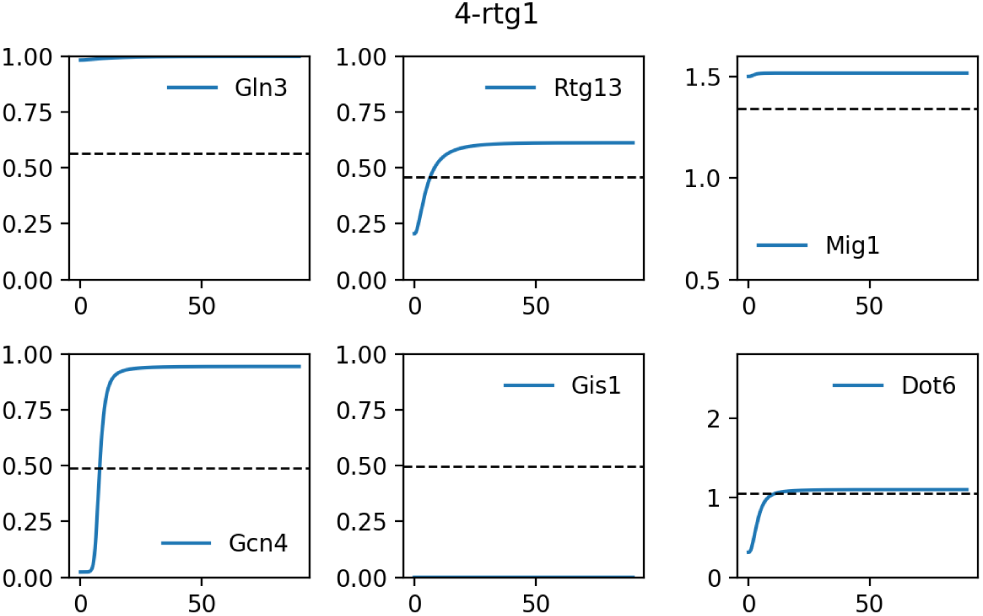

### S4.5 5-snf1

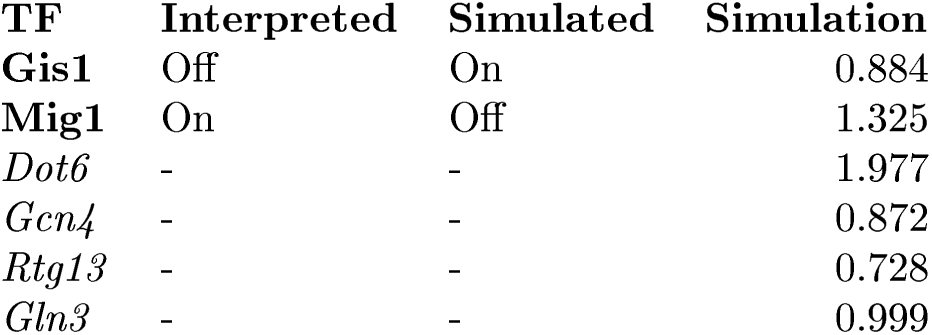

#### Description

Gasmi et al, 2014 studied a *snf1* strain (BY4741) grown in Minimal + (0.2%casa) + 2% Ethanol.

#### Representation

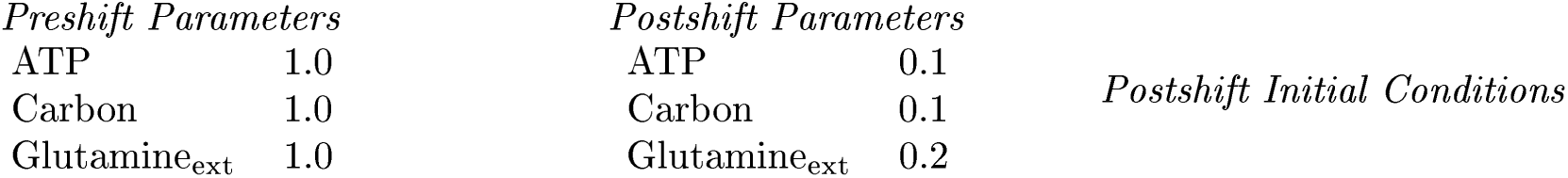

*Preshift Parameters*
|  |  |
| --- | --- |
| ATP | 1.0 |
| Carbon | 1.0 |
| Glutamine<sub>ext</sub> | 1.0 |
| *Postshift Parameters* |  |
| ATP | 0.1 |
| Carbon | 0.1 |
| Glutamine<sub>ext</sub> | 0.2 |

#### Mutant definition

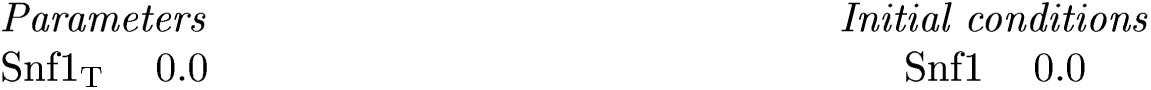

*Parameters*
|  |  |
| --- | --- |
| Snf1<sub>T</sub> | 0.0 |
| *Initial conditions* |  |
| Snf1 | 0.0 |

#### Model does not agree with experiment

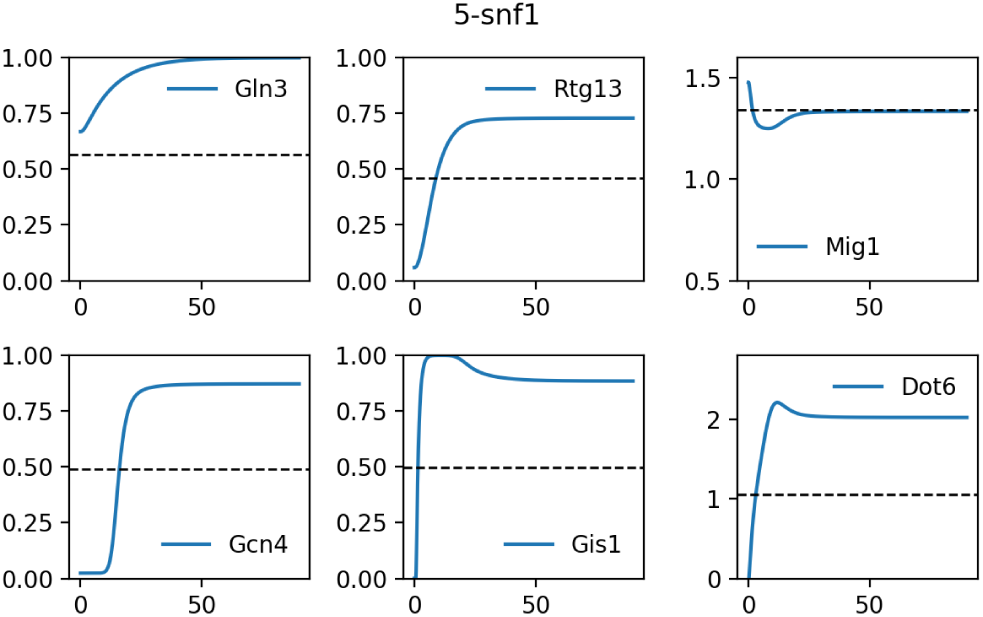

### S4.6 6-gln3 gat1

#### Readout used is Gln3 Gcn4

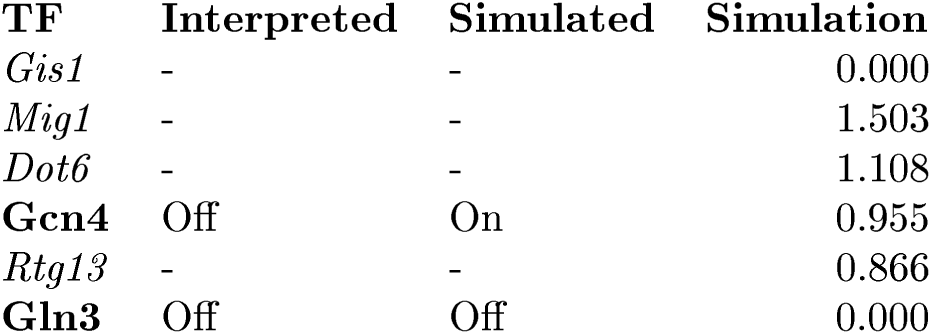

##### Description

Beck et al, 1999 studied a *gln3 gat1* strain (wt) grown in YPD + rapamycin.

##### Representation

**Representation:**
| *Preshift Parameters* |  | *Postshift Parameters* |  | *Postshift Initial Conditions* |  |
| --- | --- | --- | --- | --- | --- |
| ATP | 1.0 | ATP | 1.0 |  |  |
| Carbon | 1.0 | Carbon | 1.0 |  |  |
| Glutamine<sub>ext</sub> | 1.0 | Glutamine<sub>ext</sub> | 1.0 | TORC1 | 0 |
|  |  | TORC1<sub>T</sub> | 0.0 |  |  |

##### Mutant definition

| *Parameters* | *Initial conditions* |
| --- | --- |
| Gln3<sub>T</sub> 0.0 | Gln3 0.0 |

##### Model does not agree with experiment

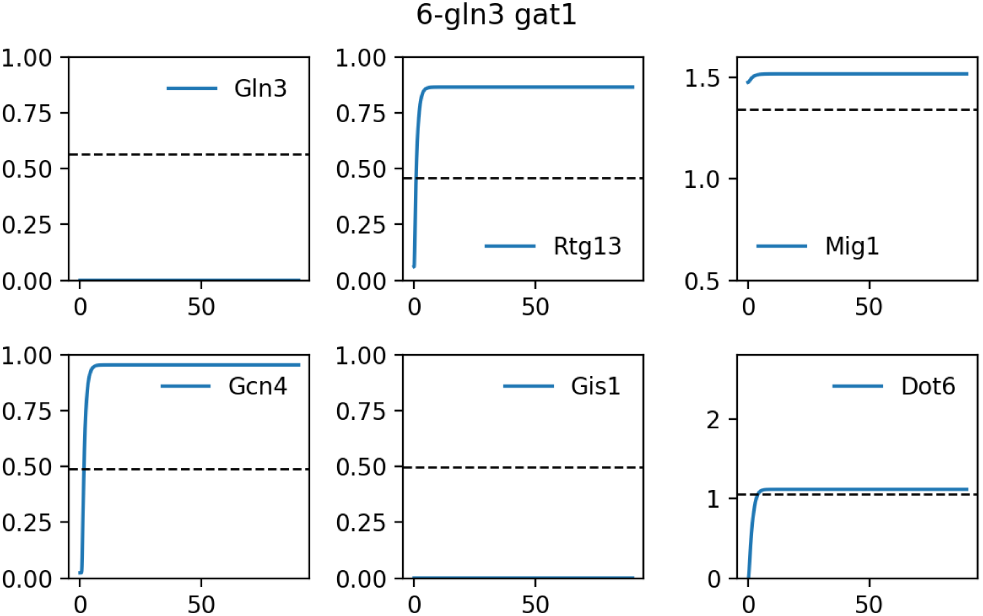

### S4.7 7-2*µ* URE2

#### Readout used is Gln3 Gcn4

| TF | Interpreted | Simulated | Simulation |
| --- | --- | --- | --- |
| *Gis1* | - | - | 0.000 |
| *Mig1* | - | - | 1.503 |
| *Dot6* | - | - | 1.108 |
| **Gcn4** | Off | On | 0.955 |
| *Rtg13* | - | - | 0.866 |
| **Gln3** | Off | On | 0.999 |

##### Description

Beck et al, 1999 studied a *2µ URE2* strain (wt) grown in YPD + rapamycin.

##### Representation

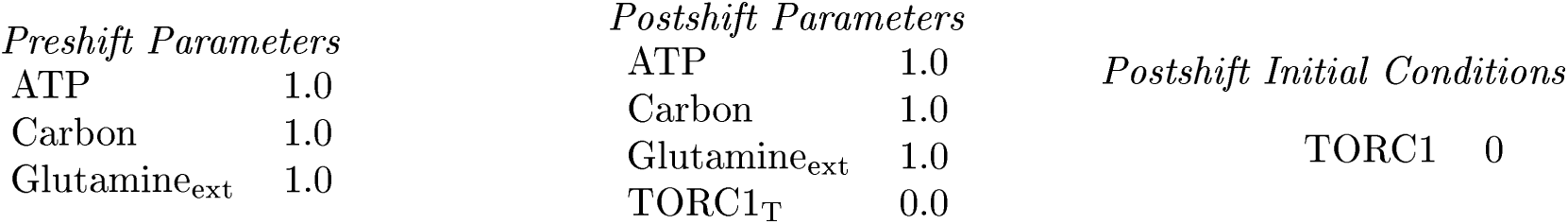

##### Mutant definition

| *Parameters* | *Initial conditions* |
| --- | --- |
| w<sub>gln3</sub> 1.2784 |  |

##### Model does not agree with experiment

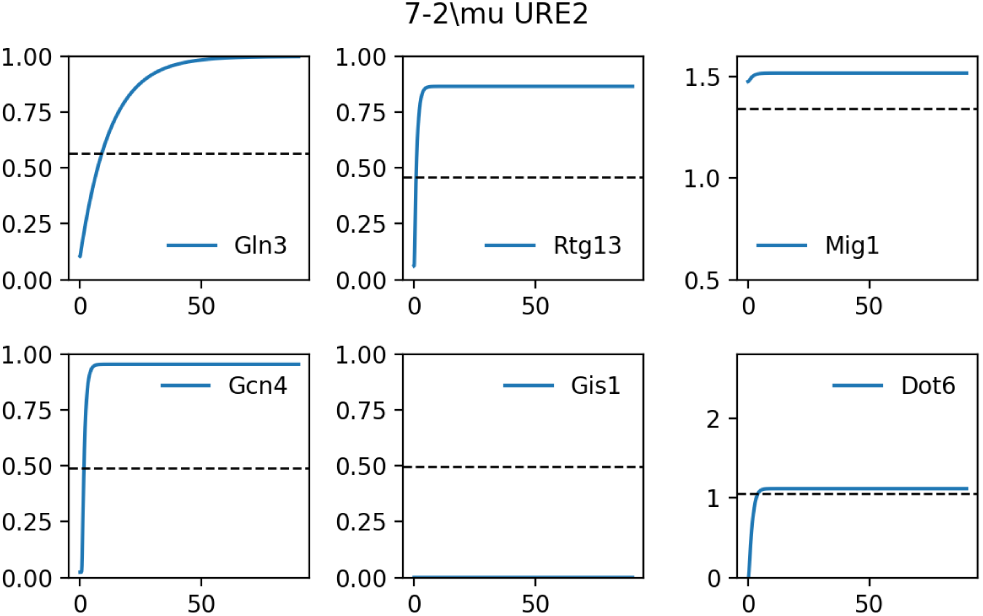

### S4.8 8-wt

#### Readout used is Gln3 Gcn4

| TF | Interpreted | Simulated | Simulation |
| --- | --- | --- | --- |
| *Gis1* | - | - | 0.000 |
| *Mig1* | - | - | 1.503 |
| *Dot6* | - | - | 1.108 |
| **Gcn4** | On | On | 0.955 |
| *Rtg13* | - | - | 0.866 |
| **Gln3** | On | On | 1.000 |

##### Description

Beck et al, 1999 studied a *wt* strain (wt) grown in YPD + rapamycin.

##### Representation

|  |  |
| --- | --- |
| ATP | 1.0 |
| Carbon | 1.0 |
| Glutamine<sub>ext</sub> | 1.0 |
| TORC1<sub>T</sub> | 0.0 |

##### Mutant definition

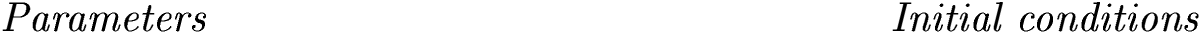

##### Model agrees with experiment

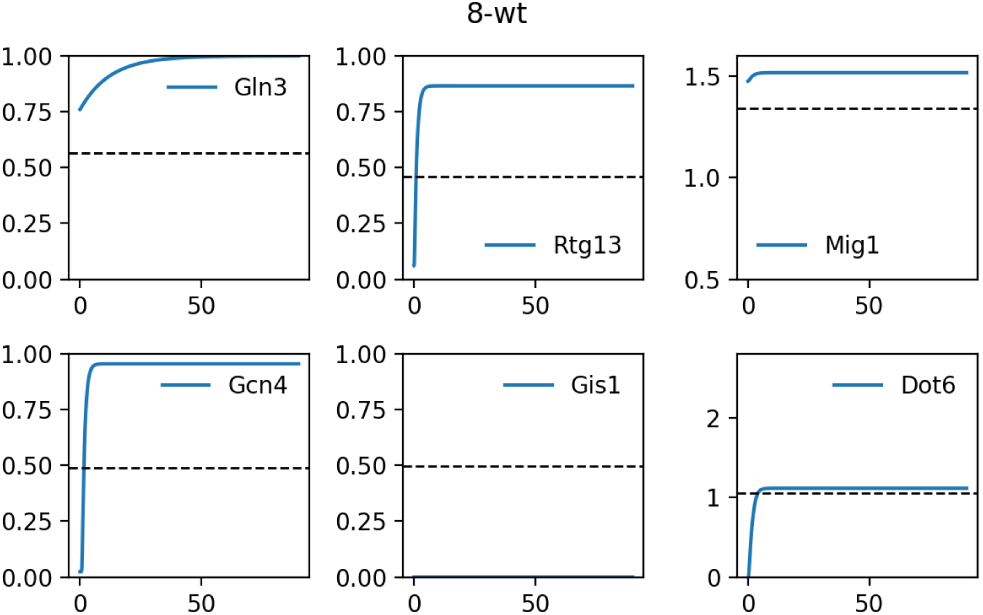

### S4.9 9-gln3

#### Readout used is Gln3 Gcn4

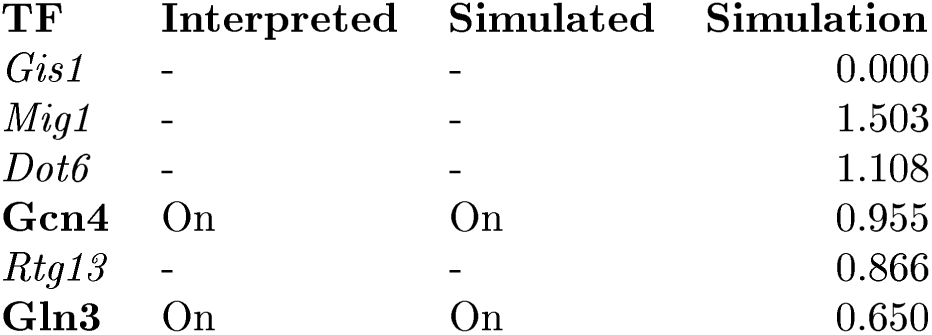

##### Description

Beck et al, 1999 studied a *gln3* strain (wt) grown in YPD + rapamycin.

##### Representation

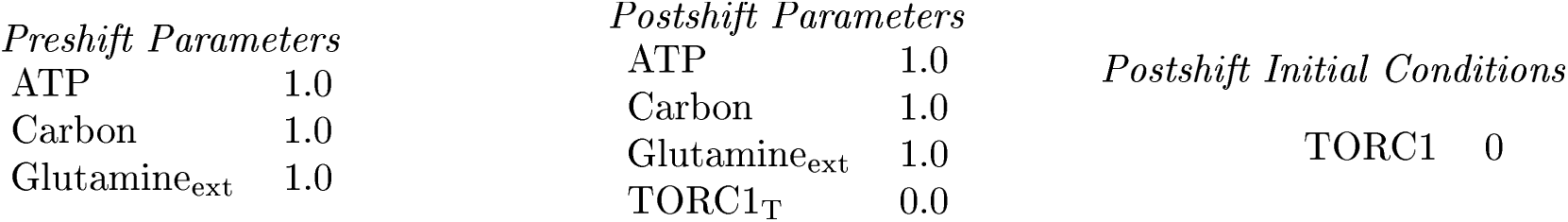

##### Mutant definition

| *Parameters* | *Initial conditions* |
| --- | --- |
| Gln3<sub>T</sub> 0.65 | Gln3 0.0 |

##### Model agrees with experiment

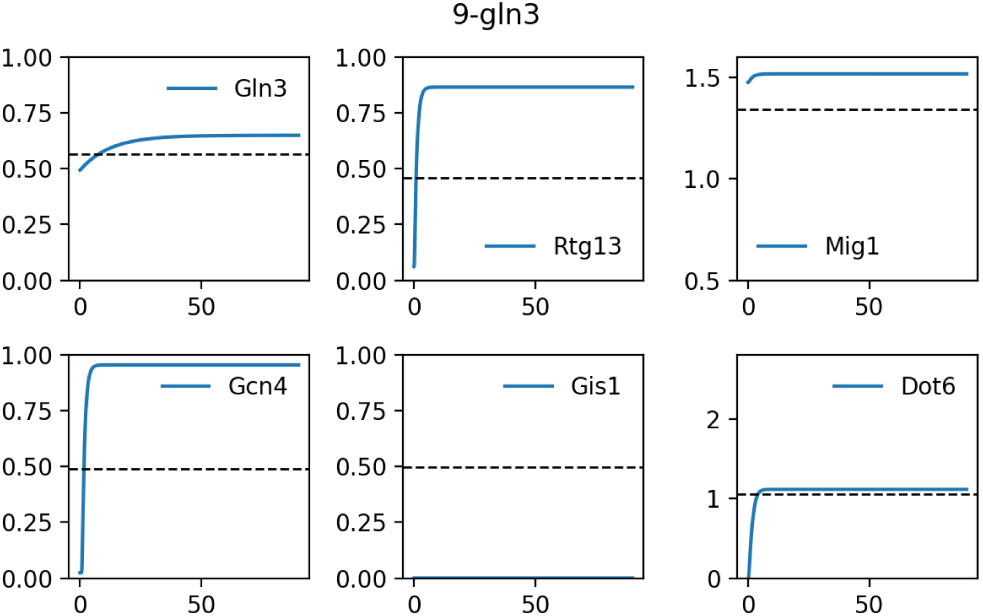

### S4.10 10-gat1

#### Readout used is Gln3 Gcn4

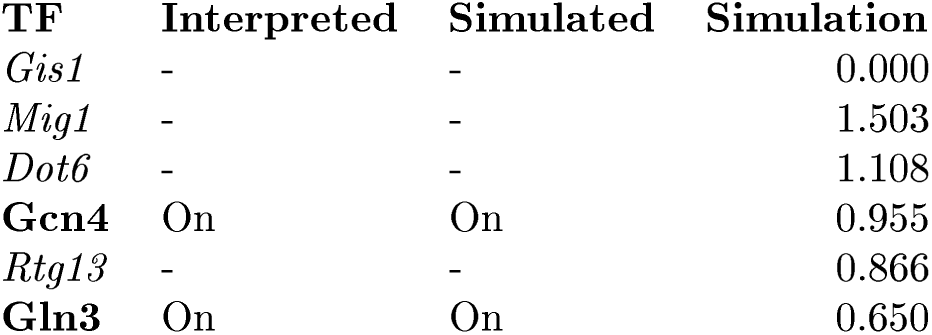

##### Description

Beck et al, 1999 studied a *gat1* strain (wt) grown in YPD + rapamycin.

##### Representation

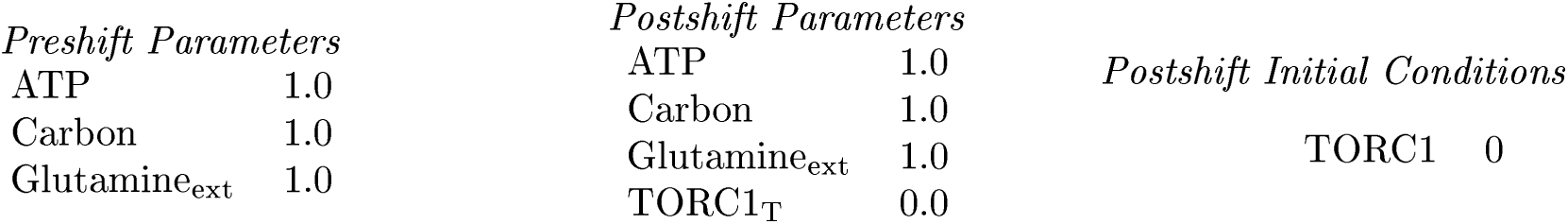

##### Mutant definition

| *Parameters* | *Initial conditions* |
| --- | --- |
| Gln3<sub>T</sub> 0.65 | Gln3 0.0 |

##### Model agrees with experiment

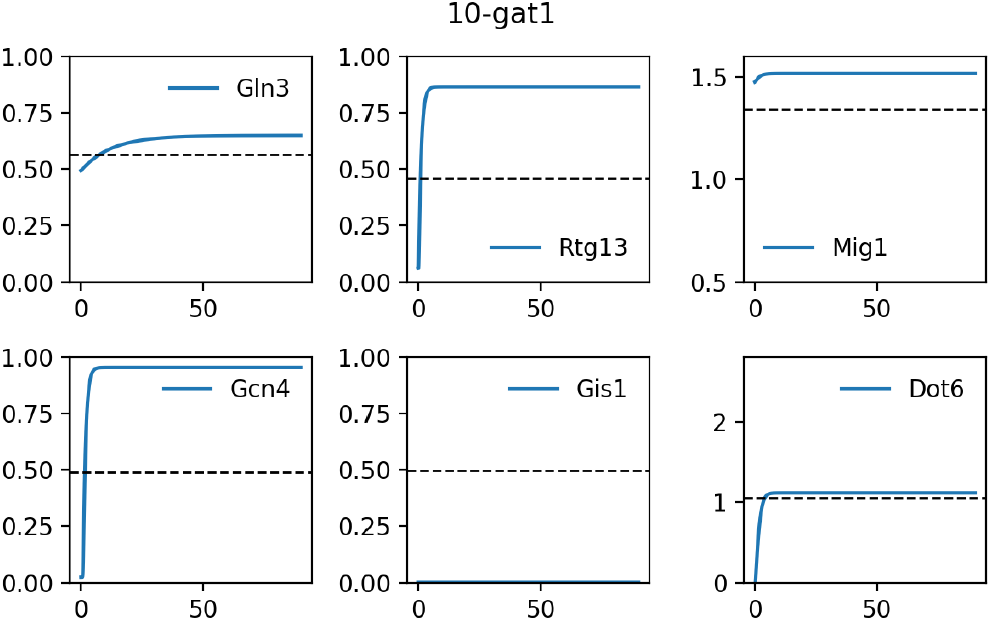

### S4.11 11-gln3

| TF | Interpreted | Simulated | Simulation |
| --- | --- | --- | --- |
| *Gis1* | - | - | 0.000 |
| *Mig1* | - | - | 1.503 |
| *Dot6* | - | - | 1.037 |
| *Gcn4* | - | - | 0.806 |
| *Rtg13* | - | - | 0.702 |
| **Gln3** | Off | On | 0.649 |

#### Description

Crespo et al, 2002 studied a *gln3* strain (TB123) grown in SD + 1mM MSX.

#### Representation

|  |  |
| --- | --- |
| ATP | 1.0 |
| Carbon | 1.0 |
| Glutamine<sub>ext</sub> | 1.0 |
| k<sub>accglu</sub> | 0.0 |

*Postshift Initial Conditions*

#### Mutant definition

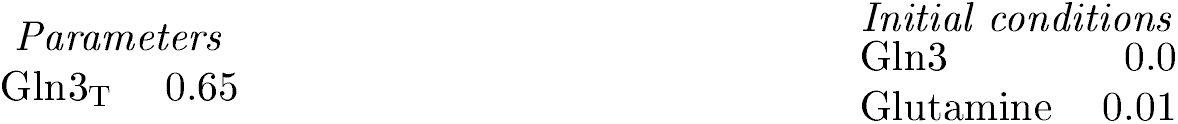

|  |  |
| --- | --- |
| *Parameters* |  |
| Gln3<sub>T</sub> | 0.65 |
| Gln3 | 0.0 |
| Glutamine | 0.01 |

#### Model does not agree with experiment

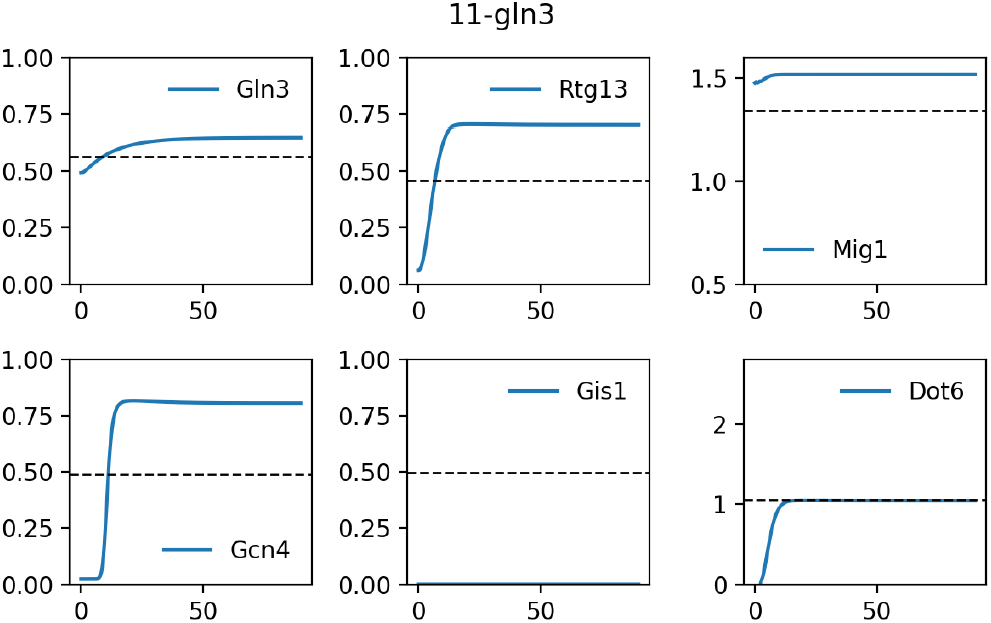

### S4.12 12-gln3 gat1

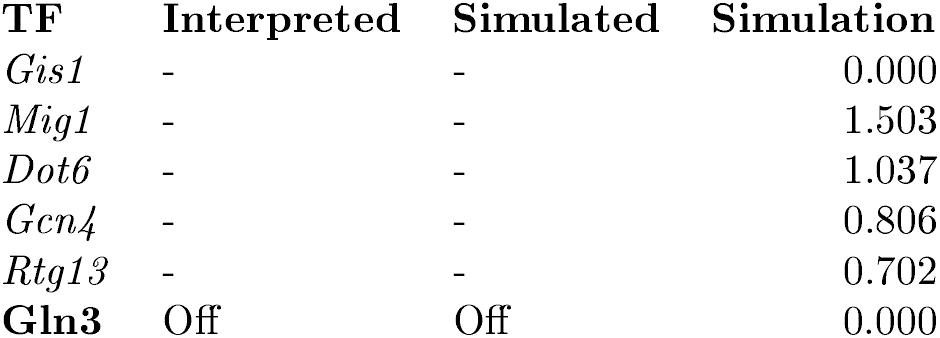

#### Description

Crespo et al, 2002 studied a *gln3 gat1* strain (TB123) grown in SD + 1mM MSX.

#### Representation

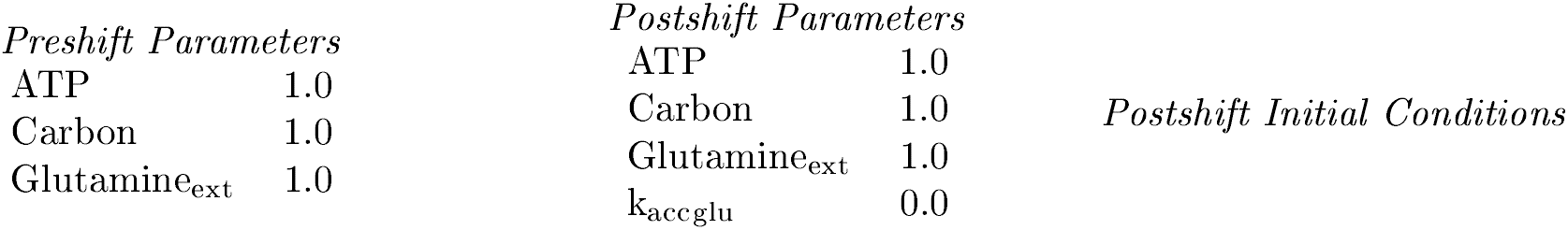

#### Mutant definition

| *Parameters* |  | *Initial conditions* |  |
| --- | --- | --- | --- |
| Gln3<sub>T</sub> | 0.0 | Gln3 | 0.0 |
|  |  | Glutamine | 0.01 |

#### Model agrees with experiment

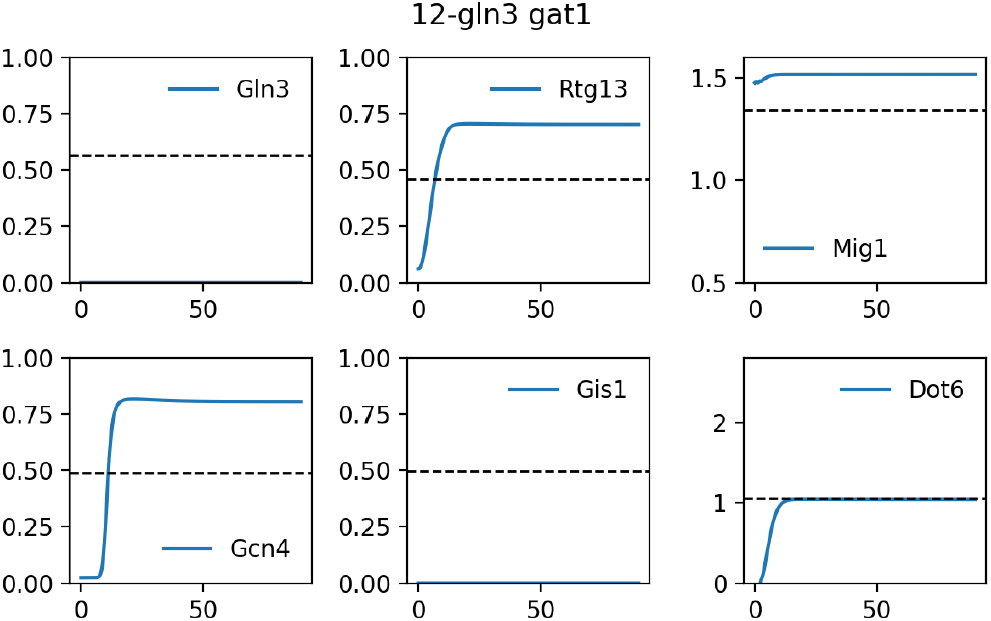

### S4.13 13-wt

| TF | Interpreted | Simulated | Simulation |
| --- | --- | --- | --- |
| *Gis1* | - | - | 0.000 |
| *Mig1* | - | - | 1.503 |
| *Dot6* | - | - | 1.037 |
| *Gcn4* | - | - | 0.806 |
| *Rtg13* | - | - | 0.702 |
| **Gln3** | On | On | 0.999 |

#### Description

Crespo et al, 2002 studied a *wt* strain (TB123) grown in SD + 1mM MSX.

#### Representation

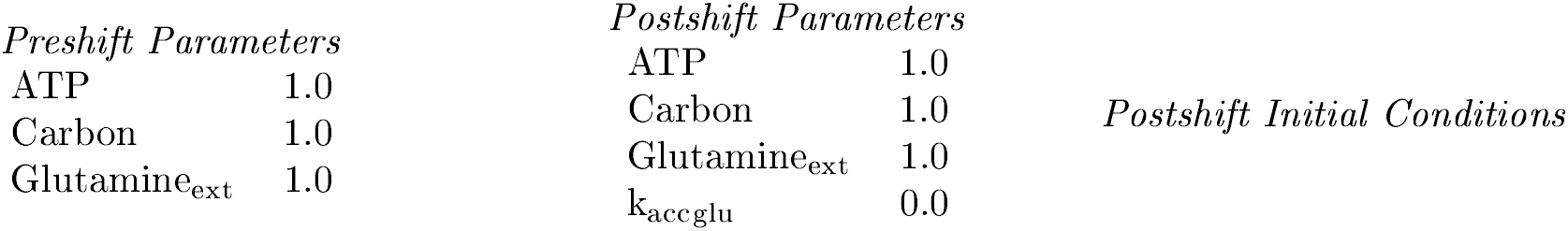

#### Mutant definition

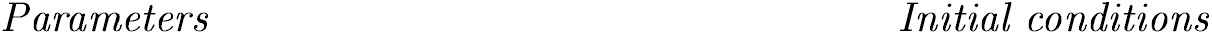

#### Model agrees with experiment

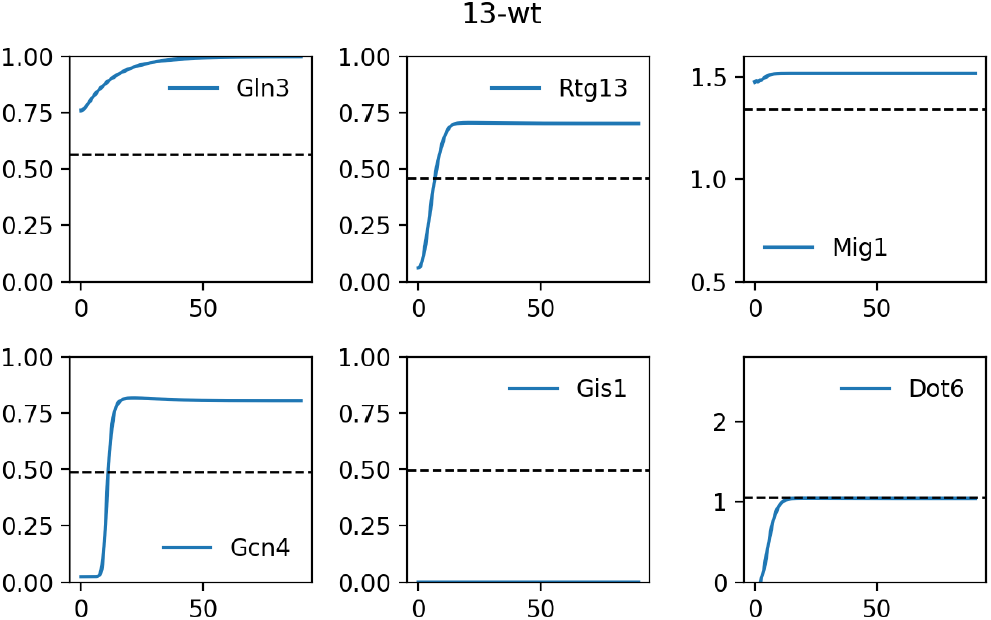

### S4.14 14-bcy1

#### Readout used is Gln3 Gcn4

| TF | Interpreted | Simulated | Simulation |
| --- | --- | --- | --- |
| *Gis1* | - | - | 0.000 |
| *Mig1* | - | - | 1.503 |
| *Dot6* | - | - | 1.108 |
| **Gcn4** | Off | On | 0.955 |
| *Rtg13* | - | - | 0.866 |
| **Gln3** | Off | On | 1.000 |

##### Description

Zurita-Martinez et al, 2005 studied a *bcy1* strain (S1278b) grown in YP Glucose + 50nM rapamycin.

##### Representation

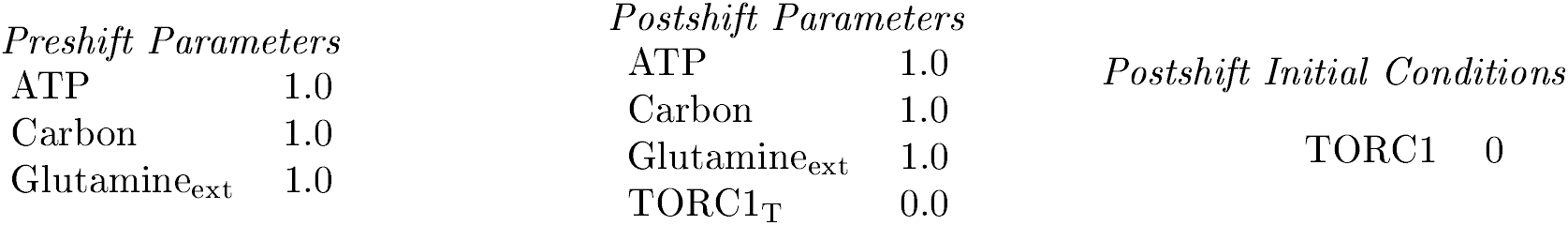

##### Mutant definition

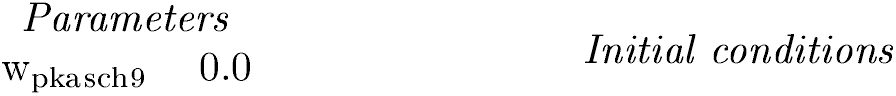

##### Model does not agree with experiment

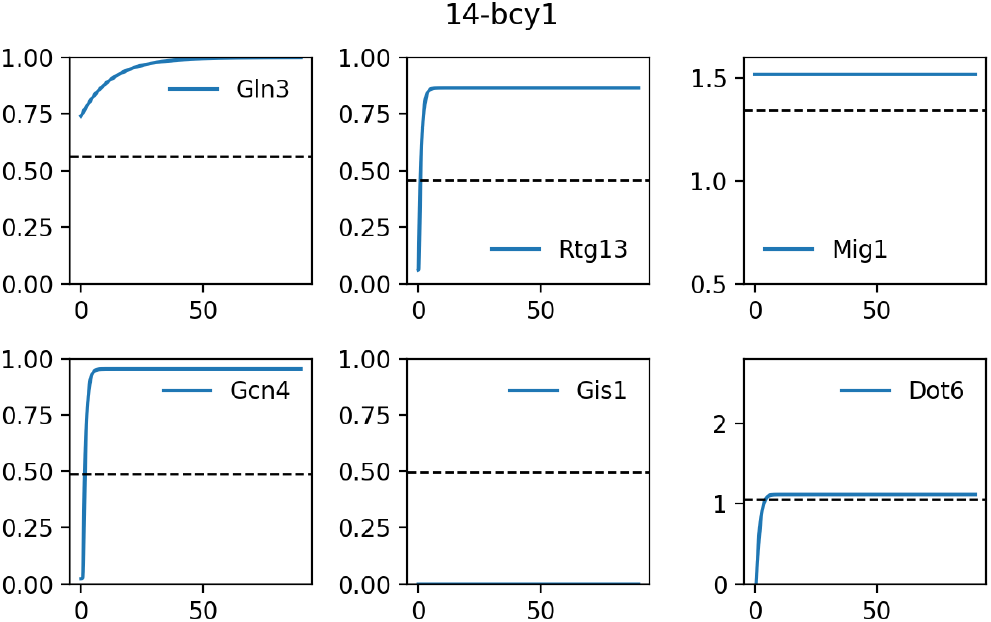

### S4.15 15-ira1

#### Readout used is Gln3 Gcn4

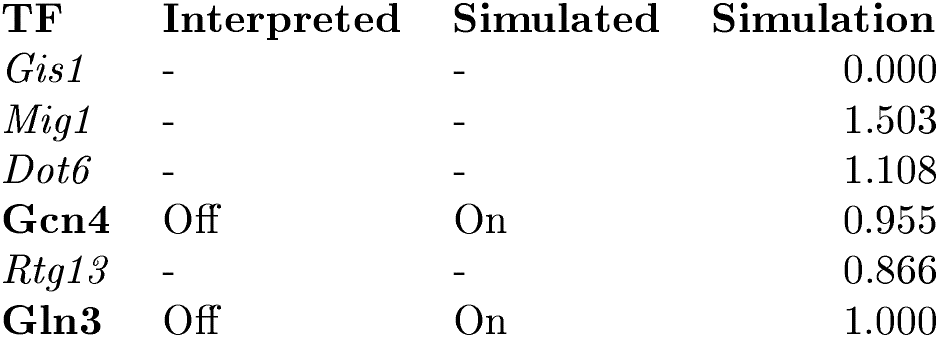

##### Description

Zurita-Martinez et al, 2005 studied a *ira1* strain (S1278b) grown in YP Glucose + 50nM rapamycin.

##### Representation

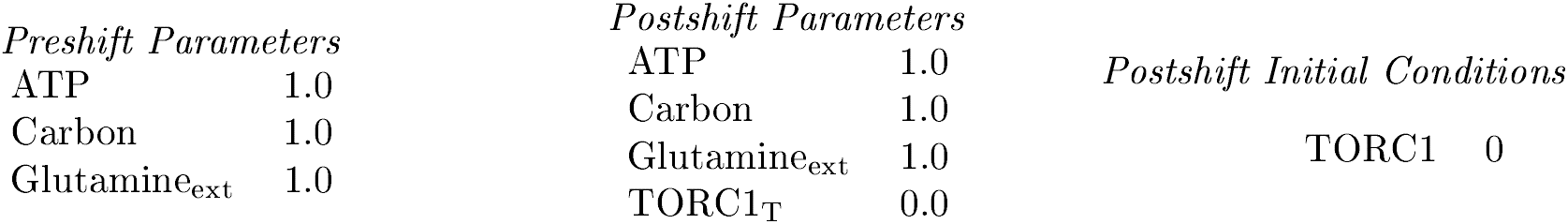

##### Mutant definition

| *Parameters* |  | *Initial conditions* |  |
| --- | --- | --- | --- |
| $w_{raspka}$ | 0.9362 | | |

##### Model does not agree with experiment

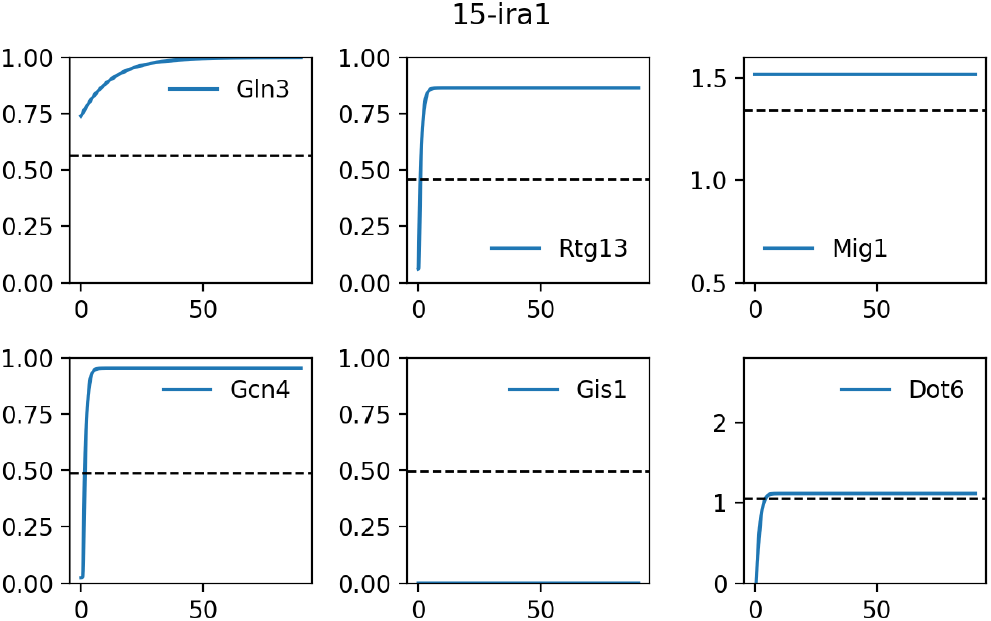

### S4.16 16-ira1 ira2

#### Readout used is Gln3 Gcn4

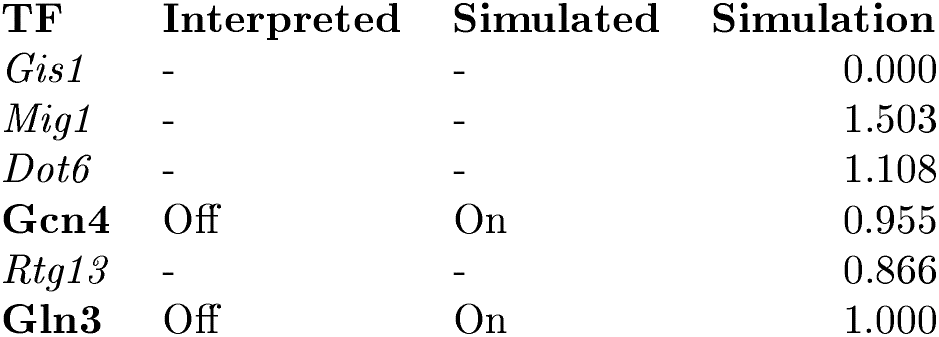

##### Description

Zurita-Martinez et al, 2005 studied a *ira1 ira2* strain (S1278b) grown in YP Glucose + 50nM rapamycin.

##### Representation

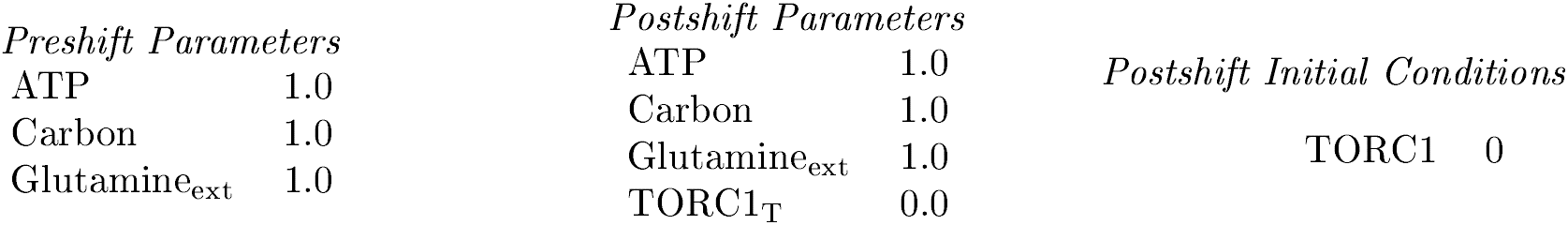

##### Mutant definition

| *Parameters* |  | *Initial conditions* |
| --- | --- | --- |
| w<sub>raspka</sub> | 0.1872 |  |

##### Model does not agree with experiment

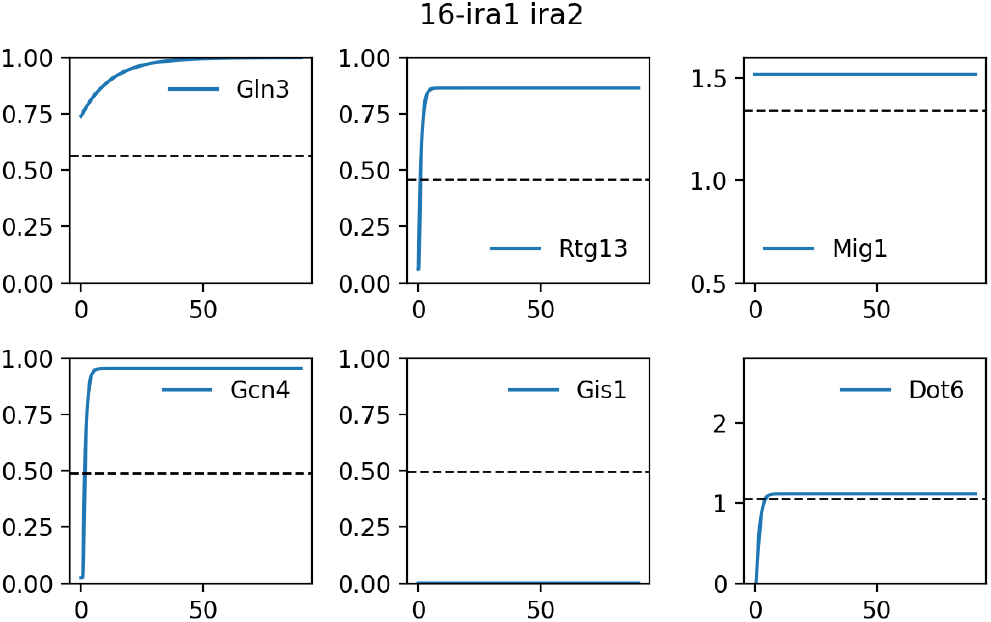

### S4.17 17-ras2

#### Readout used is Gln3 Gcn4

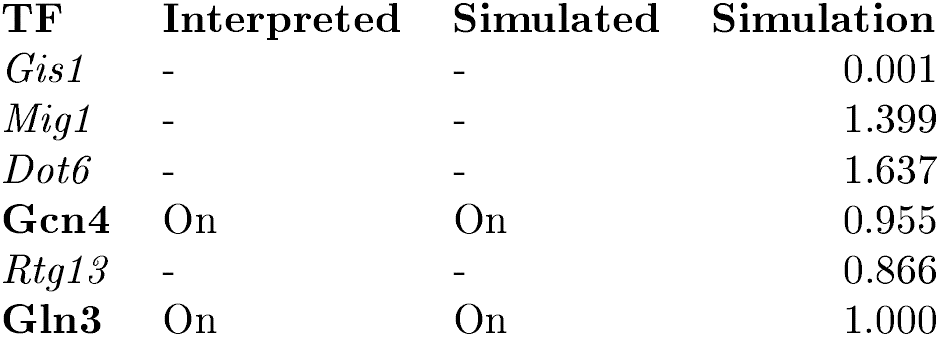

| TF | Interpreted | Simulated | Simulation |
| --- | --- | --- | --- |
| *Gis1* | - | - | 0.001 |
| *Mig1* | - | - | 1.399 |
| *Dot6* | - | - | 1.637 |
| **Gcn4** | On | On | 0.955 |
| *Rtg13* | - | - | 0.866 |
| **Gln3** | On | On | 1.000 |

##### Description

Zurita-Martinez et al, 2005 studied a *ras2* strain (MLY41a) grown in YP Glucose + 50nM rapamycin.

##### Representation

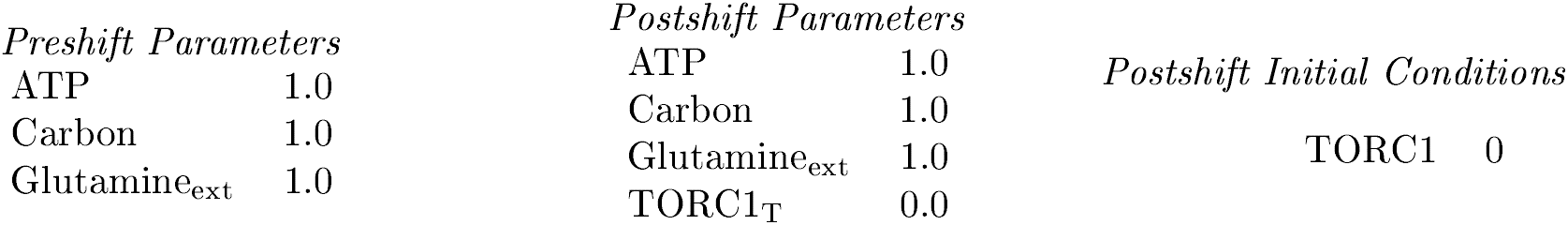

##### Mutant definition

| *Parameters* |  | *Initial conditions* |
| --- | --- | --- |
| Ras<sub>T</sub> | 0.0 |  |

##### Model agrees with experiment

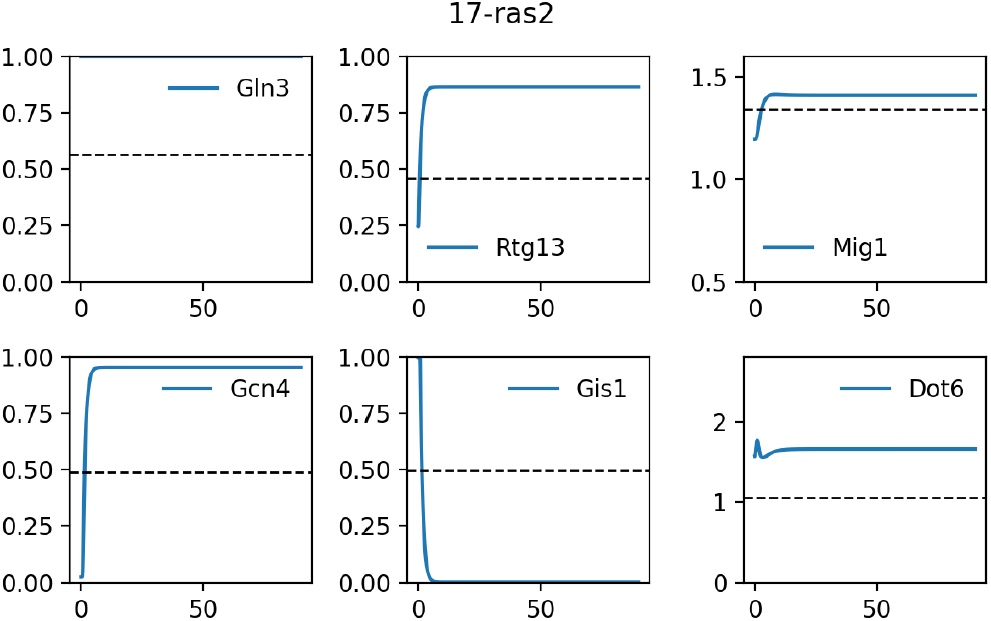

### S4.18 18-tpk1

#### Readout used is Gln3 Gcn4

| TF | Interpreted | Simulated | Simulation |
| --- | --- | --- | --- |
| *Gis1* | - | - | 0.000 |
| *Mig1* | - | - | 1.411 |
| *Dot6* | - | - | 1.578 |
| **Gcn4** | On | On | 0.955 |
| *Rtg13* | - | - | 0.866 |
| **Gln3** | On | On | 1.000 |

##### Description

Zurita-Martinez et al, 2005 studied a *tpk1* strain (MLY41a) grown in YP Glucose + 50nM rapamycin.

##### Representation

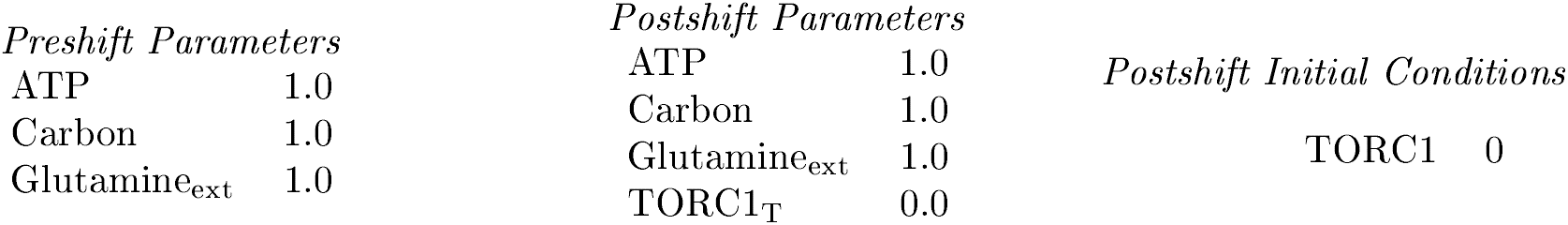

##### Mutant definition

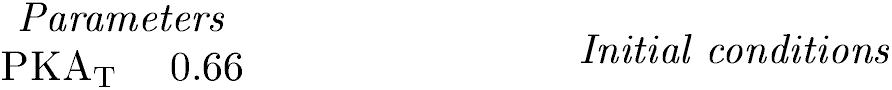

##### Model agrees with experiment

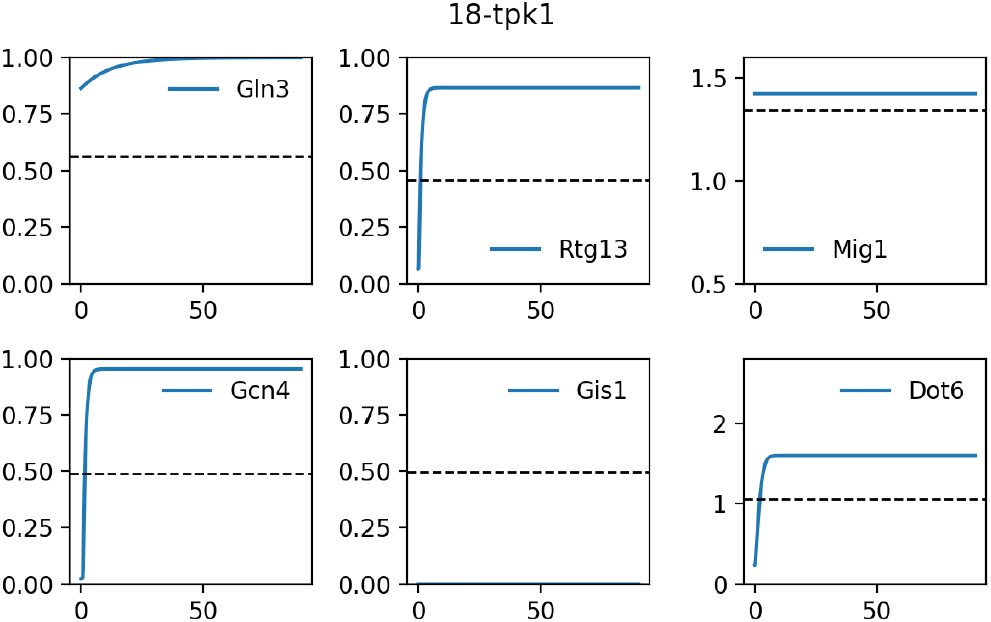

### S4.19 19-RAS2v19 gln3 gat1

#### Readout used is Gln3 Gcn4

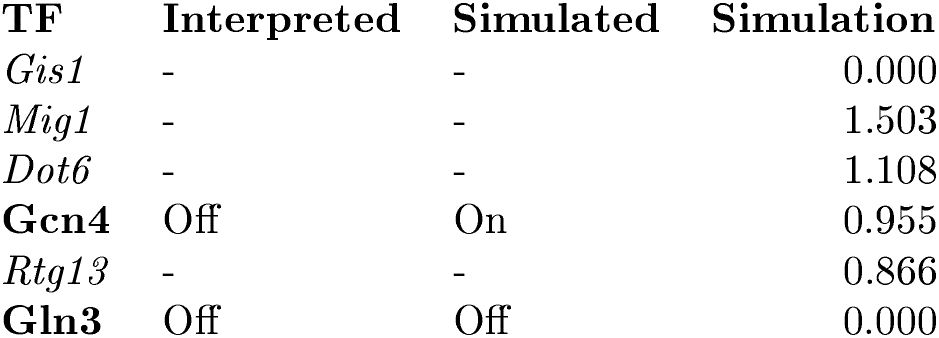

##### Description

Schmelzle et al, 2003 studied a *RAS2v19 gln3 gat1* strain (TB50a) grown in YPD + 200ng/mL rapamycin.

##### Representation

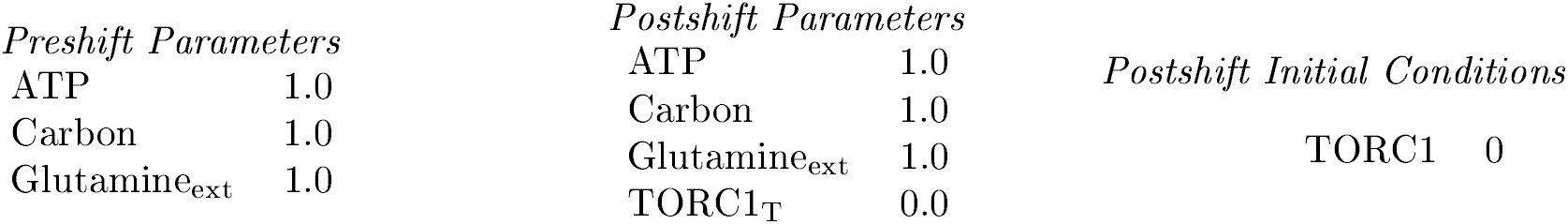

##### Mutant definition

|  |  |
| --- | --- |
| Gln3<sub>T</sub> | 0.0 |
| w<sub>raspka</sub> | 0.0 |

*Initial conditions*

##### Model does not agree with experiment

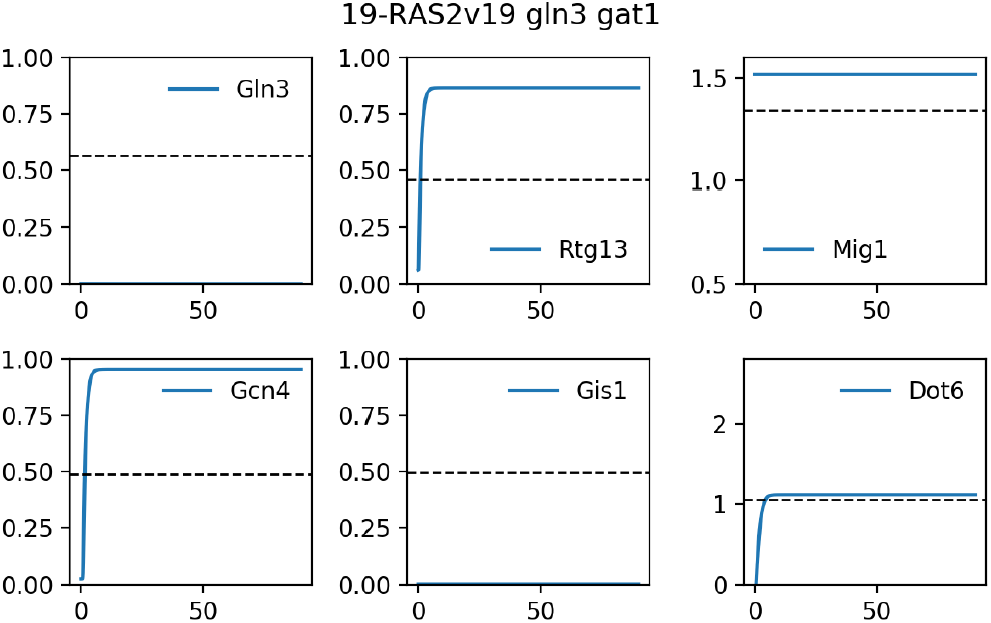

### S4.20 20-TPK1 gln3 gat1

#### Readout used is Gln3 Gcn4

| TF | Interpreted | Simulated | Simulation |
| --- | --- | --- | --- |
| *Gis1* | - | - | 0.177 |
| *Mig1* | - | - | 1.338 |
| *Dot6* | - | - | 1.874 |
| **Gcn4** | Off | On | 0.955 |
| *Rtg13* | - | - | 0.866 |
| **Gln3** | Off | Off | 0.000 |

##### Description

Schmelzle et al, 2003 studied a *TPK1 gln3 gat1* strain (TB50a) grown in YPD + 200ng/mL rapamycin.

##### Representation

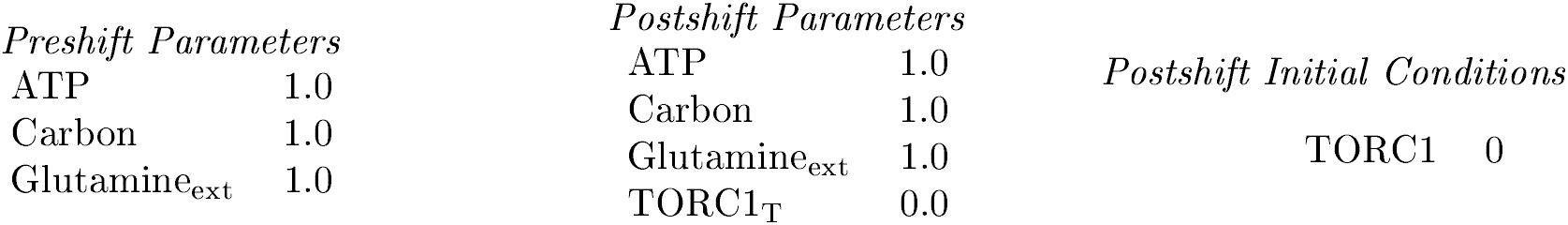

##### Mutant definition

| *Parameters* |  | *Initial conditions* |
| --- | --- | --- |
| Gln3<sub>T</sub> | 0.0 |  |
| w<sub>pkacamp</sub> | 0.0 |  |

##### Model does not agree with experiment

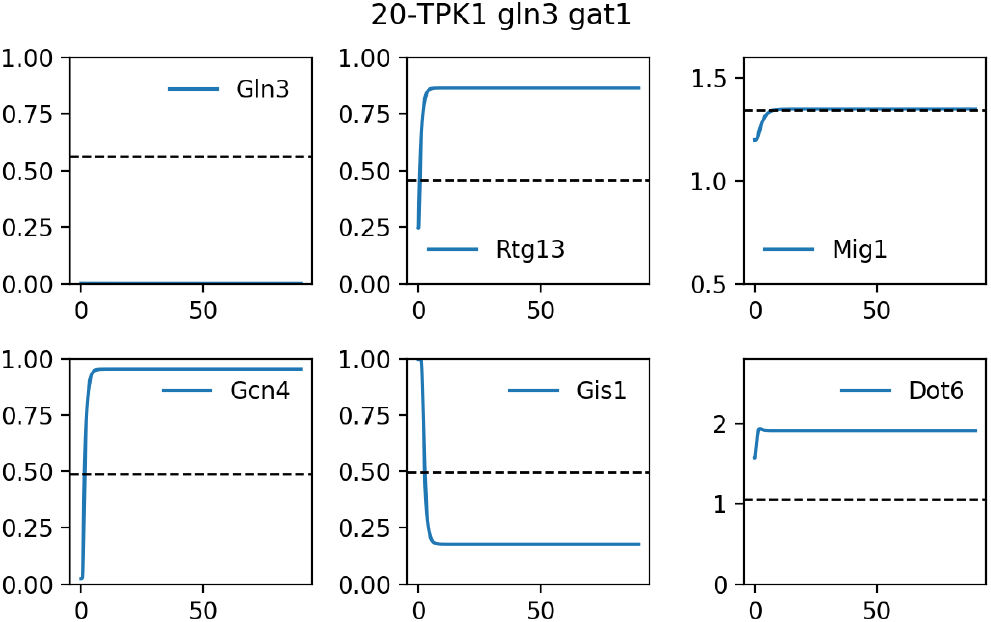

### S4.21 21-bcy1

#### Readout used is Gln3 Gcn4

| TF | Interpreted | Simulated | Simulation |
| --- | --- | --- | --- |
| *Gis1* | - | - | 0.177 |
| *Mig1* | - | - | 1.338 |
| *Dot6* | - | - | 1.874 |
| **Gcn4** | On | On | 0.955 |
| *Rtg13* | - | - | 0.866 |
| **Gln3** | On | On | 1.000 |

##### Description

Schmelzle et al, 2003 studied a *bcy1* strain (TB50a) grown in YPD + 200ng/mL rapamycin.

##### Representation

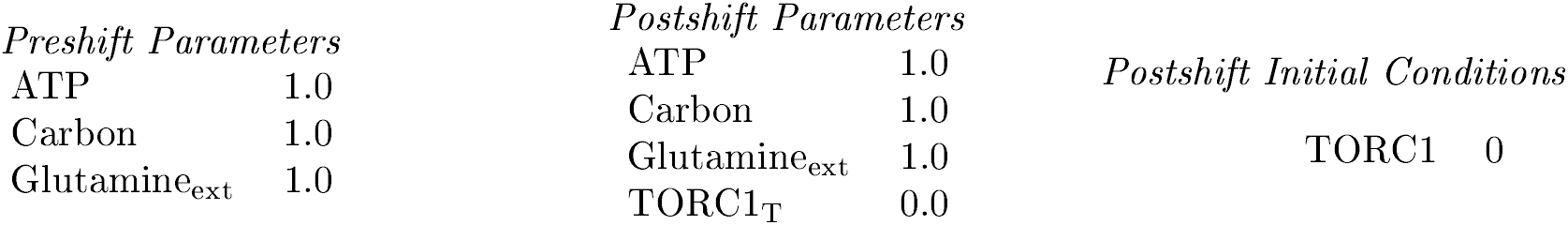

##### Mutant definition

| *Parameters* |  | *Initial conditions* |
| --- | --- | --- |
| w<sub>pkacamp</sub> | 0.0 |  |

##### Model agrees with experiment

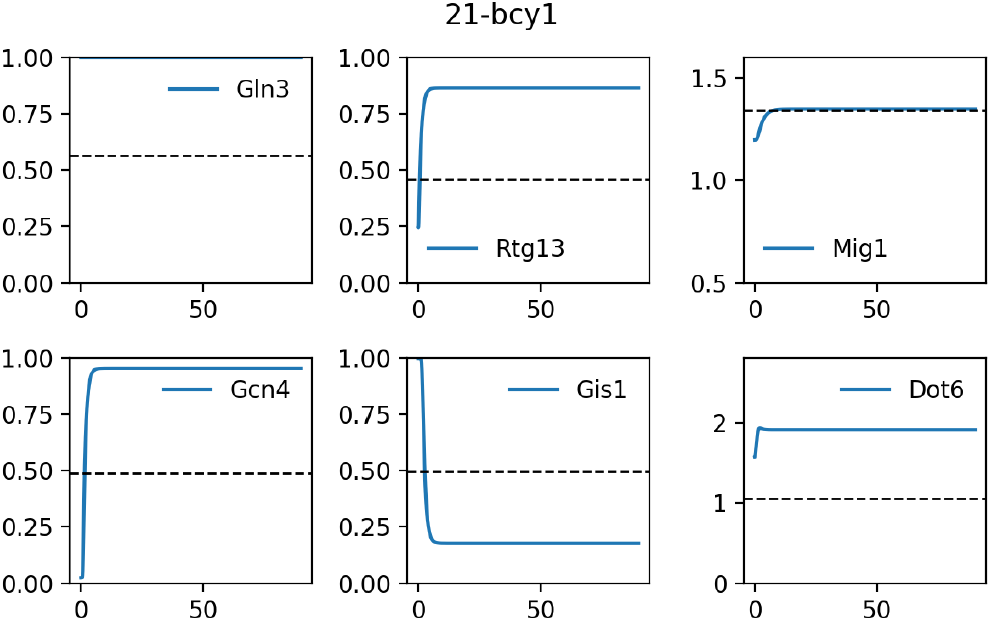

### S4.22 22-bcy1 gln3 gat1

#### Readout used is Gln3 Gcn4

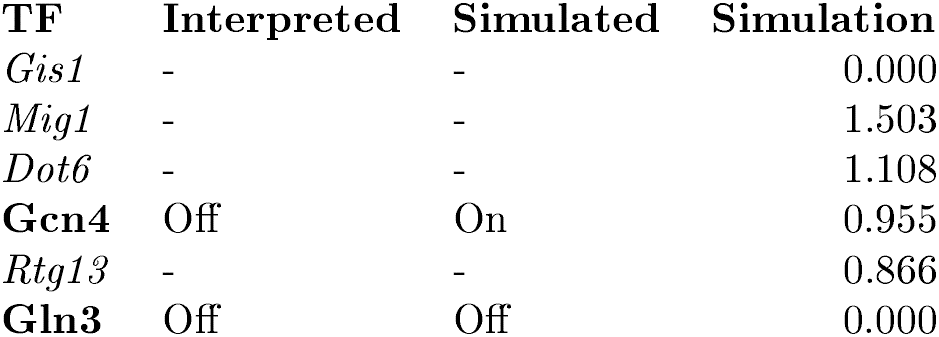

##### Description

Schmelzle et al, 2003 studied a *bcy1 gln3 gat1* strain (TB50alpha) grown in YPD + 200ng/mL rapamycin.

##### Representation

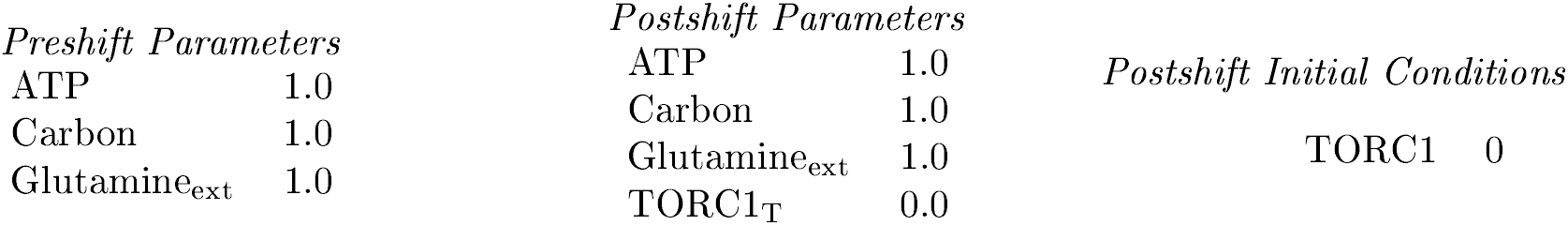

##### Mutant definition

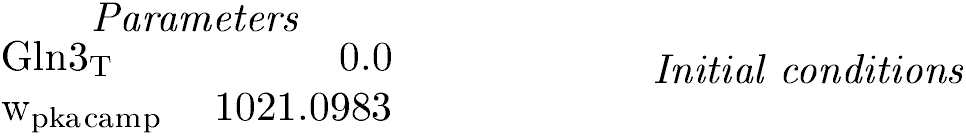

##### Model does not agree with experiment

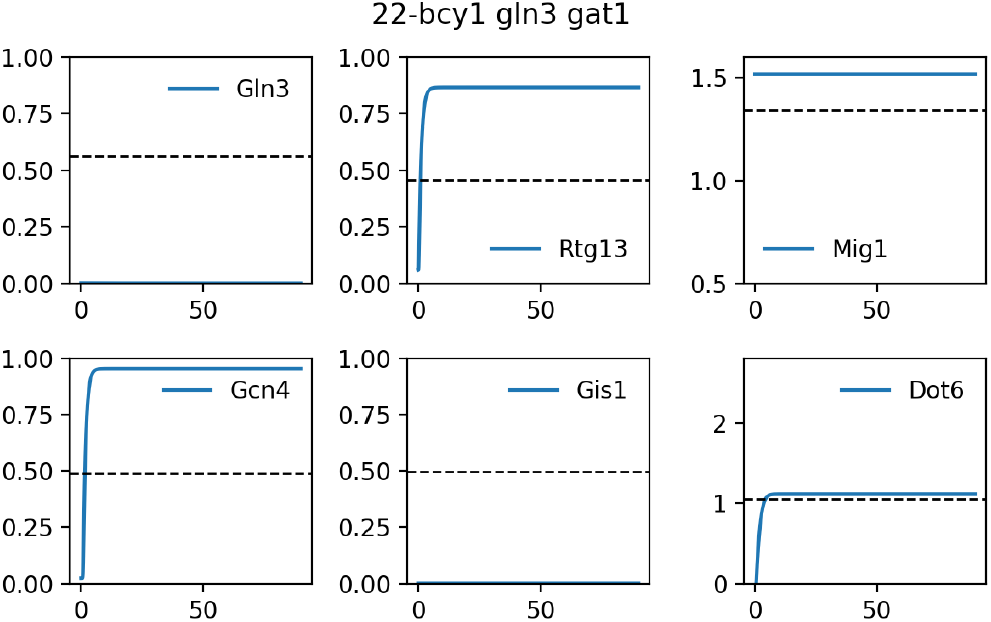

### S4.23 23-wt

| TF | Interpreted | Simulated | Simulation |
| --- | --- | --- | --- |
| *Gis1* | - | - | 0.000 |
| *Mig1* | - | - | 1.503 |
| *Dot6* | - | - | 1.037 |
| **Gcn4** | On | On | 0.806 |
| *Rtg13* | - | - | 0.702 |
| *Gln3* | - | - | 0.999 |

#### Description

Cherkasova et al, 2010 studied a *wt* strain (H1642) grown in SC + 10mM 3AT.

#### Representation

**Representation:**
| *Preshift Parameters* |  | *Postshift Parameters* |  | *Postshift Initial Conditions* |
| --- | --- | --- | --- | --- |
| ATP | 1.0 | ATP | 1.0 |  |
| Carbon | 1.0 | Carbon | 1.0 |  |
| Glutamine<sub>ext</sub> | 0.0 | Glutamine<sub>ext</sub> | 0.0 |  |
| NH4 | 2.0 | NH4 | 2.0 |  |
|  |  | k<sub>accnh4</sub> | 0.0001 |  |

#### Mutant definition

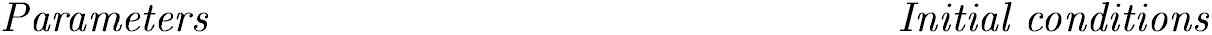

#### Model agrees with experiment

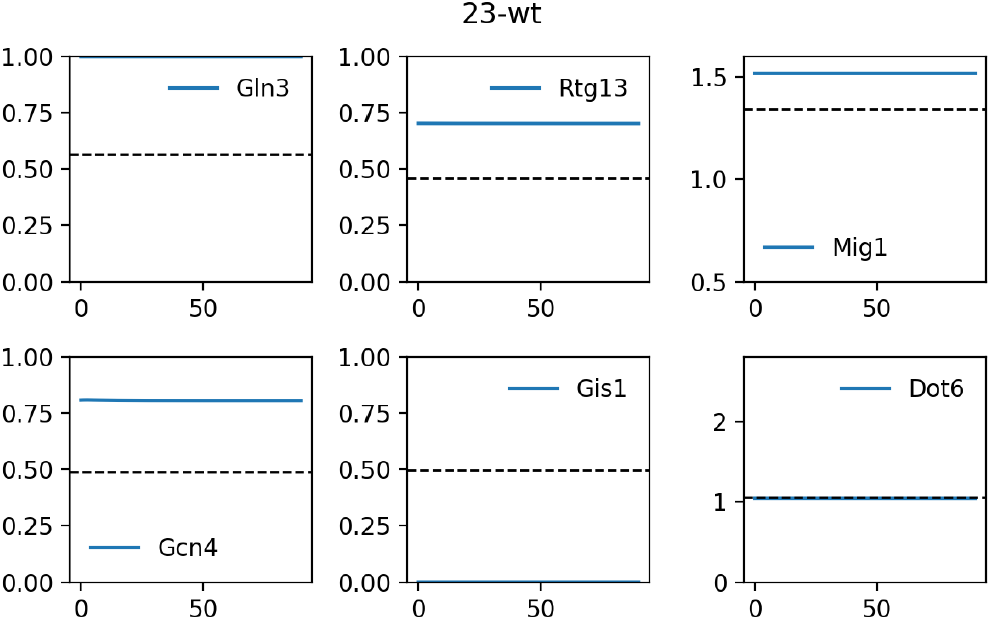

### S4.24 24-gcn2

| TF | Interpreted | Simulated | Simulation |
| --- | --- | --- | --- |
| *Gis1* | - | - | 0.000 |
| *Mig1* | - | - | 1.503 |
| *Dot6* | - | - | 1.037 |
| **Gcn4** | Off | Off | 0.024 |
| *Rtg13* | - | - | 0.702 |
| *Gln3* | - | - | 0.999 |

#### Description

Cherkasova et al, 2010 studied a *gcn2* strain (H1895) grown in SC + 10mM 3AT.

#### Representation

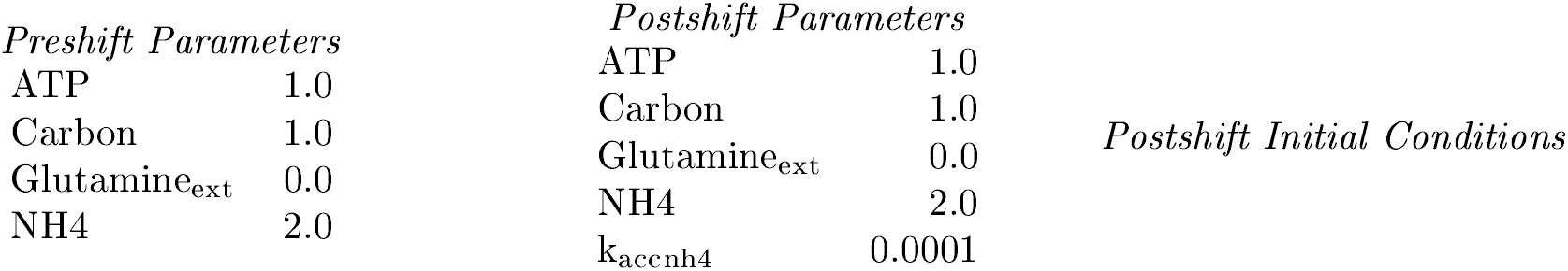

#### Mutant definition

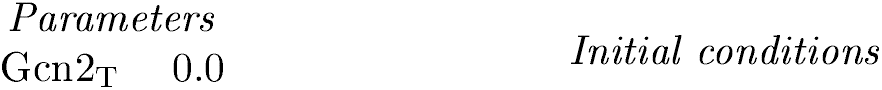

#### Model agrees with experiment

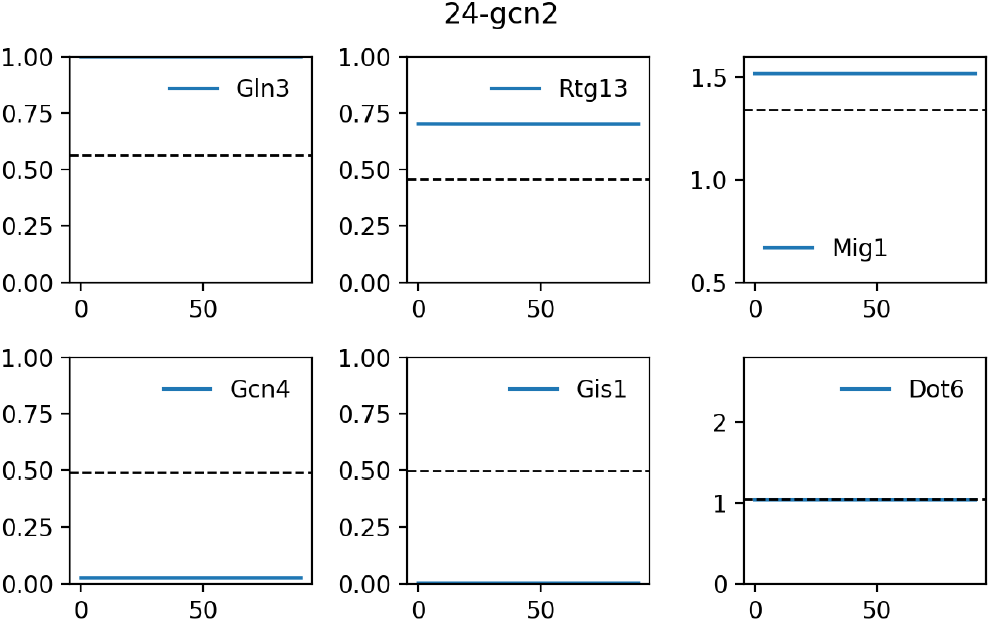

### S4.25 25-snf1

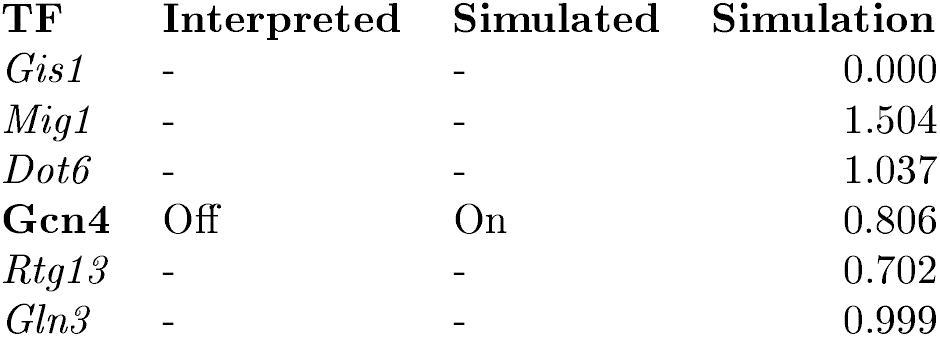

| TF | Interpreted | Simulated | Simulation |
| --- | --- | --- | --- |
| *Gis1* | - | - | 0.000 |
| *Mig1* | - | - | 1.504 |
| *Dot6* | - | - | 1.037 |
| **Gcn4** | Off | On | 0.806 |
| *Rtg13* | - | - | 0.702 |
| *Gln3* | - | - | 0.999 |

#### Description

Cherkasova et al, 2010 studied a *snf1* strain (HQY343) grown in SC + 10mM 3AT.

#### Representation

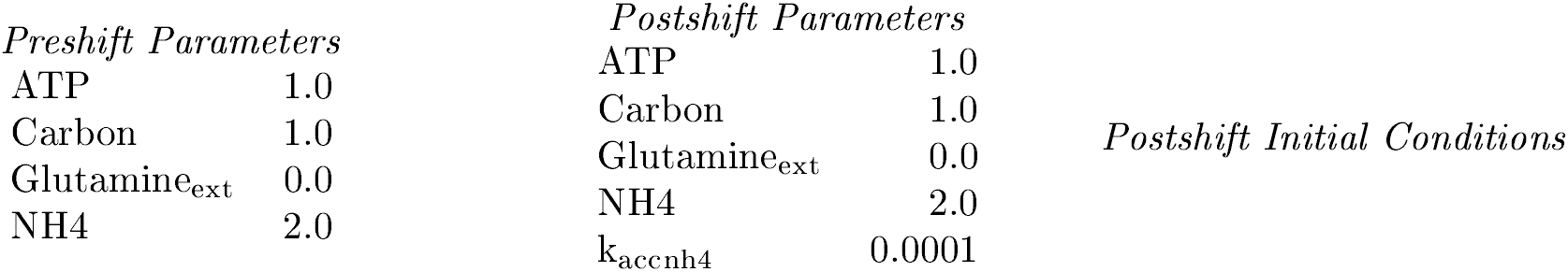

#### Mutant definition

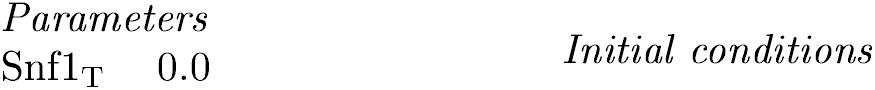

#### Model does not agree with experiment

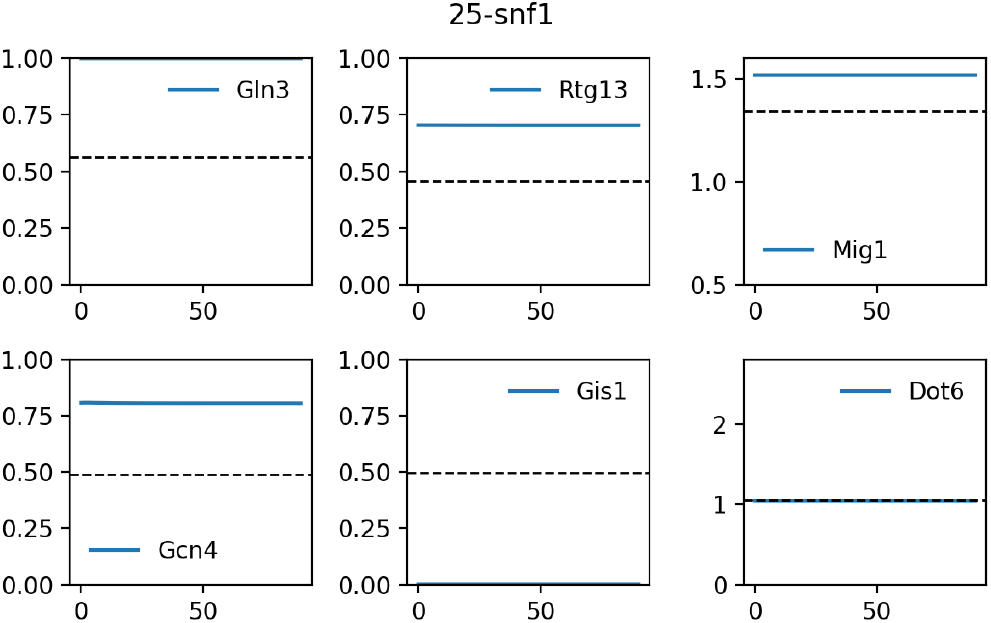

### S4.26 26-gcn2 snf1

| TF | Interpreted | Simulated | Simulation |
| --- | --- | --- | --- |
| *Gis1* | - | - | 0.000 |
| *Mig1* | - | - | 1.504 |
| *Dot6* | - | - | 1.037 |
| **Gcn4** | Off | Off | 0.024 |
| *Rtg13* | - | - | 0.702 |
| *Gln3* | - | - | 0.999 |

#### Description

Cherkasova et al, 2010 studied a *gcn2 snf1* strain (HQY344) grown in SC + 10mM 3AT.

#### Representation

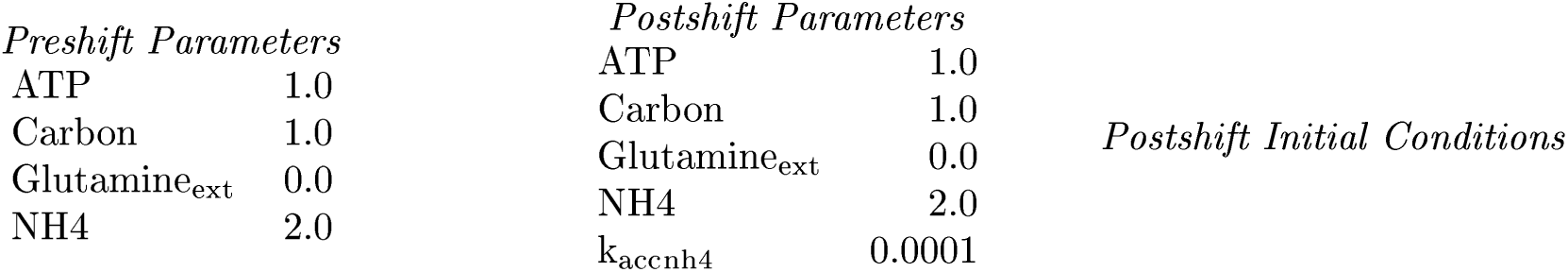

#### Mutant definition

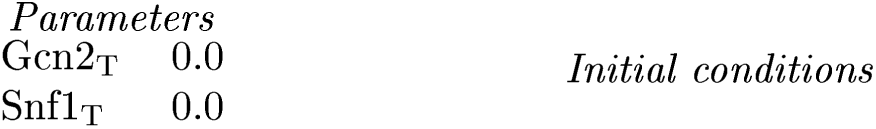

| *Parameters* |  | *Initial conditions* |
| --- | --- | --- |
| Gcn2<sub>T</sub> | 0.0 |  |
| Snf1<sub>T</sub> | 0.0 |  |

#### Model agrees with experiment

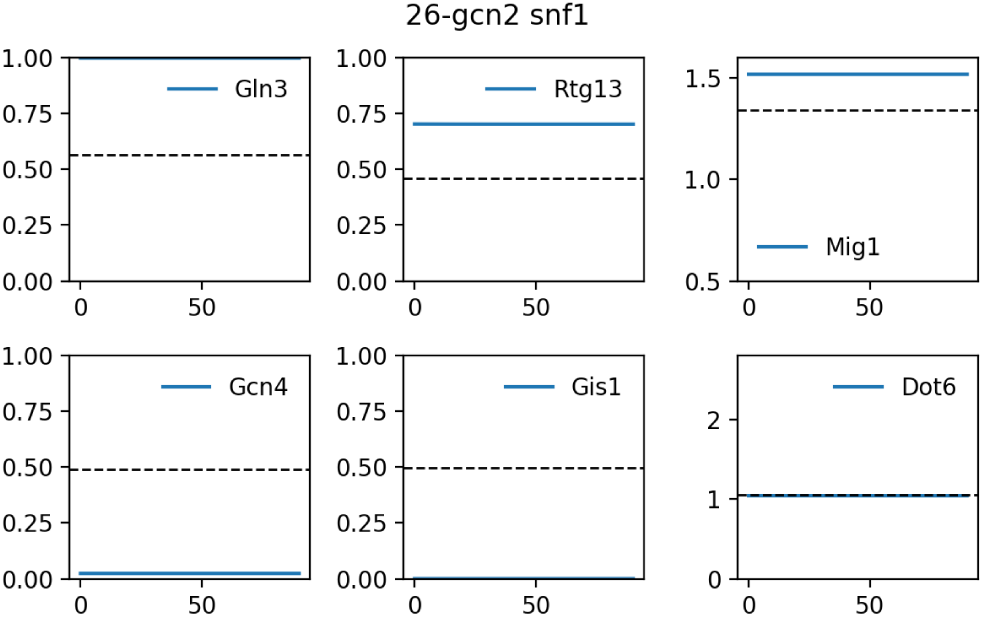

### S4.27 27-gln3 gcn4

#### Readout used is Gln3 Gcn4

| TF | Interpreted | Simulated | Simulation |
| --- | --- | --- | --- |
| *Gis1* | - | - | 0.000 |
| *Mig1* | - | - | 1.503 |
| *Dot6* | - | - | 1.108 |
| **Gcn4** | Off | Off | 0.000 |
| *Rtg13* | - | - | 0.866 |
| **Gln3** | Off | Off | 0.000 |

##### Description

Valenzuela et al, 2001 studied a *gln3 gcn4* strain (CLA-303) grown in YPD 200ng/mL rapamycin.

##### Representation

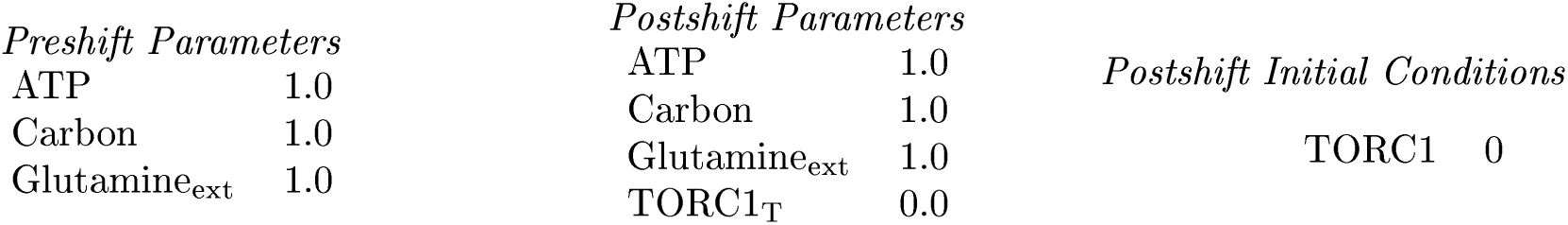

##### Mutant definition

|  |  |  |
| --- | --- | --- |
| *Parameters* |  | *Initial conditions* |
| Gcn4<sub>T</sub> | 0.0 |  |
| Gln3<sub>T</sub> | 0.0 |  |

##### Model agrees with experiment

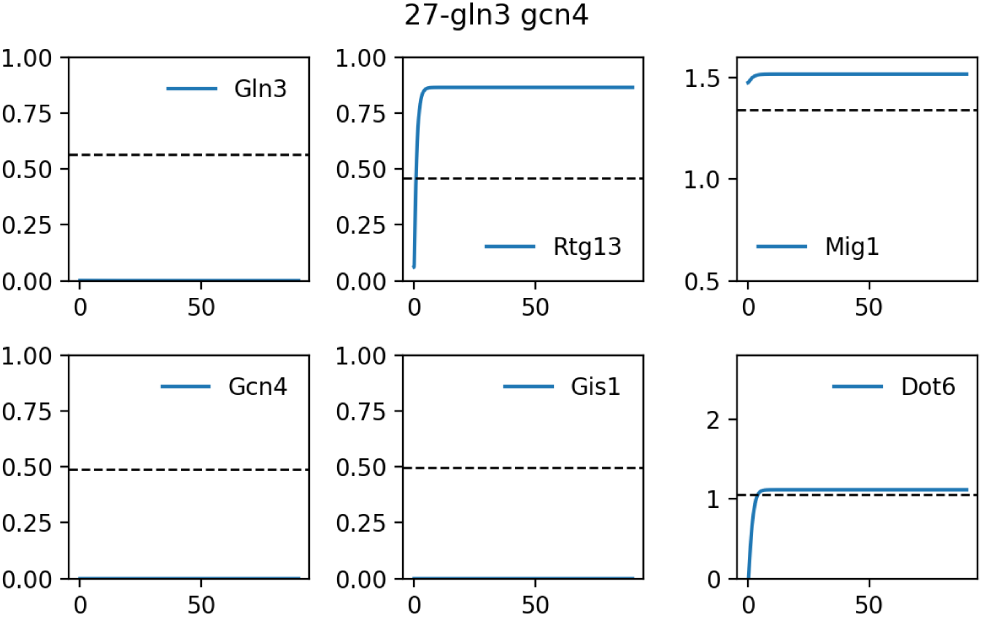

### S4.28 28-gcn4

#### Readout used is Gln3 Gcn4

| TF | Interpreted | Simulated | Simulation |
| --- | --- | --- | --- |
| *Gis1* | - | - | 0.000 |
| *Mig1* | - | - | 1.503 |
| *Dot6* | - | - | 1.108 |
| **Gcn4** | On | Off | 0.000 |
| *Rtg13* | - | - | 0.866 |
| **Gln3** | On | On | 1.000 |

##### Description

Valenzuela et al, 2001 studied a *gcn4* strain (CLA-300) grown in YPD 200ng/mL rapamycin.

##### Representation

**Representation:**
|  |  |  |  |  |
| --- | --- | --- | --- | --- |
| *Preshift Parameters* |  | *Postshift Parameters* |  | *Postshift Initial Conditions* |
| ATP | 1.0 | ATP | 1.0 |  |
| Carbon | 1.0 | Carbon | 1.0 |  |
| Glutamine<sub>ext</sub> | 0.0 | Glutamine<sub>ext</sub> | 1.0 |  |
|  |  | TORC1<sub>T</sub> | 0.0 |  |
|  |  |  |  | TORC1 0 |

##### Mutant definition

|  |  |  |
| --- | --- | --- |
| *Parameters* |  | *Initial conditions* |
| Gcn4<sub>T</sub> | 0.0 |  |

##### Model does not agree with experiment

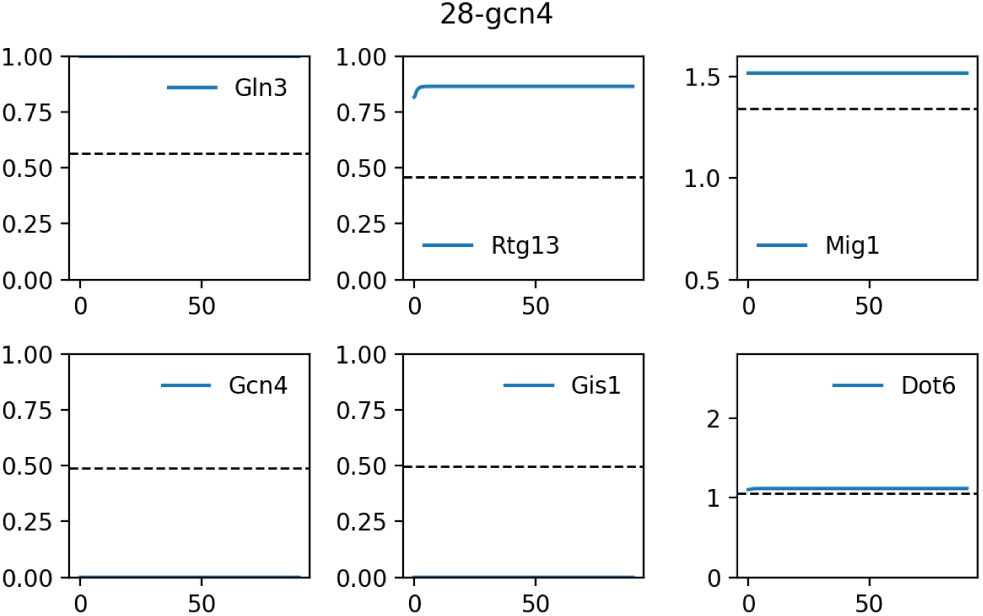

### S4.29 29-rph1 gis1

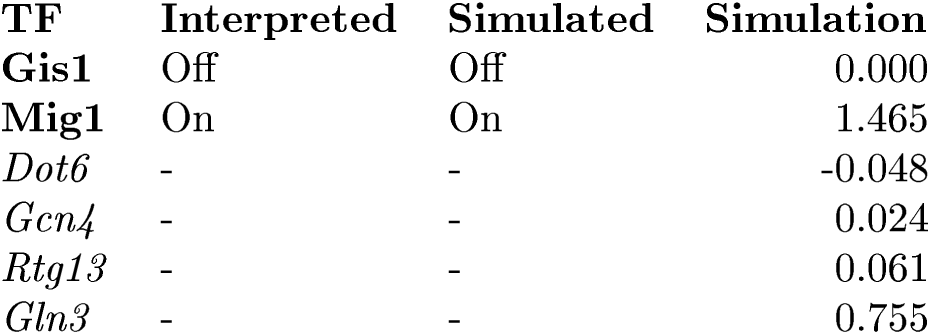

#### Description

H. Ronne et al, 1999 studied a *rph1 gis1* strain (H874) grown in SD glucose-ura.

#### Representation

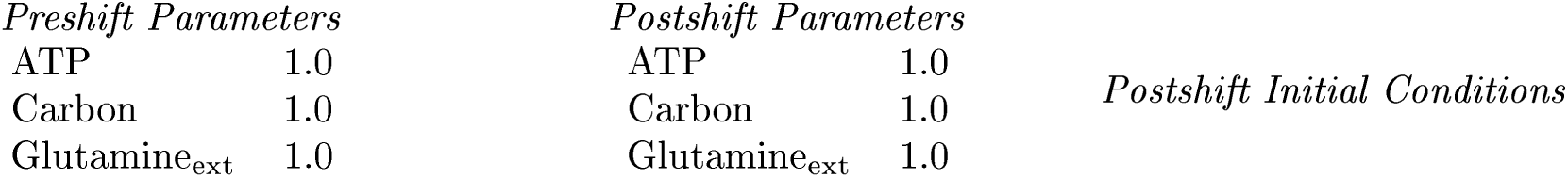

#### Mutant definition

|  |  |
| --- | --- |
| Gis1<sub>T</sub> | 0.0 |

*Initial conditions*

#### Model agrees with experiment

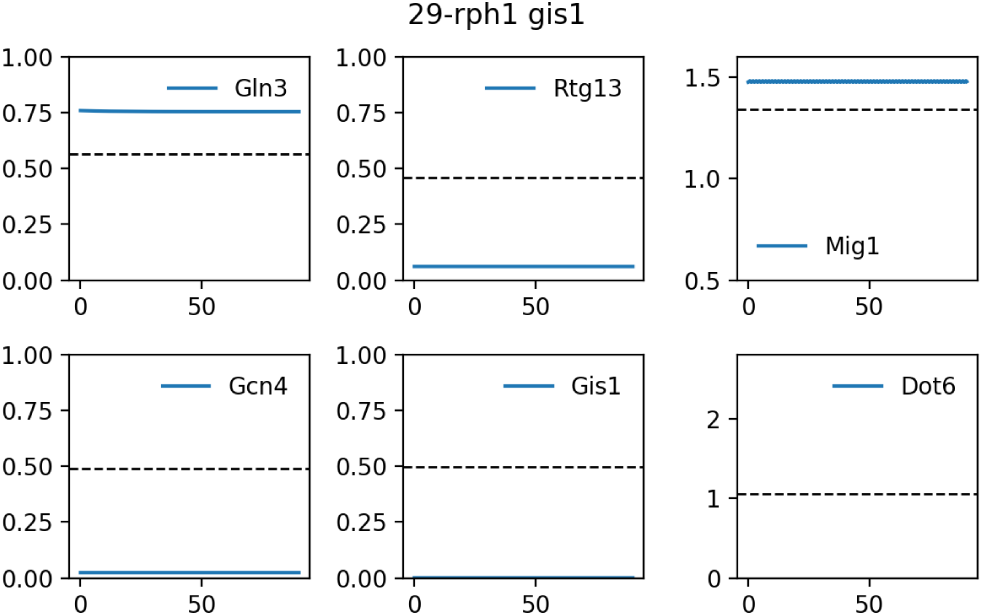

### S4.30 30-rph1 gis1

| TF | Interpreted | Simulated | Simulation |
| --- | --- | --- | --- |
| **Gis1** | Off | Off | 0.000 |
| **Mig1** | Off | Off | 1.168 |
| *Dot6* | - | - | 2.078 |
| *Gcn4* | - | - | 0.608 |
| *Rtg13* | - | - | 0.664 |
| *Gln3* | - | - | 1.000 |

#### Description

H. Ronne et al, 1999 studied a *rph1 gis1* strain (H874) grown in YP ethanol.

#### Representation

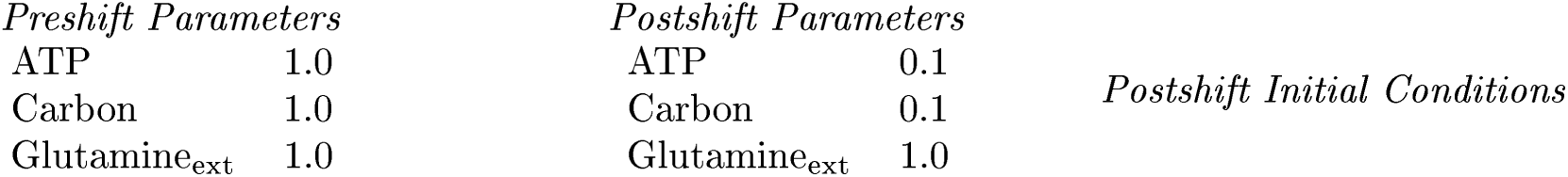

#### Mutant definition

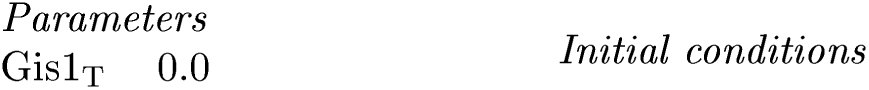

#### Model agrees with experiment

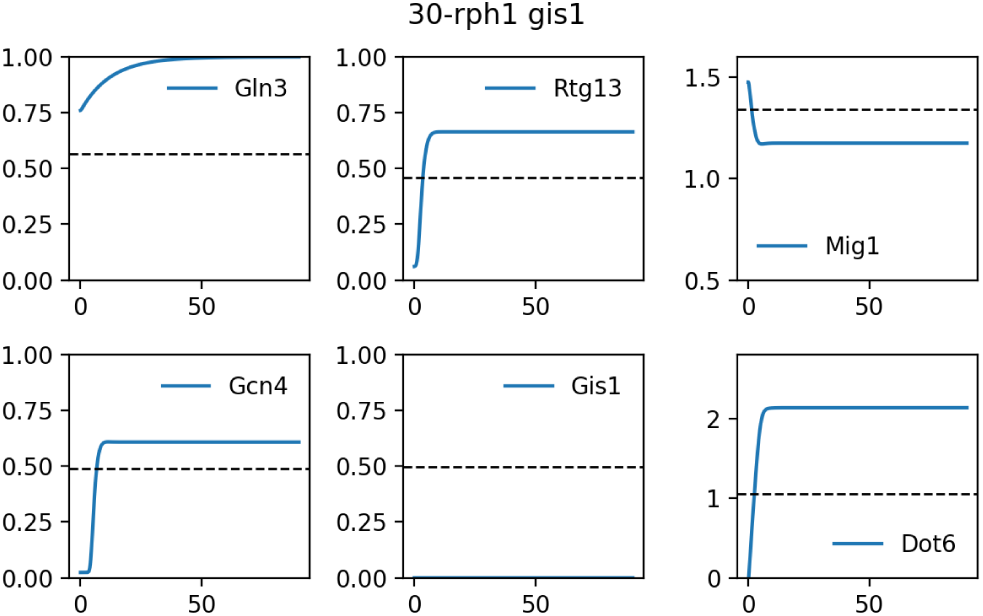

### S4.31 31-mig1 snf1 pde2

| TF | Interpreted | Simulated | Simulation |
| --- | --- | --- | --- |
| **Gis1** | Off | Off | 0.000 |
| *Mig1* | - | - | -45.173 |
| *Dot6* | - | - | -0.256 |
| *Gcn4* | - | - | 0.024 |
| *Rtg13* | - | - | 0.058 |
| *Gln3* | - | - | 0.661 |

#### Description

H. Ronne et al, 1999 studied a *mig1 snf1 pde2* strain (H808) grown in SD glucose.

#### Representation

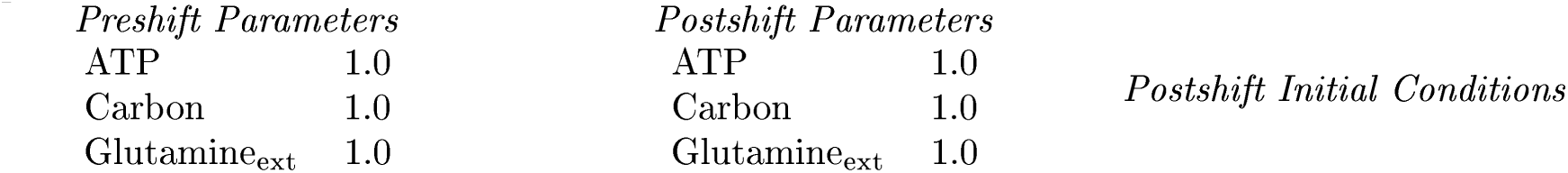

#### Mutant definition

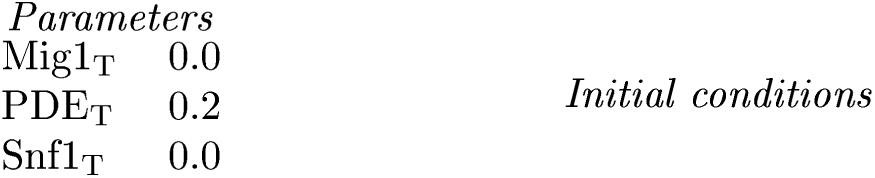

#### Model agrees with experiment

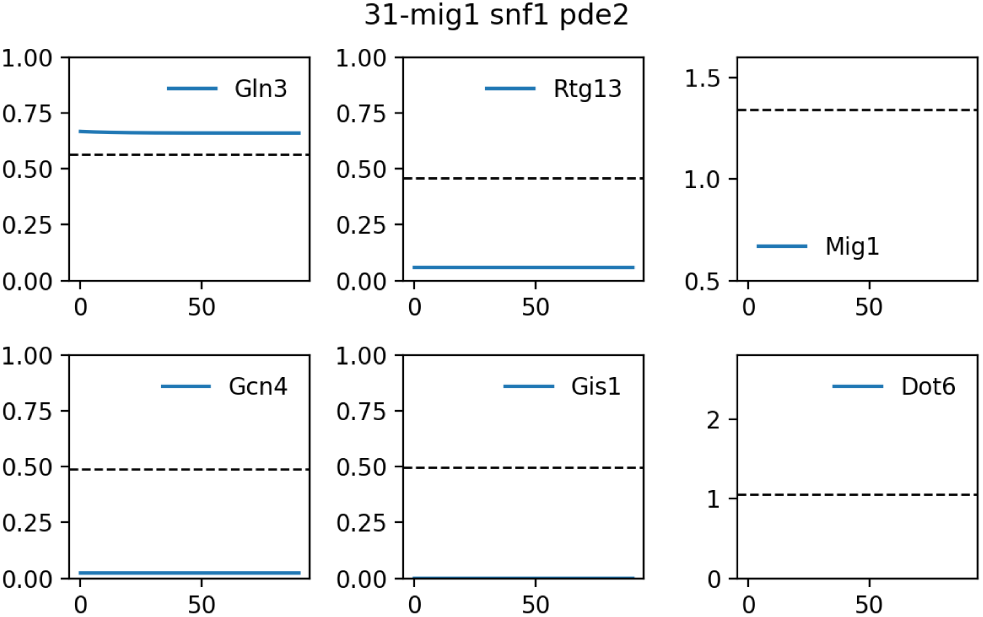

### S4.32 32-mig1 snf1 pde2

| TF | Interpreted | Simulated | Simulation |
| --- | --- | --- | --- |
| **Gis1** | Off | Off | 0.000 |
| *Mig1* | - | - | -45.182 |
| *Dot6* | - | - | -0.145 |
| *Gcn4* | - | - | 0.024 |
| *Rtg13* | - | - | 0.058 |
| *Gln3* | - | - | 0.661 |

#### Description

H. Ronne et al, 1999 studied a *mig1 snf1 pde2* strain (H808) grown in SD raffinose.

#### Representation

**Representation:**
| *Preshift Parameters* |  | *Postshift Parameters* |  | *Postshift Initial Conditions* |
| --- | --- | --- | --- | --- |
| ATP | 1.0 | ATP | 0.7 |  |
| Carbon | 1.0 | Carbon | 0.7 |  |
| Glutamine<sub>ext</sub> | 1.0 | Glutamine<sub>ext</sub> | 1.0 |  |

#### Mutant definition

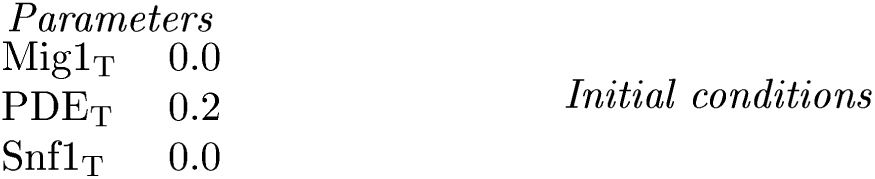

#### Model agrees with experiment

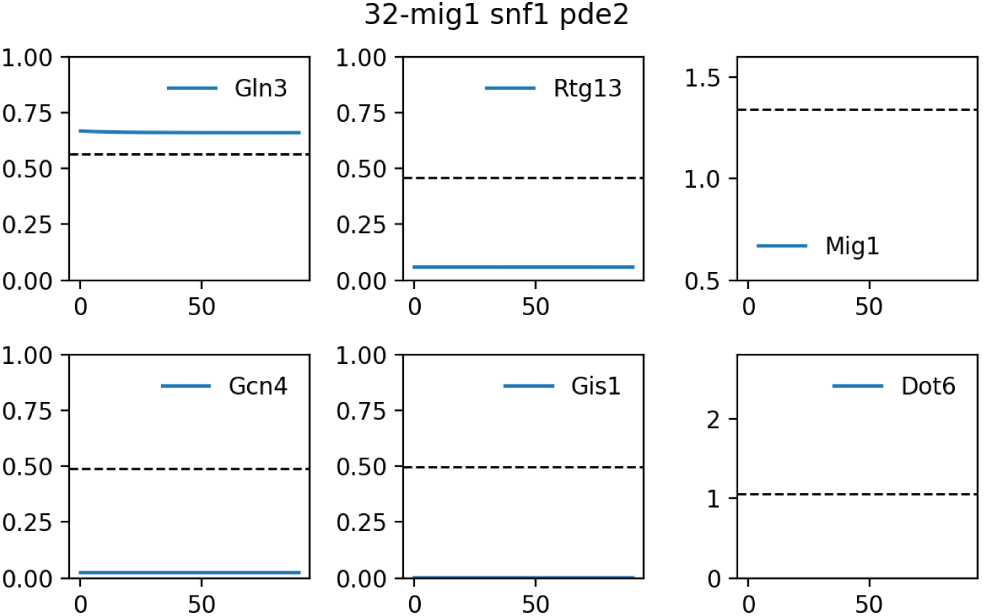

### S4.33 33-mig1 snf1 pde2

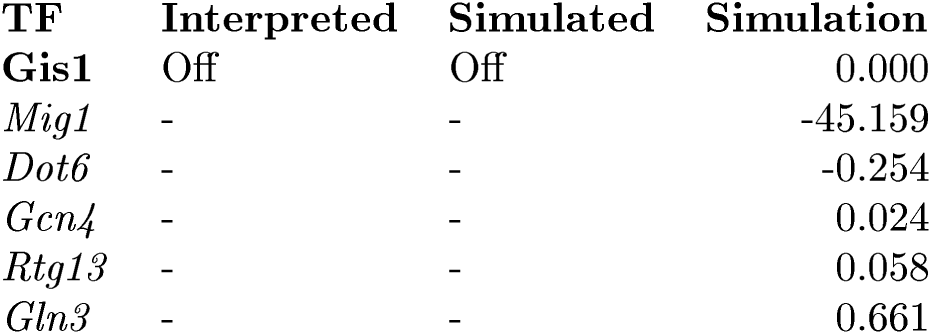

| TF | Interpreted | Simulated | Simulation |
| --- | --- | --- | --- |
| **Gis1** | Off | Off | 0.000 |
| *Mig1* | - | - | -45.159 |
| *Dot6* | - | - | -0.254 |
| *Gcn4* | - | - | 0.024 |
| *Rtg13* | - | - | 0.058 |
| *Gln3* | - | - | 0.661 |

#### Description

H. Ronne et al, 1999 studied a *mig1 snf1 pde2* strain (H808) grown in SD galactose.

#### Representation

**Representation:**
| *Preshift Parameters* |  | *Postshift Parameters* |  | *Postshift Initial Conditions* |
| --- | --- | --- | --- | --- |
| ATP | 1.0 | ATP | 0.8 |  |
| Carbon | 1.0 | Carbon | 0.8 |  |
| Glutamine<sub>ext</sub> | 1.0 | Glutamine<sub>ext</sub> | 1.0 |  |

#### Mutant definition

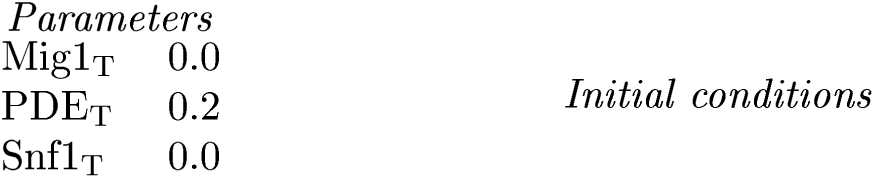

#### Model agrees with experiment

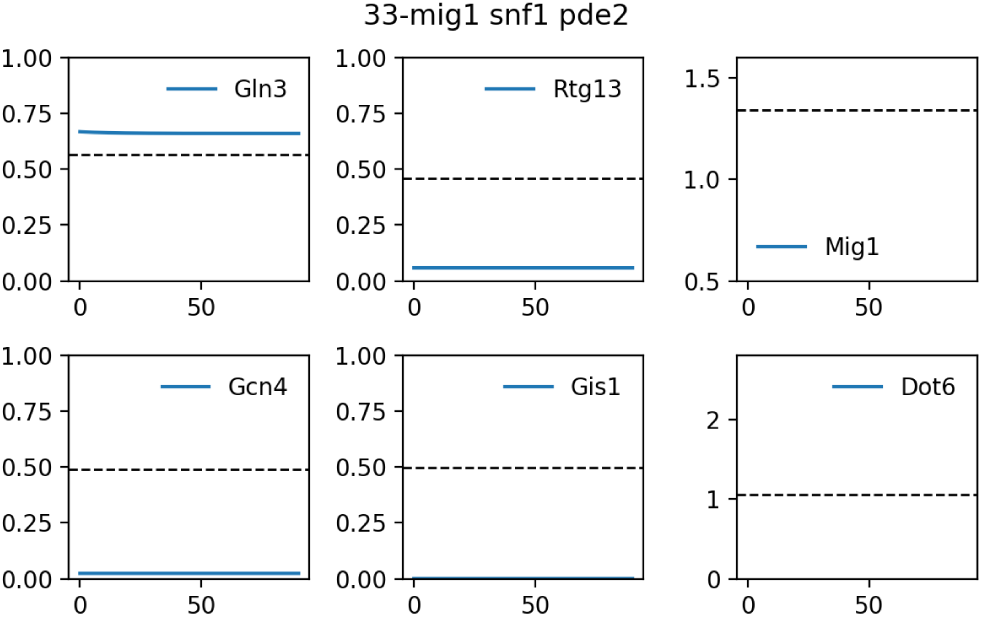

### S4.34 34-snf1

#### Readout used is Gln3 Gcn4

| TF | Interpreted | Simulated | Simulation |
| --- | --- | --- | --- |
| *Gis1* | - | - | 0.000 |
| *Mig1* | - | - | 1.504 |
| *Dot6* | - | - | 1.108 |
| **Gcn4** | Off | On | 0.955 |
| *Rtg13* | - | - | 0.866 |
| **Gln3** | Off | On | 1.000 |

##### Description

Bertram et al, 2002 studied a *snf1* strain (SZy686) grown in YPD + rapamycin.

##### Representation

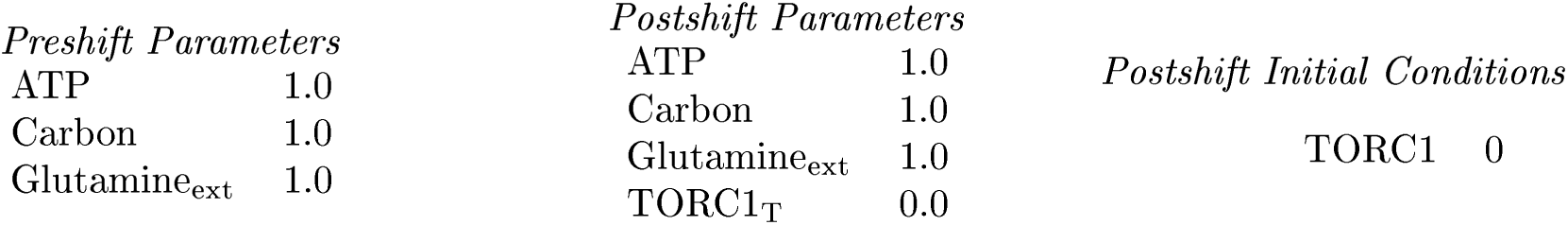

##### Mutant definition

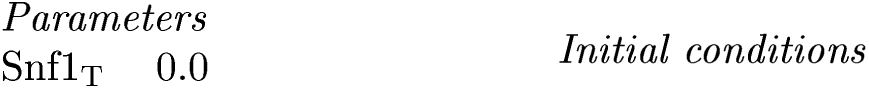

##### Model does not agree with experiment

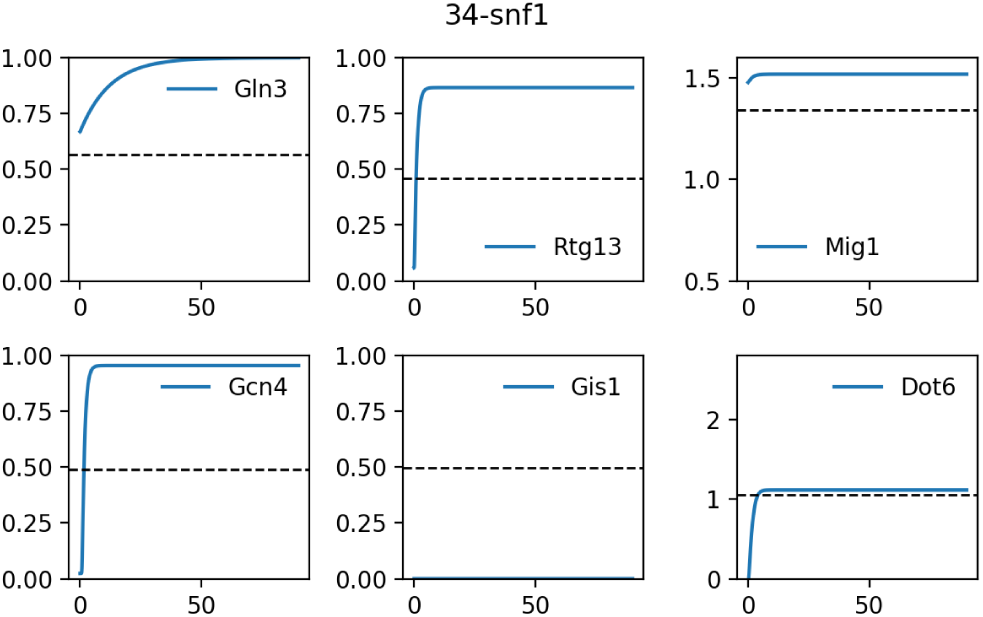

### S4.35 35-reg1

#### Readout used is Gln3 Gcn4

| TF | Interpreted | Simulated | Simulation |
| --- | --- | --- | --- |
| *Gis1* | - | - | 0.000 |
| *Mig1* | - | - | 1.478 |
| *Dot6* | - | - | 1.108 |
| **Gcn4** | On | On | 0.955 |
| *Rtg13* | - | - | 0.866 |
| **Gln3** | On | On | 1.000 |

##### Description

Bertram et al, 2002 studied a *reg1* strain (JC426) grown in YPD + rapamycin.

##### Representation

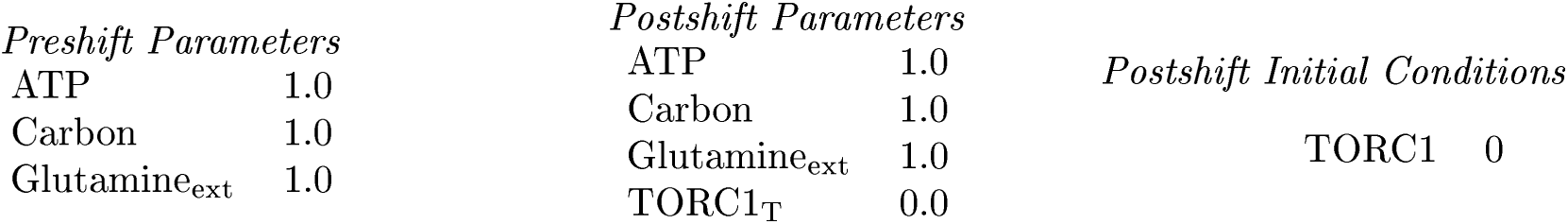

##### Mutant definition

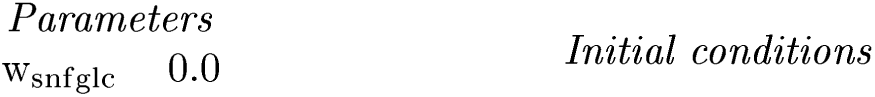

##### Model agrees with experiment

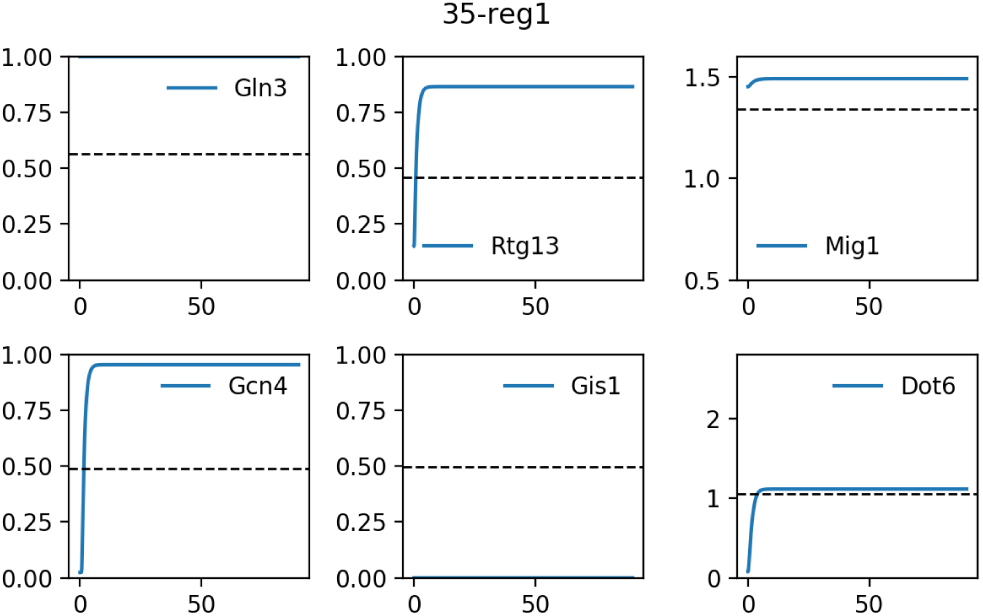

### S4.36 36-ure2

#### Readout used is Gln3 Gcn4

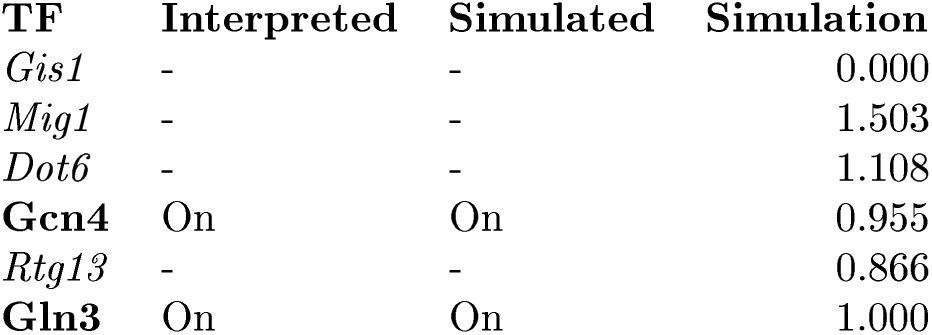

#### Description

Bertram et al, 2002 studied a *ure2* strain (Szy145) grown in YPD + rapamycin.

##### Representation

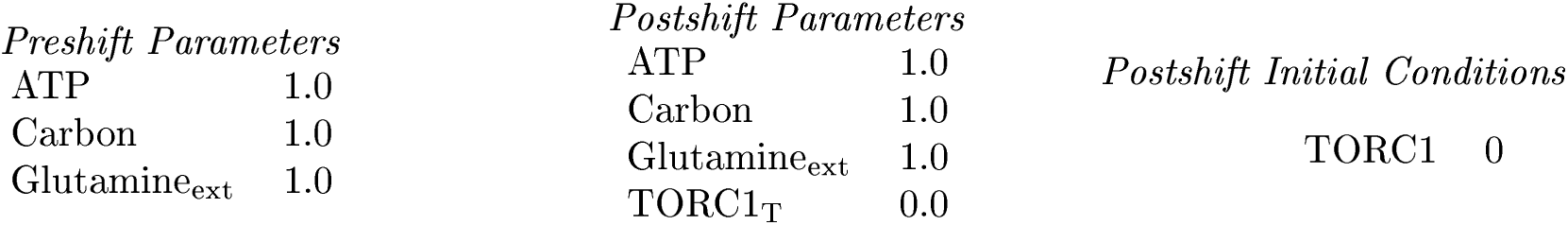

##### Mutant definition

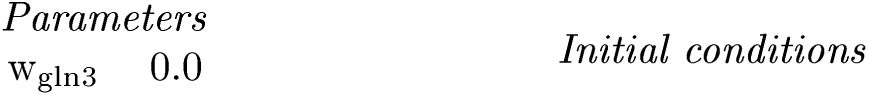

##### Model agrees with experiment

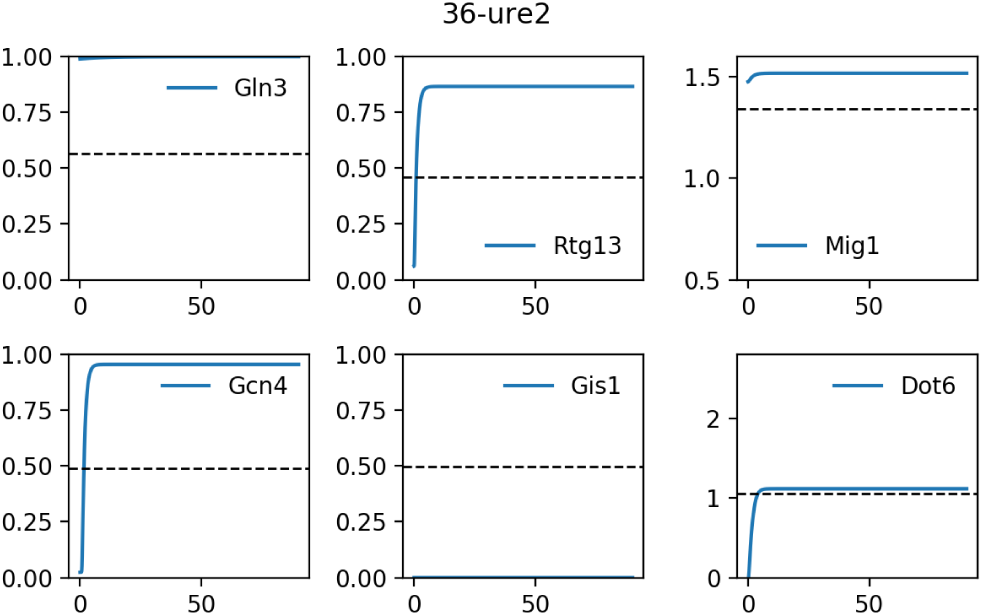

### S4.37 37-tap42-11

#### Readout used is Gln3 Gcn4

| TF | Interpreted | Simulated | Simulation |
| --- | --- | --- | --- |
| *Gis1* | - | - | 0.000 |
| *Mig1* | - | - | 1.503 |
| *Dot6* | - | - | 1.108 |
| **Gcn4** | Off | Off | 0.398 |
| *Rtg13* | - | - | 0.866 |
| **Gln3** | Off | On | 0.570 |

##### Description

Huber et al, 2009 studied a *tap42-11* strain (TB50) grown in YPD + rapamycin.

##### Representation

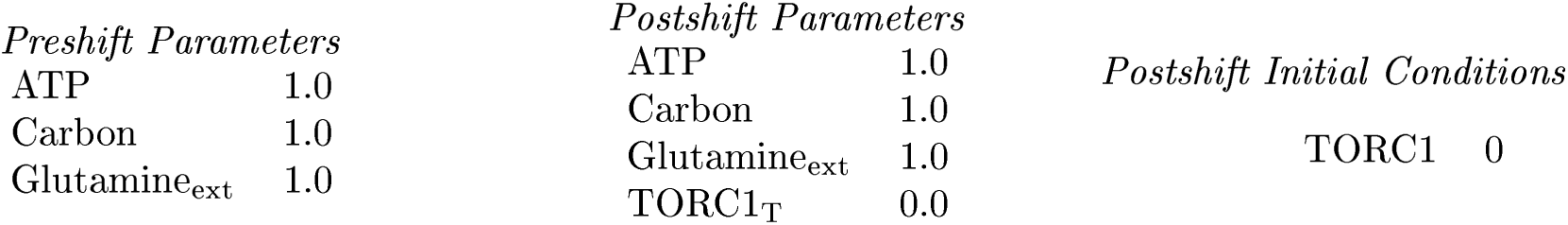

##### Mutant definition

| Parameters |  | Initial conditions |
| --- | --- | --- |
| $w_{\text{gcn}}$ | 0.0 | |
| $w_{\text{glnsit}}$ | 0.0 | |

##### Model does not agree with experiment

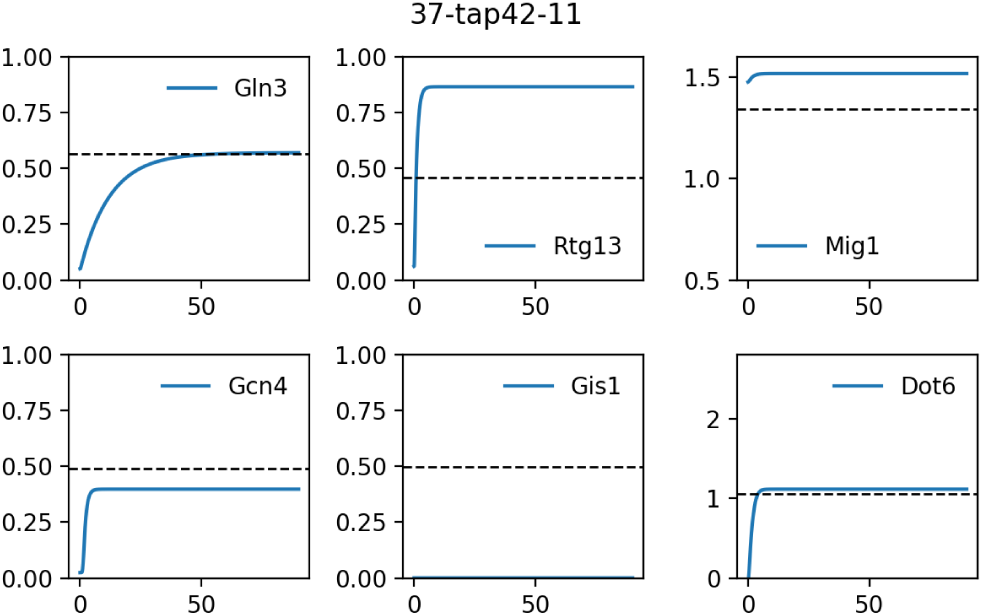

### S4.38 38-SCH9^**DE**^

#### Readout used is Gln3 Gcn4

| TF | Interpreted | Simulated | Simulation |
| --- | --- | --- | --- |
| *Gis1* | - | - | 0.000 |
| *Mig1* | - | - | 1.467 |
| *Dot6* | - | - | -0.103 |
| **Gcn4** | Off | Off | 0.024 |
| *Rtg13* | - | - | 0.866 |
| **Gln3** | Off | On | 1.000 |

##### Description

Huber et al, 2009 studied a *SCH9*^*DE*^ strain (TB50) grown in YPD + rapamycin.

##### Representation

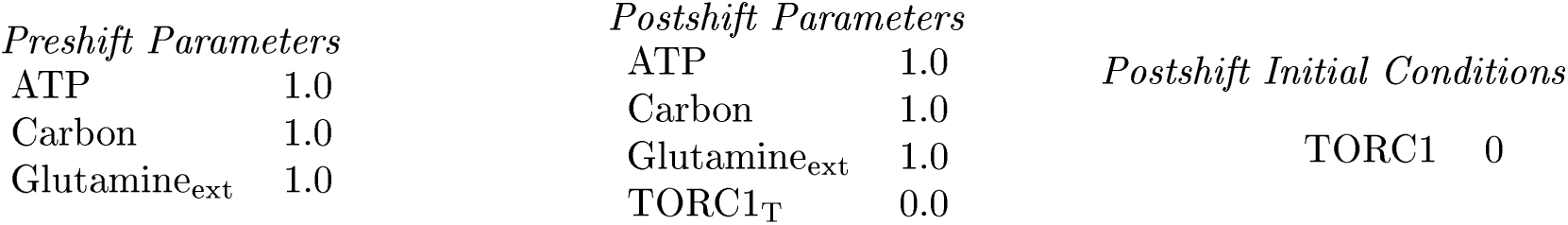

##### Mutant definition

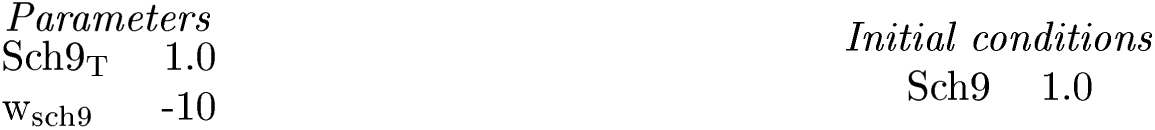

##### Model does not agree with experiment

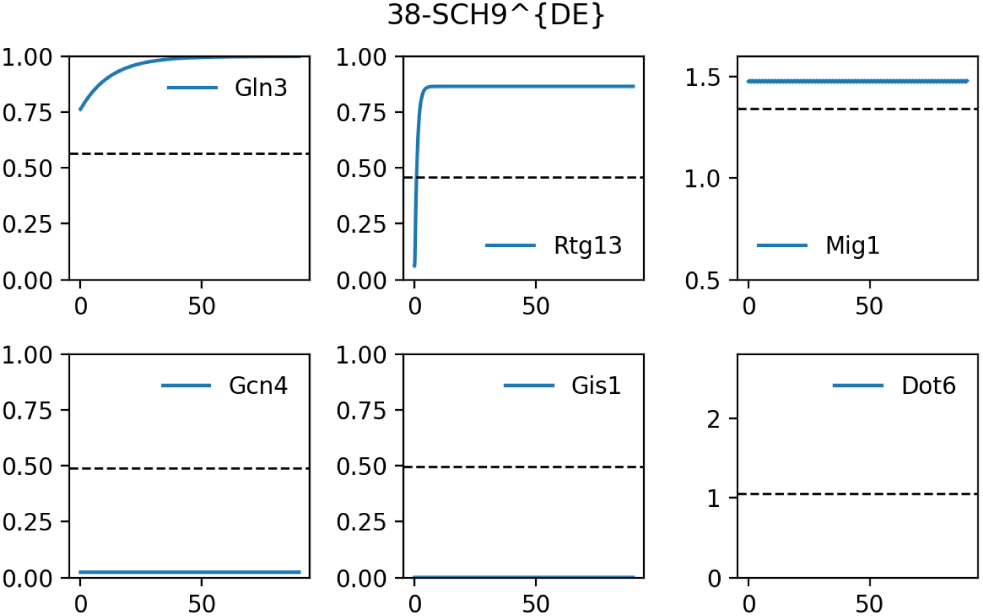

### S4.39 39-SCH9^DE^ tap42

| TF | Interpreted | Simulated | Simulation |
| --- | --- | --- | --- |
| *Gis1* | - | - | 0.000 |
| *Mig1* | - | - | 1.467 |
| *Dot6* | - | - | -0.103 |
| **Gcn4** | Off | Off | 0.024 |
| *Rtg13* | - | - | 0.866 |
| **Gln3** | Off | On | 0.596 |

#### Description

Huber et al, 2009 studied a *SCH9*^*DE*^ *tap42* strain (TB50) grown in YPD + rapamycin.

#### Representation

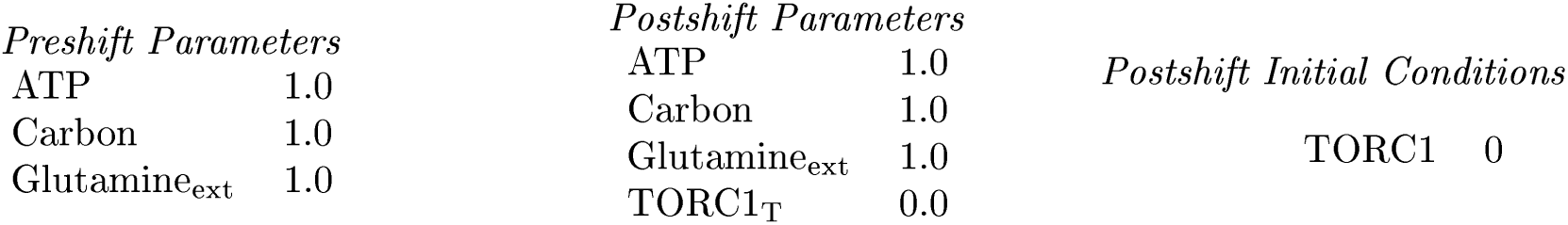

#### Mutant definition

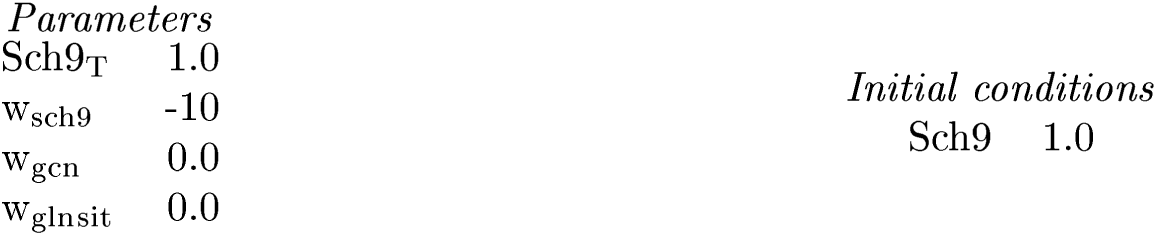

#### Model does not agree with experiment

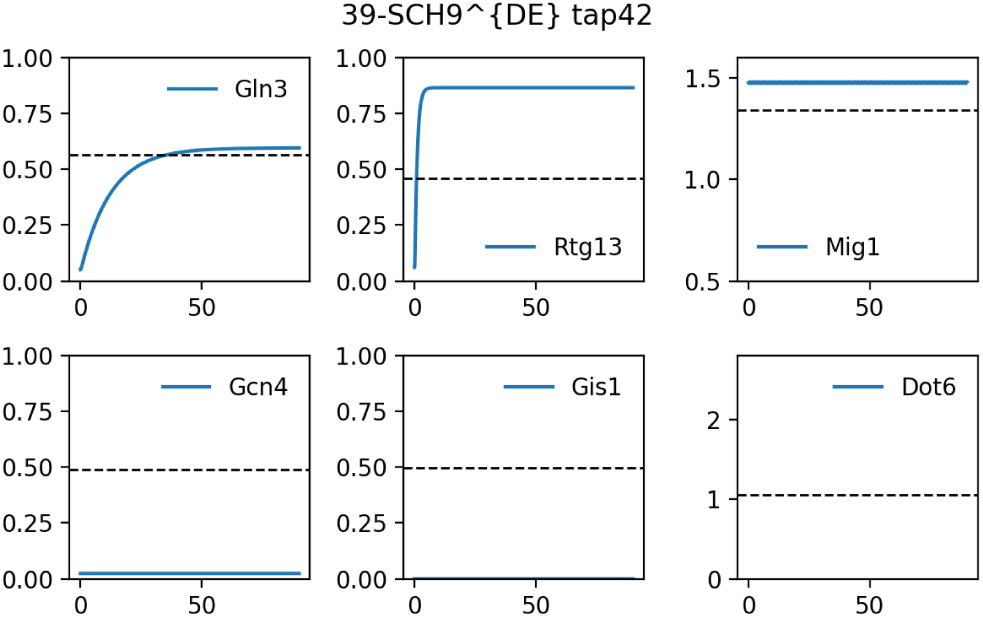

### S4.40 40-sch9

| TF | Interpreted | Simulated | Simulation |
| --- | --- | --- | --- |
| **Gis1** | On | Off | 0.046 |
| *Mig1* | - | - | 1.264 |
| *Dot6* | - | - | 1.825 |
| *Gcn4* | - | - | 0.963 |
| *Rtg13* | - | - | 0.577 |
| *Gln3* | - | - | 1.000 |

#### Description

Roosen et al, 2005 studied a *sch9* strain (W303-1) grown in YP + glycerol.

#### Representation

**Representation:**
| *Preshift Parameters* |  | *Postshift Parameters* |  | *Postshift Initial Conditions* |
| --- | --- | --- | --- | --- |
| ATP | 1.0 | ATP | 0.01 |  |
| Carbon | 1.0 | Carbon | 0.01 |  |
| Glutamine<sub>ext</sub> | 1.0 | Glutamine<sub>ext</sub> | 1.0 |  |

#### Mutant definition

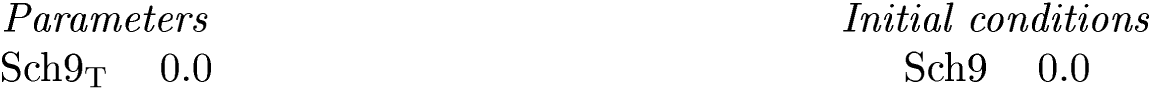

| *Parameters* | *Initial conditions* |
| --- | --- |
| Sch9<sub>T</sub> 0.0 | Sch9 0.0 |

#### Model does not agree with experiment

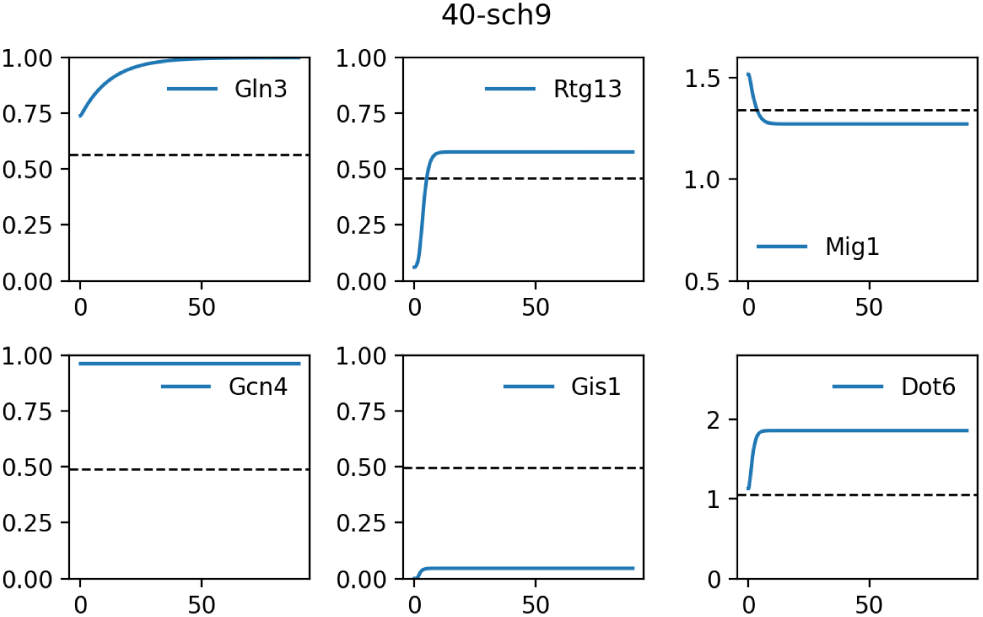

### S4.41 41-sch9 gis1

| TF | Interpreted | Simulated | Simulation |
| --- | --- | --- | --- |
| **Gis1** | Off | Off | 0.000 |
| *Mig1* | - | - | 1.264 |
| *Dot6* | - | - | 1.825 |
| *Gcn4* | - | - | 0.963 |
| *Rtg13* | - | - | 0.577 |
| *Gln3* | - | - | 1.000 |

#### Description

Roosen et al, 2005 studied a *sch9 gis1* strain (W303-1) grown in YP + glycerol.

#### Representation

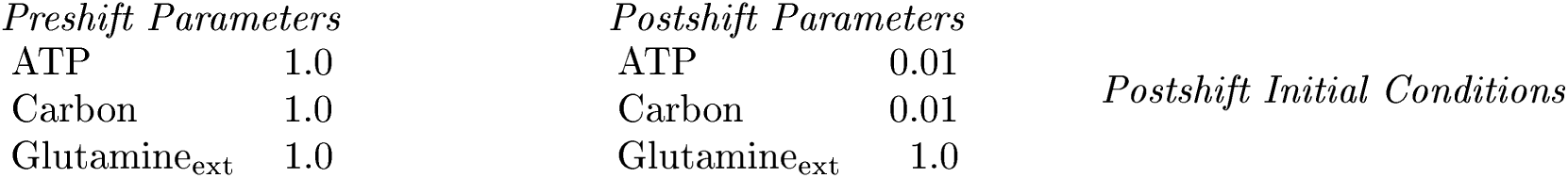

#### Mutant definition

| *Parameters* | *Initial conditions* |
| --- | --- |
| Gis1<sub>T</sub> 0.0 | Gis1 0.0 |
| Sch9<sub>T</sub> 0.0 | Sch9 0.0 |

#### Model agrees with experiment

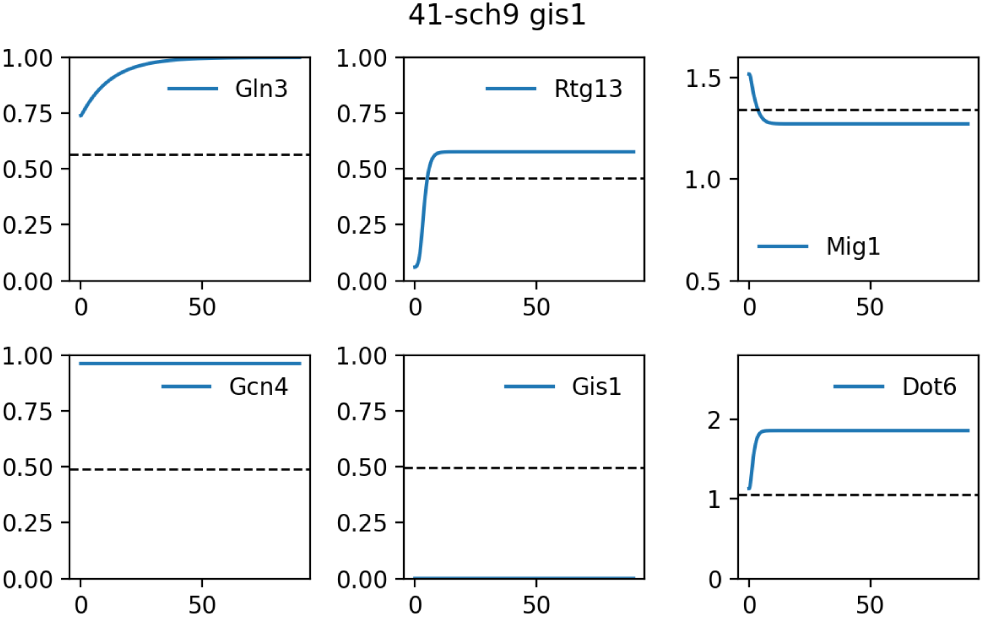

## Notes

https://github.com/amoghpj/nutrient-signaling

## References

[1] M. Conrad, J. Schothorst, H. N. Kankipati, G. Van Zeebroeck, M. Rubio-Texeira, and J. M. Thevelein. Nutrient sensing and signaling in the yeast Saccharomyces cerevisiae. FEMS microbiology reviews, 38(2):254–299, 2014.

[2] M. Schaechter, O. MaalØe, and N. O. Kjeldgaard. Dependency on medium and temperature of cell size and chemical composition during balanced growth of Salmonella Typhimurium. Journal of General Microbiology, 19(3):592–606, 1958.

[3] E. Metzl-Raz, M. Kafri, G. Yaakov, I. Soifer, Y. Gurvich, and N. Barkai. Principles of cellular resource allocation revealed by condition-dependent proteome profiling. eLife Sciences., 6:e28034, Aug 2017.

[4] C. D. Virgilio and R. Loewith. The Tor signalling network from yeast to man. The International Journal of Biochemistry & Cell Biology, 38(9):1476–1481, 2006.

[5] L. Chantranupong, R. L. Wolfson, and D. M. Sabatini. Nutrient-sensing mechanisms across evolution. Cell, 161(1):67–83, 2015.

[6] K. J. Verstrepen, D. Iserentant, P. Malcorps, G. Derdelinckx, P. V. Dijck, J. Winderickx, I. S. Pretorius, J. M. Thevelein, and F. R. Delvaux. Glucose and sucrose: Hazardous fast-food for industrial yeast? Trends in Biotechnology, 22(10):531–537, 2004.

[7] K. Mbonyi, M. Beullens, K. Detremerie, L. Geerts, and J. M. Thevelein. Requirement of one functional Ras gene and inability of an oncogenic Ras variant to mediate the glucose-induced cyclic AMP signal in the yeast Saccharomyces cerevisiae. Molecular and Cellular Biology, 8(8):3051–3057, 1988.

[8] K. Mbonyi, L. van Aelst, J. C. Argüelles, A. W. Jans, and J. M. Thevelein. Glucose-induced hyperaccumulation of cyclic amp and defective glucose repression in yeast strains with reduced activity of cyclic amp-dependent protein kinase. Molecular and Cellular Biology, 10(9):4518–4523, 1990.

[9] S. Zaman, S. I. Lippman, L. Schneper, N. Slonim, and J. R. Broach. Glucose regulates transcription in yeast through a network of signaling pathways. Molecular Systems Biology, 5(nil):nil, 2009.

[10] V. A. Cherkasova, H. Qiu, and A. G. Hinnebusch. Snf1 promotes phosphorylation of the subunit of eukaryotic translation initiation factor 2 by activating gcn2 and inhibiting phosphatases glc7 and sit4. Molecular and Cellular Biology, 30(12):2862–2873, 2010.

[11] A. Schöler and H. J. Schüller. A carbon source-responsive promoter element necessary for activation of the isocitrate lyase gene icl1 is common to genes of the gluconeogenic pathway in the yeast Saccharomyces cerevisiae. Molecular and Cellular Biology, 14(6):3613–3622, 1994.

[12] B. Magasanik and C. A. Kaiser. Nitrogen regulation in Saccharomyces cerevisiae. Gene, 290(1):1–18, 2002.

[13] D. E. Martin, T. Powers, and M. N. Hall. Regulation of ribosome biogenesis: Where is Tor? Cell Metabolism, 4(4):259–260, 2006.

[14] T. Powers and P. Walter. Regulation of ribosome biogenesis by the rapamycin-sensitive Tor-signaling pathway in Saccharomyces Cerevisiae. Molecular Biology of the Cell, 10(4):987–1000, 1999.

[15] C. Cipollina, J. van den Brink, P. Daran-Lapujade, J. T. Pronk, D. Porro, and J. H. de Winde. Saccharomyces Cerevisiae sfp1: At the crossroads of central metabolism and ribosome biogenesis. Microbiology, 154(6):1686–1699, 2008.

[16] C. J. D. Como and K. T. Arndt. Nutrients, via the Tor proteins, stimulate the association of Tap42 with type 2a phosphatases. Genes & Development, 10(15):1904–1916, 1996.

[17] P. Jorgensen. A dynamic transcriptional network communicates growth potential to ribosome synthesis and critical cell size. Genes & Development, 18(20):2491–2505, 2004.

[18] I. Mayordomo, F. Estruch, and P. Sanz. Convergence of the Target of Rapamycin and the Snf1 protein kinase pathways in the regulation of the subcellular localization of Msn2, a transcriptional activator of STRE (Stress Response Element)-regulated genes. Journal of Biological Chemistry, 277(38):35650–35656, 2002.

[19] J. Roosen, K. Engelen, K. Marchal, J. Mathys, G. Griffioen, E. Cameroni, J. M. Thevelein, C. De Virgilio, B. De Moor, and J. Winderickx. PKA and Sch9 control a molecular switch important for the proper adaptation to nutrient availability. Molecular microbiology, 55(3):862–880, 2005.

[20] T. Yorimitsu, S. Zaman, J. R. Broach, and D. J. Klionsky. Protein Kinase A and Sch9 cooperatively regulate induction of autophagy in Saccharomyces Cerevisiae. Molecular Biology of the Cell, 18(10):4180–4189, 2007.

[21] M.-A. Deprez, E. Eskes, T. Wilms, P. Ludovico, and J. Winderickx. Ph homeostasis links the nutrient sensing PKA/TORC/Sch9 ménage-à-trois to stress tolerance and longevity. Microbial Cell, 5(3):119–136, 2018.

[22] J. E. H. Hallett, X. Luo, and A. P. Capaldi. State transitions in the TORC1 signaling pathway and information processing in Saccharomyces cerevisiae. Genetics, 198(2):773–786, 2014.

[23] P. Kraikivski, K. C. Chen, T. Laomettachit, T. Murali, and J. J. Tyson. From START to FINISH: computational analysis of cell cycle control in budding yeast. NPJ Systems Biology and Applications, 1:15016, 2015.

[24] K. A. Kim, S. L. Spencer, J. G. Albeck, J. M. Burke, P. K. Sorger, S. Gaudet, and D. H. Kim. Systematic calibration of a cell signaling network model. BMC Bioinformatics, 11(1):202, 2010.

[25] J. Gunawardena. Models in biology: ‘accurate descriptions of our pathetic thinking’. BMC Biology, 12(1):29, 2014.

[26] A. Wiemken and M. Durr. Characterization of amino acid pools in the vacuolar compartment of Saccharomyces cerevisiae. Archives of Microbiology, 101(1):45–57, 1974.

[27] H. Halvorson. A study of the properties of the free amino acid pool and enzyme synthesis in yeast. The Journal of General Physiology, 38(4):549–573, 1955.

[28] F. Messenguy, D. Colin, and J.-P. Ten Have. Regulation of compartmentation of amino acid pools in Saccharomyces cerevisiae and its effects on metabolic control. European Journal of Biochemistry, 108(2):439–447, 1980.

[29] S. C. Li and P. M. Kane. The yeast lysosome-like vacuole: Endpoint and Crossroads. Biochimica et Biophysica Acta (BBA) - Molecular Cell Research, 1793(4):650–663, 2009.

[30] D. Stracka, S. Jozefczuk, F. Rudroff, U. Sauer, and M. N. Hall. Nitrogen source activates Tor (target of rapamycin) complex 1 via glutamine and independently of Gtr/Rag proteins. Journal of Biological Chemistry, 289(36):25010–25020, 2014.

[31] M. Boeckstaens, E. Llinares, P. V. Vooren, and A. M. Marini. The TORC1 effector kinase Npr1 fine tunes the inherent activity of the Mep2 ammonium transport protein. Nature Communications, 5(1):3101, 2014.

[32] C. Garmendia-Torres, A. Goldbeter, and M. Jacquet. Nucleocytoplasmic oscillations of the yeast transcription factor Msn2: Evidence for periodic PKA activation. Current Biology, 17(12):1044–1049, 2007.

[33] D. Gonze, M. Jacquet, and A. Goldbeter. Stochastic modelling of nucleocytoplasmic oscillations of the transcription factor Msn2 in yeast. Journal of The Royal Society Interface, 5(Suppl 1):S95–S109, 2008.

[34] P. Cazzaniga, D. Pescini, D. Besozzi, G. Mauri, S. Colombo, and E. Martegani. Modeling and stochastic simulation of the Ras/cAMP/PKA pathway in the yeast Saccharomyces cerevisiae evidences a key regulatory function for intracellular guanine nucleotides pools. Journal of Biotechnology, 133(3):377–385, 2008.

[35] D. Pescini, P. Cazzaniga, D. Besozzi, G. Mauri, L. Amigoni, S. Colombo, and E. Martegani. Simulation of the Ras/cAMP/PKA pathway in budding yeast highlights the establishment of stable oscillatory states. Biotechnology Advances, 30(1):99–107, 2012.

[36] J. Stewart-Ornstein, S. Chen, R. Bhatnagar, J. S. Weissman, and H. El-Samad. Model-guided optogenetic study of PKA signaling in budding yeast. Molecular Biology of the Cell, 28(1):221–227, 2017.

[37] T. Williamson, J.-M. Schwartz, D. B. Kell, and L. Stateva. Deterministic mathematical models of the camp pathway in Saccharomyces Cerevisiae. BMC Systems Biology, 3(1):70, 2009.

[38] K. Gonzales, Ö. Kayikçi, D. G. Schaeffer, and P. M. Magwene. Modeling mutant phenotypes and oscillatory dynamics in the Saccharomyces cerevisiae cAMP-PKA pathway. BMC Systems Biology, 7(1):40, 2013.

[39] R. García-Salcedo, T. Lubitz, G. Beltran, K. Elbing, Y. Tian, S. Frey, O. Wolkenhauer, M. Krantz, E. Klipp, and S. Hohmann. Glucose de-repression by yeast AMP-activated protein kinase Snf1 is controlled via at least two independent steps. FEBS Journal, 281(7):1901–1917, 2014.

[40] N. Welkenhuysen, J. Borgqvist, M. Backman, L. Bendrioua, M. Goksör, C. B. Adiels, M. Cvijovic, and S. Hohmann. Single-cell study links metabolism with nutrient signaling and reveals sources of variability. BMC Systems Biology, 11(1):59, 2017.

[41] P. K. U. Vinod and K. V. Venkatesh. Quantification of the effect of amino acids on an integrated mtor and insulin signaling pathway. Molecular BioSystems, 5(10):1163, 2009.

[42] A. G. Sonntag, P. D. Pezze, D. P. Shanley, and K. Thedieck. A modelling-experimental approach reveals Insulin Receptor Substrate (IRS)-dependent Regulation of Adenosine Monosphosphate-Dependent Kinase (AMPK) By Insulin. FEBS Journal, 279(18):3314–3328, 2012.

[43] P. D. Pezze, S. Ruf, A. G. Sonntag, M. Langelaar-Makkinje, P. Hall, A. M. Heberle, P. R. Navas, K. van Eunen, R. C. Tölle, J. J. Schwarz, H. Wiese, B. Warscheid, J. Deitersen, B. Stork, E. Fäßler, S. Schäuble, U. Hahn, P. Horvatovich, D. P. Shanley, and K. Thedieck. A systems study reveals concurrent activation of AMPK and mTOR by amino acids. Nature Communications, 7(1):13254, 2016.

[44] K. Thobe, C. Sers, and H. Siebert. Unraveling the regulation of mTORC2 using logical modeling. Cell Communication and Signaling, 15(1):6, 2017.

[45] N. Sengupta, P. Vinod, and K. Venkatesh. Crosstalk between cAMP-PKA and MAP kinase pathways is a key regulatory design necessary to regulate Flo11 expression. Biophysical Chemistry, 125(1):59–71, 2007.

[46] N. Welkenhuysen, B. Schnitzer, L. Österberg, and M. Cvijovic. Robustness of nutrient signaling is maintained by interconnectivity between signal transduction pathways. Frontiers in Physiology, 9(nil):nil, 2019.

[47] S. Colombo, D. Ronchetti, J. M. Thevelein, J. Winderickx, and E. Martegani. Activation state of the Ras2 protein and glucose-induced signaling in Saccharomyces cerevisiae. Journal of Biological Chemistry, 279(45):46715–46722, 2004.

[48] D. Jian, Z. Aili, B. Xiaojia, Z. Huansheng, and H. Yun. Feedback Regulation of Ras2 Guanine Nucleotide Exchange Factor (Ras2-GEF) Activity of Cdc25p By Cdc25p Phosphorylation in the Yeast Saccharomyces cerevisiae. FEBS Letters, 584(23):4745–4750, 2010.

[49] P. Ma, S. Wera, P. V. Dijck, and J. M. Thevelein. The pde1-encoded low-affinity phosphodiesterase in the yeast Saccharomyces cerevisiae has a specific function in controlling agonist-induced cAMP signaling. Molecular Biology of the Cell, 10(1):91–104, 1999.

[50] S. Colombo. Involvement of distinct g-proteins, Gpa2 and Ras, in glucose-and intracellular acidification-induced cAMP signalling in the yeast Saccharomyces cerevisiae. The EMBO Journal, 17(12):3326–3341, 1998.

[51] A. Zhang and W. Gao. Mechanisms of protein kinase Sch9 regulating Bcy1 in Saccharomyces cerevisiae. FEMS microbiology letters, 331(1):10–16, 2012.

[52] R. Hatakeyama and C. D. Virgilio. Unsolved mysteries of Rag GTPase signaling in yeast. Small GTPases, 7(4):239–246, 2016.

[53] J. Urban, A. Soulard, A. Huber, S. Lippman, D. Mukhopadhyay, O. Deloche, V. Wanke, D. Anrather, G. Ammerer, H. Riezman, et al. Sch9 is a major target of TORC1 in Saccharomyces cerevisiae. Molecular cell, 26(5):663–674, 2007.

[54] R. Nicastro, F. Tripodi, M. Gaggini, A. Castoldi, V. Reghellin, S. Nonnis, G. Tedeschi, and P. Coccetti. Snf1 phosphorylates Adenylate Cyclase and negatively regulates Protein Kinase A-dependent transcription in Saccharomyces cerevisiae. Journal of Biological Chemistry, 290(41):24715–24726, 2015.

[55] R. Nicastro, F. Tripodi, C. Guzzi, V. Reghellin, S. Khoomrung, C. Capusoni, C. Compagno, C. Airoldi, J. Nielsen, L. Alberghina, and P. Coccetti. Enhanced amino acid utilization sustains growth of cells lacking Snf1/AMPK. Biochimica et Biophysica Acta (BBA) - Molecular Cell Research, 1853(7):1615–1625, 2015.

[56] I. Pedruzzi, N. Burckert, P. Egger, and C. D. Virgilio. Saccharomyces Cerevisiae Ras/cAMP pathway controls Post-Diauxic Shift Element-dependent transcription through the zinc finger protein gis1. The EMBO Journal, 19(11):2569–2579, 2000.

[57] D. Castermans, I. Somers, J. Kriel, W. Louwet, S. Wera, M. Versele, V. Janssens, and J. M. Thevelein. Glucose-induced posttranslational activation of protein phosphatases PP2a and PP1 in yeast. Cell Research, 22(6):1058–1077, 2012.

[58] P. G. Bertram, J. H. Choi, J. Carvalho, T.-F. Chan, W. Ai, and X. F. S. Zheng. Convergence of Tornitrogen and Snf1-glucose signaling pathways onto Gln3. Molecular and Cellular Biology, 22(4):1246–1252, 2002.

[59] J. L. Crespo, T. Powers, B. Fowler, and M. N. Hall. The Tor-controlled transcription activators Gln3, Rtg1, and Rtg3 are regulated in response to intracellular levels of glutamine. Proceedings of the National Academy of Sciences, 99(10):6784–6789, 2002.

[60] V. A. Cherkasova and A. G. Hinnebusch. Translational control by Tor and Tap42 through dephosphorylation of eIF2*α* kinase Gcn2. Genes & Development, 17(7):859–872, 2003.

[61] A. G. Hinnebusch. Translational regulation of GCN4 and the general amino acid control of yeast. Annu. Rev. Microbiol., 59:407–450, 2005.

[62] C. D. Virgilio and R. Loewith. Cell growth control: Little eukaryotes make big contributions. Oncogene, 25(48):6392–6415, 2006.

[63] K. Hirimburegama, P. Durnez, J. Keleman, E. Oris, R. Vergauwen, H. Mergelsberg, and J. M. Thevelein. Nutrient-induced activation of trehalase in nutrient-starved cells of the yeast Saccharomyces cerevisiae: cAMP is not involved as second messenger. Journal of General Microbiology, 138(10):2035–2043, 1992.

[64] J. Chaillot, F. Tebbji, J. Malick, and A. Sellam. Dot6 Is a Major Regulator of Cell Size and a Transcriptional Activator of Ribosome Biogenesis in the Opportunistic Yeast Candida albicans. Genetics, page genetics.301872.2018, Dec 2018.

[65] V. C. Özalp, T. R. Pedersen, L. J. Nielsen, and L. F. Olsen. Time-Resolved Measurements of Intracellular ATP in the Yeast Saccharomyces cerevisiae using a New Type of Nanobiosensor. Journal of Biological Chemistry, 285(48):37579–37588, 2010.

[66] M.-P. Péli-Gulli, A. Sardu, N. Panchaud, S. Raucci, and C. D. Virgilio. Amino acids stimulate TORC1 through Lst4-lst7, a GTPase-activating protein complex for the Rag family GTPase Gtr2. Cell Reports, 13(1):1–7, 2015.

[67] N. Panchaud, M.-P. Peli-Gulli, and C. D. Virgilio. Amino acid deprivation inhibits TORC1 through a GTPase-activating protein complex for the Rag family GTPase Gtr1. Science Signaling, 6(277):ra42–ra42, 2013.

[68] K. Matsumoto, I. Uno, Y. Oshima, and T. Ishikawa. Isolation and characterization of yeast mutants deficient in Adenylate Cyclase and cAMP-dependent protein kinase. Proceedings of the National Academy of Sciences, 79(7):2355–2359, 1982.

[69] J. M. Thevelein. Fermentable sugars and intracellular acidification as specific activators of the Ras-Adenylate Cyclase signalling pathway in yeast: the relationship to nutrient-induced cell cycle control. Molecular Microbiology, 5(6):1301–1307, 1991.

[70] L. Barrett, M. Orlova, M. Maziarz, and S. Kuchin. Protein Kinase A contributes to the negative control of Snf1 protein kinase in Saccharomyces cerevisiae. Eukaryotic Cell, 11(2):119–128, 2011.

[71] J. E. H. Hallett, X. Luo, and A. P. Capaldi. Snf1/AMPK promotes the formation of Kog1/Raptor-bodies to increase the activation threshold of TORC1 in budding yeast. eLife, 4(nil):nil, 2015.

[72] M. Prouteau, A. Desfosses, C. Sieben, C. Bourgoint, N. L. Mozaffari, D. Demurtas, A. K. Mitra, P. Guichard, S. Manley, and R. Loewith. TORC1 organized in inhibited domains (TOROIDs) regulate TORC1 activity. Nature, 550(7675):265–269, 2017.

[73] M. Crauwels, M. C. V. Donaton, M. B. Pernambuco, J. Winderickx, J. H. de Winde, and J. M. Thevelein. The Sch9 protein kinase in the yeast Saccharomyces cerevisiae controls cAPK activity and is required for nitrogen activation of the fermentable-growth-medium-induced (FGM) pathway. Microbiology, 143(8):2627–2637, 1997.

[74] A. Zhang, Y. Shen, W. Gao, and J. Dong. Role of Sch9 in regulating Ras-cAMP signal pathway in Saccharomyces cerevisiae. FEBS letters, 585(19):3026–3032, 2011.

[75] W. A. Wilson, S. A. Hawley, and D. Hardie. Glucose repression/derepression in budding yeast: Snf1 protein kinase is activated by phosphorylation under derepressing conditions, and this correlates with a high AMP:ATP ratio. Current Biology, 6(11):1426–1434, 1996.

[76] T. Beck and M. N. Hall. The Tor signalling pathway controls nuclear localization of nutrient-regulated transcription factors. Nature, 402(6762):689–692, 1999.

[77] S.-P. Hong, F. C. Leiper, A. Woods, D. Carling, and M. Carlson. Activation of yeast Snf1 and mammalian AMP-activated protein kinase by upstream kinases. Proceedings of the National Academy of Sciences, 100(15):8839–8843, 2003.

[78] R. N. Gutenkunst, J. Waterfall, F. Casey, K. Brown, C. R. Myers, and J. P. Sethna. Universally sloppy parameter sensitivities in systems biology models. PLoS Computational Biology, preprint(2007):e189, 2005.

[79] I. Tavassoly, J. Parmar, A. Shajahan-Haq, R. Clarke, W. Baumann, and J. Tyson. Dynamic modelling of the interaction between autophagy and apoptosis in mammalian cells. CPT: Pharmacometrics & Systems Pharmacology, 4(4):263–272, 2015.

[80] L. Valenzuela, C. Aranda, and A. Gonzalez. Tor modulates Gcn4-dependent expression of genes turned on by nitrogen limitation. Journal of Bacteriology, 183(7):2331–2334, 2001.

[81] K. A. Staschke, S. Dey, J. M. Zaborske, L. R. Palam, J. N. McClintick, T. Pan, H. J. Edenberg, and R. C. Wek. Integration of General Amino Acid Control and Target of Rapamycin (Tor) regulatory pathways in nitrogen assimilation in yeast. Journal of Biological Chemistry, 285(22):16893–16911, 2010.

[82] Z. Liu and R. A. Butow. A transcriptional switch in the expression of yeast tricarboxylic acid cycle genes in response to a reduction or loss of respiratory function. Molecular and Cellular Biology, 19(10):6720–6728, 1999.

[83] Z. Liu, T. Sekito, M. Špı’rek, J. Thornton, and R. A. Butow. Retrograde signaling is regulated by the dynamic interaction between Rtg2p and Mks1p. Molecular cell, 12(2):401–411, 2003.

[84] H. R. D. Balciunas. Yeast genes Gis1-4: Multicopy suppressors of the Gal - phenotype of Snf1 Mig1 Srb8/10/11 cells. Molecular and General Genetics MGG, 262(4-5):589–599, 1999.

[85] N. Gasmi, P.-E. Jacques, N. Klimova, X. Guo, A. Ricciardi, F. Robert, and B. Turcotte. The switch from fermentation to respiration in Saccharomyces cerevisiae is regulated by the Ert1 transcriptional activator/repressor. Genetics, 198(2):547–560, 2014.

[86] R. A. Butow and N. G. Avadhani. Mitochondrial signaling: the retrograde response. Molecular cell, 14(1):1–15, 2004.

[87] M. J. Brauer, C. Huttenhower, E. M. Airoldi, R. Rosenstein, J. C. Matese, D. Gresham, V. M. Boer, O. G. Troyanskaya, and D. Botstein. Coordination of growth rate, cell cycle, stress response, and metabolic activity in yeast. Molecular Biology of the Cell, 19(1):352–367, 2007.

[88] A. A. Granados, J. M. J. Pietsch, S. A. Cepeda-Humerez, I. L. Farquhar, G. Tkačik, and P. S. Swain. Distributed and dynamic intracellular organization of extracellular information. Proc. Natl. Acad. Sci. U.S.A., 115(23):6088–6093, Jun 2018.

[89] J. C. Ewald. How yeast coordinates metabolism, growth and division. Current Opinion in Microbiology, 45(nil):1–7, 2018.

[90] T. Laomettachit, K. C. Chen, W. T. Baumann, and J. J. Tyson. A model of yeast cell-cycle regulation based on a standard component modeling strategy for protein regulatory networks. PLOS ONE, 11(5):e0153738, 2016.

[91] J. J. Tyson, T. Laomettachit, and P. Kraikivski. Modeling the dynamic behavior of biochemical regulatory networks. Journal of Theoretical Biology, 462(nil):514–527, 2019.

[92] A. Rohatgi. Webplotdigitizer, 2017.

[93] C. A. Schneider, W. S. Rasband, and K. W. Eliceiri. NIH Image to ImageJ: 25 years of image analysis. Nature methods, 9(7):671–675, 2012.

[94] J. R. Magnus and H. Neudecker. The commutation matrix: Some properties and applications. The Annals of Statistics, 7(2):381–394, 1979.

[95] K.-C. Li. On principal hessian directions for data visualization and dimension reduction: Another application of stein’s lemma. Journal of the American Statistical Association, 87(420):1025–1039, 1992.

[96] N. C. Barbet, U. Schneider, S. B. Helliwell, I. Stansfield, M. F. Tuite, and M. N. Hall. Tor controls translation initiation and early g1 progression in yeast. Molecular Biology of the Cell, 7(1):25–42, 1996.

[97] A. Huber, B. Bodenmiller, A. Uotila, M. Stahl, S. Wanka, B. Gerrits, R. Aebersold, and R. Loewith. Characterization of the rapamycin-sensitive phosphoproteome reveals that Sch9 is a central coordinator of protein synthesis. Genes & Development, 23(16):1929–1943, 2009.

